# The geography of metacommunities: landscape characteristics drive geographic variation in the assembly process through selecting species pool attributes

**DOI:** 10.1101/2023.06.20.545715

**Authors:** Gabriel Khattar, Pedro Peres-Neto

## Abstract

Metacommunity ecology traditionally disregards that the dominant life-histories observed in species pools are selected by the characteristics of landscapes where the assembly process takes place. Recognizing the importance of this relationship is relevant because it integrates macroecological principles into metacommunity theory, generating a greater understanding about the ecological causes underlying broad-scale geographic variation in the relative importance of assembly mechanisms. To demonstrate that, we employed simulation models in which species pools with the same initial distribution of niche breadths and dispersal abilities interacted in landscapes with contrasting characteristics. By assessing the traits of species that dominated the metacommunity in each landscape type, we determined how different landscape characteristics select for different life-history strategies at the metacommunity level. We also analyzed the simulated data to derive predictions about the causal links between landscape characteristics, dominant life-histories in species pools, and their mutual influence on empirical inferences about the assembly process. We provide empirical support to these predictions by contrasting the assembly process of moth metacommunities in a tropical versus a temperate mountainous landscape. Collectively, our simulation models and empirical analyses illustrate how our framework can be formalized as an inferential tool for investigating the geography of metacommunity assembly.

## Introduction

Metacommunity ecology integrates core ecological assembly mechanisms such as dispersal limitation, environmental selection, and ecological drift to better understand and explain a multitude of ecological patterns (Mouquet and Loreau 2003; Vellend et al. 2014; Fournier et al. 2017; Koffel et al. 2022). Theoretical models have advanced our knowledge about the importance and links among these mechanisms mostly by systematically manipulating parameters that govern two different metacommunity components: (1) the attributes of species pools that form metacommunities (e.g., degree of ecological specialization, niche differentiation, and dispersal ability), and; (2) the characteristics (e.g., environmental heterogeneity, connectivity) of landscapes where the metacommunity takes place. For instance, by manipulating species ecological specialization and dispersal abilities along with landscape environmental heterogeneity and connectivity, one can observe simulated metacommunity dynamics varying along a continuum from niche-deterministic to neutrally-stochastic assembly (e.g., Viana and Chase 2019, Thompson et al. 2020). By using simulation models that combine varying levels of species pools attributes and landscape characteristics, these studies can provide insights into how patterns observed in empirical metacommunities can be linked to different combinations of independent ecological mechanisms (e.g., Ovaskainen et al. 2019, Guzman et al. 2022).

However, by adopting cross-factorial designs wherein the parameters that set landscape characteristics and species pool attributes are independently manipulated, one makes the implicit assumption that these two metacommunity components are independent axes in the assembly process (e.g., Thompson et al. 2020). In other words, this modeling design implicitly assumes that the dominant life-history strategies observed in species pools forming metacommunities are not inherently influenced by the characteristics of landscapes where the assembly process takes place. While heuristic, this assumption renders metacommunity ecology blind to the well-established links between species pool attributes and landscape characteristics that underlie ecogeographical rules, macroecological hypotheses, and biogeographic patterns. For instance, Janzen’s seasonality hypothesis states that latitudinal variation in the degree of spatial and temporal variation in landscape environmental conditions explains latitudinal clines in the degree of ecological specialization of species in the regional pools (Janzen 1967; Ghalambor 2006; Sheldon et al. 2018). Rapoport’s rule (i.e., the increase in species geographic ranges with latitude, Stevens 1989, Ruggiero and Werenkraut 2007) is thought to be a consequence of the dominance of strong disperses in temperate landscapes where temporal variability in habitat conditions is high. Additionally, theoretical approaches have demonstrated that the spatial distribution of environmental conditions (from random to highly autocorrelated) and the degree of physical connectivity among patches select the overall level of dispersal ability of species in the regional pool because it determines the costs and risks of dispersal events that allow species to reach suitable patches (Büchi et al. 2009; Fournier et al. 2017). Collectively, these empirical and theoretical studies demonstrate that the influence of landscape attributes on the dominant life-history strategies observed in regional pools is common in nature and, therefore, should be integrated into the theoretical framework of metacommunity ecology.

In this study, we aim at demonstrating that a conceptual framework centered on the dependency of species pool characteristics on landscape attributes is valuable for understanding the geography of metacommunity assembly, i.e., understanding variation in the relative importance of mechanisms that assemble different metacommunities distributed along broad-scale ecological gradients or across biogeographic regions. Our proposed conceptual framework can be described as a partial mediation model (Figure 1) in which landscape attributes (i.e., independent variables) determine the degree of specialization and dispersal ability of species that dominate species pools at the metacommunity scale (i.e., mediator variables). Jointly, these two (model) compartments dictate the relative importance of different assembly mechanisms (i.e., exogenous variable). Putting in ecological terms: landscape attributes that vary across large-scale gradients (e.g., seasonality or spatial autocorrelation of environmental features) should determine geographic changes in the dominant traits and life-history strategies observed in species pools that form metacommunities (Peres-Neto et al. 2012, Henriques-Silva et al. 2015). These non-random associations between landscape characteristics and species pool attributes, as demonstrated in this study, underly predictable shifts in the relative importance of assembly mechanisms.

**Figure 1:**
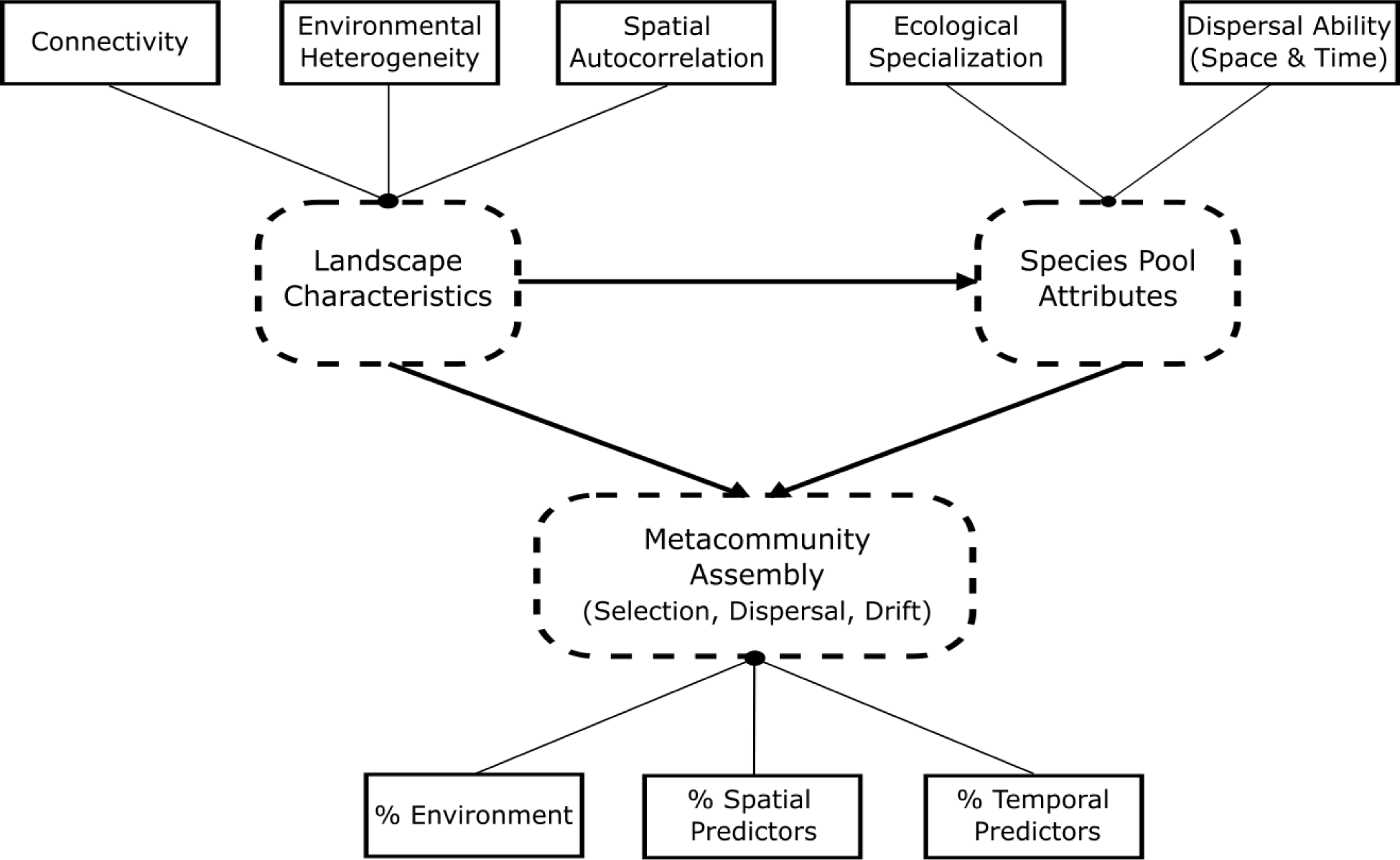
A mediation model for the geography of metacommunity assembly. It incorporates the effects of both landscape (exogenous variables) and species pool (mediator variables) attributes on the relative importance of selection, dispersal, drift (i.e., Endogenous variable). Dashed, round-edges boxes represent theoretical constructs, i.e., components of the metacommunity theory that are inferred from measurable variables and patterns observed in empirical metacommunities (solid rectangles). “%” represent the amount of variation in community composition explained by environmental variables, and spatial and temporal predictors. The variation explained by their covariation (i.e., joint contribution) is omitted.

To validate and illustrate the utility of our conceptual framework, we built a process-based metacommunity model wherein species pools with the same initial distribution of continuous traits (here, ecological specialization and dispersal ability) were allowed to colonize and reach coexistence in landscapes varying in their levels of seasonality, physical connectivity, and spatial structure (autocorrelation) of environmental (habitat) conditions. By assessing the degree of ecological specialization and dispersal ability of the species that could persist and dominate the metacommunity in each type of landscape, we were able to understand how different landscape characteristics select for different dominant life-history strategies observed in species pools. Additionally, our model allowed us to understand whether geographic clines on life-history strategies, commonly recognized as the outcome of broad-scale variation in evolutionary and historical mechanisms, can emerge as a result of ecological dynamics operating at the finer spatial and temporal scales of metacommunity dynamics (Henriques-Silva et al. 2015, Mittelbach and Schemske 2015).

Following, we analyzed the resulting (simulated) metacommunities using analytical approaches commonly used to study empirical metacommunities. This allowed us to understand how the interdependences of species pools and landscapes influence our empirical inferences about the relative importance of different assembly processes. We used variation partitioning to contrast the effects of environmental factors (typically assumed as the result of niche-based processes), spatial (typically assumed as the result of dispersal limitation), and temporal variation (typically assumed as the result of temporal autocorrelation in population dynamics generated by neutral mechanisms) in explaining variation in community composition (but see relevant conceptual and statistical limitations of this approach when applied to empirical data - Gilbert and Bennett 2010, Peres-Neto and Legendre 2010, Ovaskainen et al. 2019). Since we used a simulation model that incorporates known processes and lacks missing predictors (such as unmeasured spatiotemporal environmental variables that influence species distribution), we can use variation partitioning to draw direct inferences from the observed patterns, which may be challenging when using empirical data. By establishing the links between landscape attributes, species pool characteristics, and associated inferences about community assembly derived from statistical models, we were able to foster theoretical predictions about the variation in the relative importance of mechanisms observed in empirical metacommunities located at different parts of large-scale ecological gradients or in different biogeographic regions (e.g., Qian and Ricklefs 2012; Myers et al. 2013; Nishizawa et al. 2022).

To provide empirical support for some of the theoretical predictions derived from our conceptual framework, we analyzed data on moth metacommunities in a tropical and a temperate mountainous landscape that show distinct patterns of spatial and temporal environmental heterogeneity (Zuloaga and Kerr 2017). As such, we were able to contrast core predictions generated by our conceptual model against the outcomes observed in metacommunities that exhibit significant variation in both species pool characteristics and landscape attributes.

## Methods

We aimed at understanding how different landscape characteristics select for different levels of ecological specialization and dispersal ability in the species pools that form metacommunities. To do so, we modeled metacommunity dynamics in space and time in landscapes with varying levels of different attributes. We then recorded the overall niche breadth and dispersal ability of species that were able to persist and dominate the resulting simulated metacommunities in each type of landscape. For the sake of brevity, we only briefly describe how we simulated landscapes and metacommunity dynamics here. An extended description and R code to recreate our simulation setup and further analyses are found in Supp. Material I.

### Simulated landscapes

We generated a total of 216 types of landscapes considering a wide range of spatiotemporal heterogeneity levels (8), physical connectivity (9 levels), and spatial distribution of environmental conditions (3 levels) (Figure 2). These landscape attributes have been shown to modulate the mechanisms underlying species coexistence which, in turn, influence metacommunity dynamics (Büchi et al. 2009; Moritz et al. 2013; Fournier et al. 2017).

**Figure 2.**
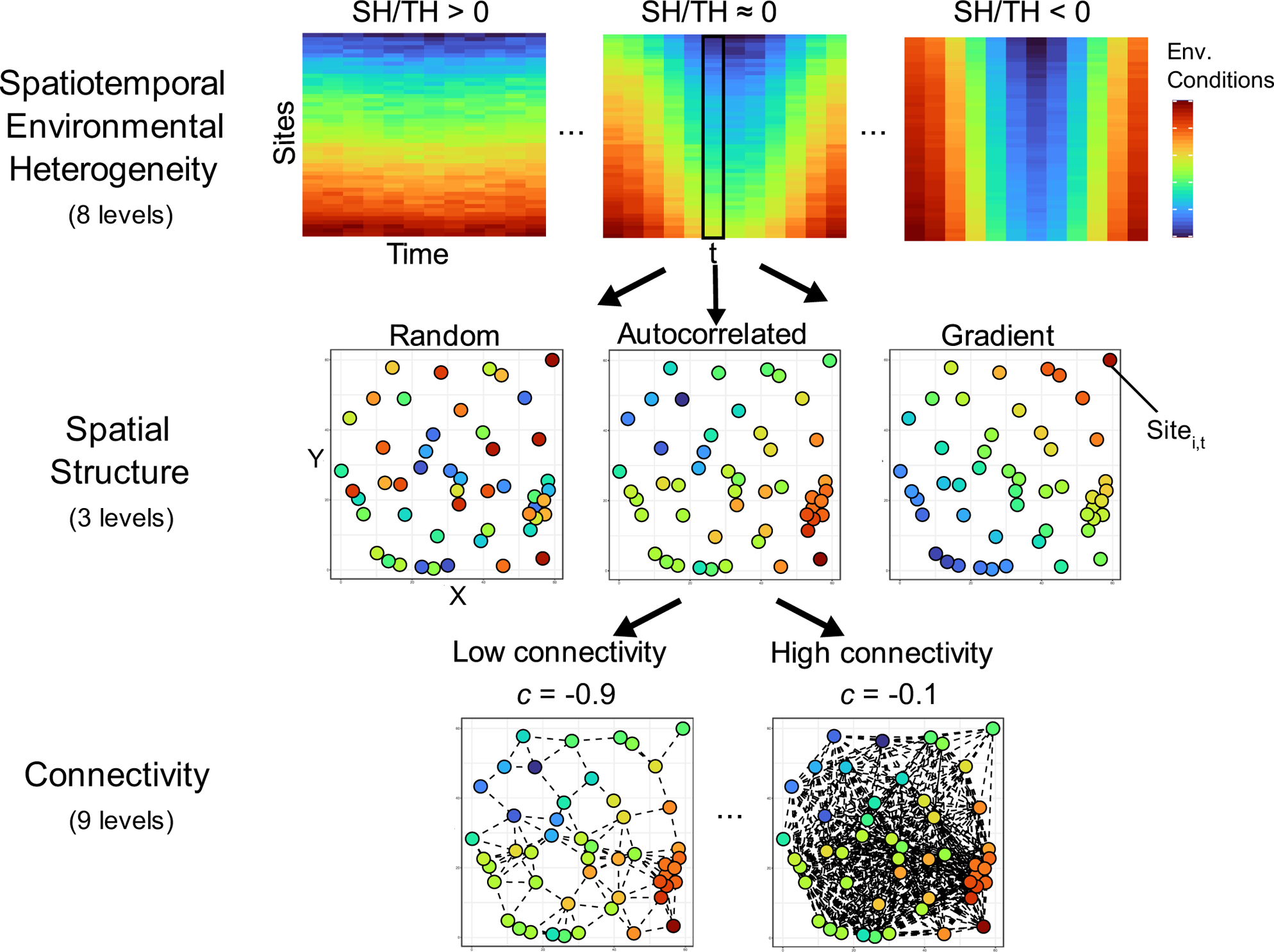
Schematic representation of simulated landscape characteristics. Spatiotemporal environmental heterogeneity SH/TH is calculated as the log of the ratio between the average variance of environmental conditions in space (SH) and the average variance of environmental conditions in time (TH). Spatial structure represents the type of spatial distribution of environmental conditions considered in the simulations - from totally random, through autocorrelated landscape, to a linear gradient. Connectivity decayed exponentially with distance at rate *c* and values below a fixed threshold were truncated to 0.

We started by randomly distributing 60 patches in a geographic space defined by x and y coordinates ranging from 0 to 60. The degree of physical connectivity between pairs of patches was set as a negative exponential function of their distance. Pairwise connectivity defines the weighted probabilities of spatial dispersal between pairs of patches (see *Species pools and metacommunity dynamics* and Supp. Material I). Connectivity values below a threshold of 10^-4^ were truncated to 0 to generate truly disconnected pairs of patches (as in Fournier et al. 2017). By varying the degree of exponential decay in connectivity but keeping this threshold constant, we generated landscapes with contrasting degrees of average connectivity among patches.

The environmental conditions in each landscape were set to range within [0,5] to scale with species environmental optima (see Species pools and metacommunity dynamics and Supp. Material I) and varied in space either randomly, autocorrelated, or as a function of a linear gradient. To generate different levels of spatiotemporal environmental heterogeneity, we set local environmental conditions to follow a sinusoid function with 100 periods (e.g., 100 years), each composed of 12-time steps, plus a random error ∼*N*(0,0.1). We considered different amplitudes for the sinusoidal variation in the environment over time to allow for different levels of seasonality in the landscape. The resultant 60 (patches) × 1200 (time steps) matrix containing the environmental values was used to calculate an index of spatiotemporal environmental heterogeneity (SH/TH). SH/TH was calculated as the log of the ratio between the average variance of the environment in space (i.e., SH - average variance across columns) and the average variance of the environment through time (i.e., TH - average variance across rows). SH/TH values greater than 0 are observed in landscapes with stronger environmental heterogeneity in space than time (i.e., spatially heterogenous but aseasonal landscapes). When SH/TH values are close to 0, the level of environmental heterogeneity is similar in space and time; and values smaller than 0 indicate that environmental heterogeneity is stronger through time than in space (spatially homogenous but highly seasonal landscapes).

### Species pools and metacommunity dynamics

At the beginning of each of the simulation runs, we generated species pools containing 100 species each. Our operational definition of a species pool is the same as the one used in most empirical studies in metacommunity ecology, i.e., the set of species across all local communities within a metacommunity (*sensu* Fukami 2015). Species environmental optima (μ), environmental tolerance (σ), and dispersal ability (η; i.e., here defined as emigration propensity) were randomly drawn from continuous uniform distributions with ranges [0, 5], [0.1, 2], and [0.01, 0.5], respectively. This ensured that: (1) all simulation runs were seeded with species pools with the same initial distribution of trait values; (2) different combinations of σ and η (i.e., different life-history strategies) are equally likely to be observed across all initial metacommunities (e.g., specialists and poor dispersers, specialists and strong dispersers, generalists and poor dispersers, and generalists and strong dispersers).

Our mechanistic simulations, largely inspired by Büchi and Vuilleumier (2014), Shoemaker and Melbourne (2016), and Thompson et al. (2020), modeled metacommunity dynamics as a function of within-patch selection, dispersal in space and time, and ecological drift. Within-patch selection determined the influence of density-dependent and density-independent mechanisms on population dynamics. It was modeled as a Beverton-Holt growth model (Beverton and Holt 1957) with generalized Lotka-Volterra competition. We followed Büchi and Vuilleumier (2014) and scaled species maximum performances (i.e., growth rate) along the environmental gradient based on species-specific niche breadth σ. That ensured that all species would have similar cumulative growth rates along the environmental gradient regardless of their degree of ecological specialization (i.e., same areas below the performance-environment curves, see Supp. Material I Figure SI). As such, any artificial advantages that may have influenced the persistence and dominance of either specialists or generalists in different landscapes were removed.

Here we assumed stabilizing competition in which intraspecific competition is stronger than interspecific competition. This assumption is relevant because stabilizing competition favors coexistence by increasing the chances of locally rare species to keep positive population growth when locally dominant species have reached equilibrium at high abundances (i.e., the so-called “invasibility criterium” for coexistence Chesson 2000, Grainger et al. 2019). By assuming stabilizing competition, we increased the chances of species with different life-history strategies to coexist in suitable habitats and, consequently, persist in the metacommunity. While we acknowledge that competition types other than stabilizing may be important to metacommunity dynamics (see Thompson et al. 2020, Wisnoski and Shoemaker 2022), exploring their impact on the relationships between landscapes and species pools fall outside the scope of our current study. After the within-patch selection stage, the final population sizes were drawn from a Poisson distribution so that the effects of ecological drift into the outcomes of biotic interactions and habitat selection could be incorporated (following Shoemaker and Melbourne 2016).

Following within-patch selection, individuals in each patch at any given time are free to disperse based on their species-specific dispersal ability (*η*). To align our framework with recent developments in metacommunity ecology (e.g., Wisnoski et al. 2019, Wisnoski and Shoemaker 2022), we considered two types of dispersal: spatial and temporal (Buoro and Carlson 2014). Dispersal in space and time can be understood as alternative risk-spreading strategies that maximize species persistence in metacommunities under varying spatial and temporal environmental heterogeneity (Buoro and Carlson 2014; Holyoak et al. 2020). Considering temporal dispersal in metacommunities is relevant because, akin spatial dispersal, it promotes local and regional coexistence when local abiotic and biotic conditions favor different species in different periods (i.e., via temporal storage effects, Chesson 2000, Wisnoski and Shoemaker 2022).

Spatial dispersal was spatially explicit, meaning that any given patch is more likely to receive spatial immigrants from connected patches that are in proximity than distant patches. Temporal dispersal was meant to reproduce *in silico* any physiological (e.g., diapause, dormancy) and/or behavioral strategies (e.g., hiding in refugia) that buffer local extinctions by enabling organisms to avoid unfavorable biotic and abiotic conditions at the costs of neither reproducing nor consuming available resources. This was operationalized by temporally removing individuals from local communities and allowing them to return to the same patch in the future with probabilities that decayed exponentially over time at a fixed rate. That is, individuals that, for instance, temporally dispersed in time *t* are more likely to return sooner (e.g., *t+1*) than in distant time periods (e.g., *t+12*).

We simulated different dispersal scenarios to further determine the effects of different dispersal strategies (i.e., spatial vs temporal) on species persistence in the regional pool. This was done by manipulating the values of the parameter *DS* (Dispersal Strategy). *DS* is the probability of success in binomial trials that determined the number of individuals out of the total number of emigrants of any species in each patch at a given time that would temporally disperse; the remaining individuals would then disperse in space. We modeled three different scenarios. In the “Equal” scenario, species had the same probability of emigrating through either spatial or temporal dispersal (*DS*=*0.5*), whereas, in the “Mainly temporal dispersal” and “Mainly spatial dispersal” scenarios, *DS* was set as very high (0.99) and very low (0.01), respectively, for all species.

### Simulation iterations

For each parameter combination and dispersal scenario, we ran 20 independent replicates, yielding 12960 simulations runs in total (20 replicates x 8 SH/TH levels x 9 connectivity levels x 3 types of the spatial structure of environment x 3 dispersal scenarios). Population dynamics of the 100 initial species in the regional pool were carried across all 60 patches over 1200-time steps (i.e., 100 complete seasonal cycles). We seeded each patch in the first 120-time steps with species local (patch level) populations randomly drawn from a Poisson distribution (**λ** = 0.5). This allowed the chance for establishment and population growth for all species, provided that local abiotic and biotic conditions were suitable. In addition, the random placement of species populations across patches allowed those with similar habitat conditions to develop communities with dissimilar compositions due to random dispersal and priority effects (Thompson et al. 2020). To ensure that model summaries were carried out in stable rather than transient metacommunities, only communities in the last seasonal cycle (last 12 time-steps) were analysed. This decision was supported by sensitivity analyses (not shown) that demonstrated stabilization of species pools (rate of regional extinctions close to zero) after approximately 700 time-steps.

### Analyzing simulated metacommunities

We determined the dominant life-history of coexisting species in the regional pool (i.e., all species that persisted in the metacommunity in the last 12 time-steps) by calculating the regional-relative-abundance weighted mean of niche breadth and dispersal ability at the last seasonal cycle of each of the 12960 simulations. By doing so, we were able to derive theoretical predictions underlying the life-history strategies that maximized species persistence and dominance across different landscape types.

Our simulations were designed to generate insights into how landscape attributes and species pool characteristics influence inferences of the relative importance of different assembly mechanisms based on analytics commonly used to infer processes in empirical metacommunities (e.g., Cottenie 2005; Soininen 2014; Gálvez et al. 2022). To do so, we used variation partitioning (Borcard et al. 1992; Peres-Neto et al. 2006) to estimate the contribution of different groups of variables to the variation in community composition in the simulated data. Our simulations replicated data commonly collected in metacommunity studies and are analyzed using the same inferential approach. This enabled us to compare and contextualize our theoretical findings with those of empirical studies (e.g., Nishizawa et al. 2022 and see “Empirical support” below).

Variation partitioning was applied to the final patch-by-species-by-time matrix. We started by calculating pairwise compositional dissimilarities matrices and then using generalized dissimilarity models (GDMs, Ferrier et al. 2007) to fit these as a function of environment, spatial structure (here represented by positive Moran’s eigenvector maps, MEMs, calculated based on the patch geographic xy coordinates, Dray et al. 2006), and temporal structure (here represented by Asymmetric eigenvector maps, AEMs, Blanchet et al. 2008). Pairwise dissimilarities were calculated using the abundance-based Bray-Curtis index, which is widely used in observational studies and serves as the benchmark index for the link and variance functions of GDMs (see more details in Ferrier et al. 2007). Traditionally, the amount of variation in pairwise compositional dissimilarity matrices explained by environmental variables alone is considered a proxy for site selection via environmental filtering; the variation explained by spatial variables alone is linked to spatial autocorrelation on species distributions caused by the spatial structure of demographic events such as dispersal (Cottenie 2005, Beisner et al. 2006); the variation explained by temporal variables alone is linked to temporal autocorrelation on species dynamics associated with demographic events that are not related to extrinsic environmental factors (Legendre and Gauthier 2014). The variation explained by the covariation among variables (i.e., their joint contribution) was also estimated, albeit their association with specific ecological mechanisms is less clear in real empirical data that contain missing variables in the model (but see Peres-Neto et al. 2012 for possible interpretations). Finally, we ranked the relative importance of each component (in ascending order of importance) to facilitate comparisons across our 12960 independent simulation rounds.

All simulations and statistical analyses described above and below were conducted using R (v.4.1.0) (R Core Team 2023). AEMs and MEMs were calculated using the *adespatial* package (Dray et al. 2022). We used the *vegan* (Oksanen et al. 2020) package to calculate compositional dissimilarities and the *gdm* package (Manion et al. 2018) to fit the generalized dissimilarity models.

### Identifying interdependencies between landscapes characteristics, species pools attributes, and assembly mechanisms

We used path analysis to identify the causal interdependencies (pathways) between landscape attributes (i.e., exogenous variables: SH/TH, connectivity, and spatial structure of environment [ordinal; 1= Random, 2= Autocorrelated, 3 = Gradient]), species pool characteristics (i.e., mediators: niche breadth and dispersal ability), and the variation partitioning components (i.e., endogenous variables) across the 12960 simulation rounds. All predictors were standardized (mean = 0 and standard deviation =1) prior to model fit to allow comparing fitted relationships. The direction (i.e., positive/negative) and magnitude (i.e., standardized estimates) of pathways represented the general theoretical predictions derived from our simulations. Given the nature of the data used to fit the path analysis (i.e., simulated rather than empirical), the p-values of parameter estimates should not be used to determine the relevance of pathways. This is because the large number of replications required to detect patterns and trends in simulation models with stochastic components leads to decreased p-values even when effect sizes are negligible (see more in White et al. 2014). Moreover, simulation results from the same set of parameters may not be seen as true replicates. As such, the importance of paths in the fitted models was determined based solely on their standardized partial estimates (i.e., effect sizes).

To understand whether theoretical predictions change in direction and/or magnitude across dispersal scenarios (i.e., “Equal dispersal,” “Mainly temporal dispersal,” and “Mainly spatial dispersal”), we contrasted the results of a path analysis considering all dispersal scenarios pooled together (global model) against the results of a path analysis for each dispersal scenario separately. This assessment is relevant because it allowed us to understand whether predictions of the global model were prone to change in magnitude (i.e., quantitively) and/or direction (i.e., qualitatively) when considering metacommunities with different types of preferential dispersal. We used AIC_c_ to evaluate the fit of path models that considered different combinations of the linear and quadratic terms of predictors. The models that considered only linear terms (simplest) were identified as the best fitting path models in most cases. We used the *piecewiseSEM* R package (Lefcheck 2020) to fit the path models across dispersal scenarios.

### Empirical support: the assembly of moth metacommunities in tropical and temperate mountainous landscapes

To provide empirical support for the core theoretical predictions derived from our conceptual and simulation model (see below), we analyzed published data on moth metacommunities in two mountainous landscapes: The tropical Mount Cameroon (hereafter MTC, Maicher et al. 2019) and the temperate H.J. Andrews experimental forest (hereafter AEF, Miller and Jones 2005; Highland et al. 2013). Tropical and temperate mountainous landscape mountains drastically differ in spatiotemporal environmental heterogeneity (Zuloaga and Kerr 2017). Moths in both datasets were collected using light traps along an elevational gradient (MTC: from 35 to 2000 meters above sea level; AEF: from 400 to 1400 meters above sea level). Sampling was carried out at different moments of the reproductive season in each region (AEF: we used data May 2004 to October 2004; MTC: Different moments of the dry season and the transition between dry to wet and wet to dry seasons, see more details in references and Supp Material I). In general, moths are good spatial dispersers when adults and can persist in the landscape through prolonged juvenile diapause (Lees and Zilli 2019). As a result, both spatial and temporal dispersal are likely to influence the structure of moth metacommunities.

To estimate and contrast the degree of spatiotemporal environmental heterogeneity (SH/TH) we used sample coordinates to extract monthly temperature (mean, max, minimum monthly values) and precipitation data at 1 km × 1 km resolution (CHELSA data, Karger et al. 2017). We then performed a principal component analysis on the temperature variables and log-transformed precipitation (and then standardized to mean = 0 and standard deviation = 1) and used the sample scores on the first two PC axes as a proxy of the climatic conditions of each site across different time periods of the year. In the MTC, PC 1 accounted for 76.2% of the variance in the climate data and 2 accounted for 22.1% (cumulative proportion of 98.3%). In the AEF, PC 1 accounted for 74.2% of the variance in the climate data and 2 accounted for 20.7% (cumulative proportion of 94.9%).

We estimated the climatic tolerance of each species through the tolerance index of Doledec (2000) using the package “ade4” (Thioulouse et al. 2018). This index estimates species-specific climatic tolerance (i.e., niche breadth) based on the dispersal of samples that contain the target species in the multivariate climatic space. We pooled together data on moths and climate variables to estimate climatic tolerance in the same multivariate space. By doing so, we could contrast the degree of ecological specialization of species observed in both datasets. Lastly, we inferred the relative importance of different assembly mechanisms in both landscapes using variation partitioning (following the same steps described in *Analyzing simulated metacommunities*). This was done by estimating the variation in the community composition data explained by climate (PC1 and PC2), space (spatial MEMs), and time (temporal AEMs).

## Results and Discussion

For purposes of tractability and synthesis, we focused our discussion on the two pathways with the highest importance (i.e., larger partial coefficients) for each mediator (i.e., niche breadth and dispersal ability) and endogenous (i.e., components of the variation partitioning) variables in the fitted path models (Figures 3 and 4). Due to qualitative similarities in the results of path models fitted considering each dispersal scenario separately (see tables SI-SIV in Supp. Material I), we only report the results of the fitted path models considering data on all dispersal scenarios pooled together. However, we also highlight and discuss cases in which there were differences in the direction of pathways across dispersal scenarios. Additionally, our discussion focuses on the direction of the two most relevant pathways linking exogenous (i.e., landscape attributes) and moderator variables to the isolated contribution of environment, spatial MEMs, and temporal AEMs on variation partitioning. By doing so, we could determine and understand the influence of landscapes characteristics and species pool attributes on the assembly process when assessed through components of the variation partitioning that have well-established associations with distinct ecological mechanisms (Ovaskainen et al. 2019; Viana and Chase 2019*b*; Guzman et al. 2022). Nonetheless, the complete set of numerical relationships estimated by path analyses across all dispersal scenarios and considering the full set of variation partitioning components can be found in Supp. Material I.

**Figure 3:**
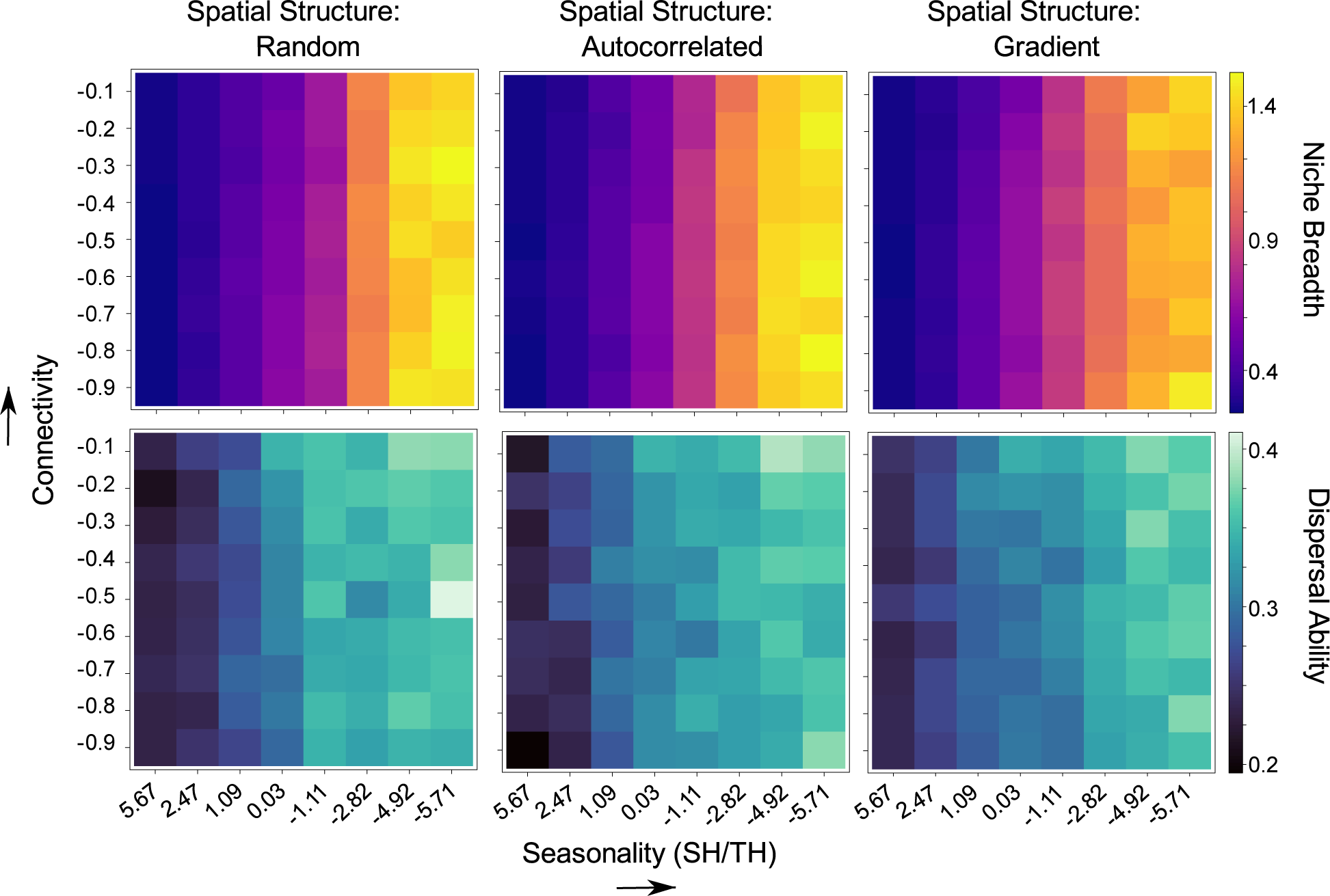
Landscape attributes determine the dominant life-history strategies in species pools. Aseasonal (SH/TH > 0) and poorly connected landscapes selected for environmental specialists (i.e., narrow niche breadth) that were also weak dispersers (i.e., low dispersal ability). Seasonal (SH/TH < 0) and highly connected landscapes favored the dominance of environmental generalists that were also strong dispersers. These are the results reported for the “Equal” dispersal scenario where species were equally likely to disperse spatially and temporally. The results for the “Mostly Spatial” and “Mostly Temporal” dispersal scenarios are reported in Supp. Material I, Figures SII-SIII.

**Figure 4.**
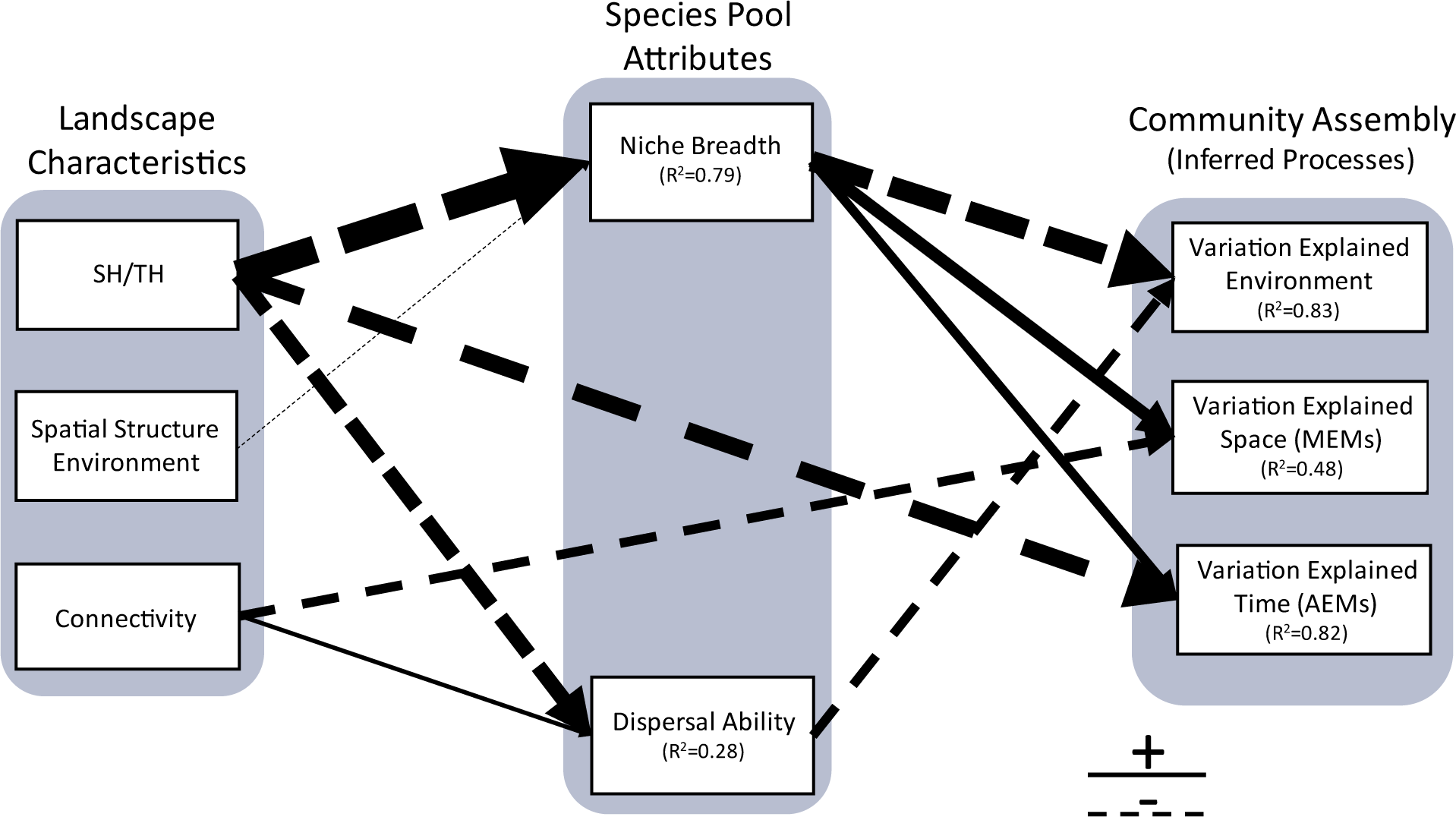
Theoretical predictions derived from path analysis considering the relationships between landscape attributes, the dominant life-history strategies in species pools, and the variation partitioning components. Only the two pathways with the largest effects on each moderator (i.e., species pool characteristics) and exogenous (community assembly) variables are reported here. Arrow widths are proportional to the effects sizes estimated. Results reported considering all dispersal scenarios pooled together. The numerical results obtained from path analyses considering each dispersal scenario pooled together and separately are reported in Supp. Information 1 (Tables SI-SIV). SH/TH = Spatiotemporal environmental heterogeneity index

### Theoretical Predictions: landscape attributes influence the degree of ecological specialization and dispersal ability of dominant species in the regional pool

Our simulation clearly showed that seasonality (measured as the ratio between spatial and temporal heterogeneity, SH/TH) was the most important factor determining the degree of ecological specialization of the dominant species in the regional pool (Figures 3 and 4). Ecological specialization was favored in aseasonal landscapes where environmental heterogeneity was higher in space than in time (SH/TH > 0). In contrast, ecological generalization was favored in highly seasonal landscapes where environmental heterogeneity is higher in time than in space (SH/TH < 0). Notably, we observed an increase in the persistence of ecological specialists in seasonal landscapes when we considered the “Mainly temporal dispersal” scenario (see Figure SII). These findings highlight the importance of temporal dispersal to the coexistence of specialists and generalists in (temporally) fluctuating environments (Chesson 2000; Wisnoski and Shoemaker 2022).

The spatial structure of the environment also influenced the overall niche breadth of species pools, but this relationship was relatively weak (Figures 3 and 4, Supp. Material I Table SI). When environmental conditions were randomly distributed across the landscape, an increase in the dominance of generalists in the regional pool was observed. The lack of spatial structure in habitat conditions increases the chances of environmental specialists being isolated in patches surrounded by unsuitable habitat conditions (Büchi and Vuilleumier 2014; Fournier et al. 2017). Since isolation increases populations’ chances to become locally extinct due to demographic stochasticity, the lack of spatial structure of environmental conditions should increase the isolation and, consequently, the local and regional extinction of ecological specialists.

Dispersal ability is influenced by seasonality and the level of connectivity in landscapes (Figures 3 and 4), though the strength of these relationships varied across dispersal scenarios (see Supp. Material I, Tables SI-SIV). When metacommunity dynamics were primarily driven by spatial dispersal (i.e., the “Mostly spatial” scenario), dispersal ability increased at intermediate levels of seasonality, but increased linearly with physical connectivity. This finding suggests that highly connected landscapes reduce the risks associated with spatial dispersal, by increasing the likelihood of species successfully colonizing suitable patches when environmental heterogeneity is equally strong in space and time (see Kubisch et al. 2014 and references within). In contrast, when species mainly dispersed over time (the “Mostly Temporal” scenario) or had equal chances of dispersing in space and time (the “Equal” scenario), spatiotemporal environmental heterogeneity emerged as the most important landscape attribute, exhibiting a negatively correlation with the dominant level of dispersal in the metacommunity. This implies that species’ abilities to disperse in time is critical to their persistence in highly seasonal landscapes. Additionally, these results illustrate that spatial and temporal dispersal are risk-spreading strategies favored by different levels of spatial environmental heterogeneity (Buoro and Carlson 2014).

Taken together, our findings provide theoretical support for macroecological hypotheses and ecogeographic rules invoked to explain latitudinal clines on species’ ecological specialization and dispersal ability. Our results demonstrate that these patterns can emerge as a consequence of ecological processes operating only at metacommunity scales (i.e., no need to evoke evolution). For instance, Janzen’s seasonality hypothesis posits that the high elevational stratification of climate and the low seasonality of tropical mountainous landscapes should favor the dominance of environmental specialists whose spatial distributions are restricted to different types of climate (Janzen 1967). Conversely, strong seasonality in temperate regions should favor the dominance of species that have broad physiological tolerances and are less sensitive to spatial variation in climate (Sheldon and Tewksbury 2014).

Our conceptual and simulation models successfully replicated the expected relationship between environmental temporal variability and the optimal level of dispersal ability in the regional pools that shape metacommunities (e.g., Jocque et al. 2010; Sheard et al. 2020). We found that weak dispersers highly specialized in local conditions dominate local communities and increase their persistence in the regional pool when the environment is temporally homogenous. In contrast, high temporal variability of environmental conditions favors species with increased dispersal ability that can escape from temporally unsuitable local conditions (Figure 3). Given that (i) geographic ranges reflect the ability of species to disperse (Alzate and Onstein 2022, but see Lester et al. 2007); and (ii) the strength of seasonality (particularly in temperature) increases from the equator to the poles, our simulations were able to recreate the underlying conditions that lead to an increase in range size as a function of latitude, as predicted by the Rapoport’s rule (Gaston et al. 2008).

### Theoretical Predictions: landscape and species pool attributes influence inferences about the relative importance of assembly mechanisms

Our theoretical framework allowed understanding how the characteristics of the landscape where metacommunity dynamics take place and the attributes of species pools that form metacommunities can influence empirical inferences about the relative importance of assembly mechanisms (Figure 4). The unique contribution of the environment (via variation partitioning) captures the importance of species-environment sorting in community assembly (Cottenie 2005, Ovaskainen et al. 2019). Path analyses fit to the results of all dispersal scenarios pooled together suggest that the influence of environmental variation on community composition had an negative relationship with the degree of niche breadth and dispersal ability of the dominant species in the regional pool.

Not surprising, these results indicate that the strength of environmental selection on community composition increases when species pools are dominated by environmental specialists that are also weak disperses (i.e., the species sorting paradigm). However, it is noteworthy that the direction of the relationship between dispersal ability and the contribution of environmental variables in the variation partitioning is not constant across dispersal scenarios (see Supp. Material I, tables SI-SIV). When spatial dispersal is as, or more, frequent than temporal dispersal (i.e., the Equal and Mostly Spatial scenarios), dispersal ability increases the relative importance of environmental selection in community assembly. This suggests that the strength of environment-community composition relationships can increase with spatial dispersal when it increases the probability of specialists reaching and persisting in all suitable habitats available. Conversely, the relationship becomes negative when dispersal is constrained to be mostly temporal (i.e., under the Mostly Temporal scenario). This pattern suggests that “seed banks” buffer the extinction of populations in unsuitable local conditions, decreasing the strength of the match between community composition and environment (Wisnoski et al. 2019).

The importance of the unique contribution of space (spatial MEMs) is typically associated with the influence of dispersal limitation in community assembly. We observed that the importance of space increased with niche breadth but decreased with landscape connectivity (Figure 4). This result indicates that spatial autocorrelation in community composition that cannot be associated with the spatial structure of the environment emerges when species that do not respond strongly to environmental gradients are physically limited to dispersing to neighboring patches.

The amount of variation in the community matrix explained by the unique contribution of temporal variation (AEMs) is typically associated with temporal autocorrelation in population dynamics generated by neutral mechanisms such as demographic stochasticity (Legendre and Gauthier 2014). Our simulations indicate that the importance of neutral dynamics tend to increase when temporal environmental variation is weak (i.e., aseasonal landscapes) and generalists dominate the metacommunity (Figure 4). Under these conditions, stochastic events of colonization events and local extinctions outweighs the influence of niche-based mechanisms in generating temporal autocorrelation in population dynamics. These results are aligned with previous empirical studies demonstrating that the stochastic signatures of temporal changes in community composition increase in aseasonal landscapes where environmental heterogeneity is stronger in space than in time (e.g., Khattar et al. 2021).

In summary, our model demonstrates that landscape attributes and species pool characteristics are strongly associated and should not be considered as independent axes in the assembly process. They also suggest that this link should lead to variation in the relative importance of assembly mechanisms along broad-scale gradients that encompass variation in key landscape attributes. In the next section, we provide empirical evidence for key predictions derived from our models and illustrate how our conceptual framework can generate insights into the geography of community assembly.

### Empirical Support

A strong prediction derived from our theoretical model is that in landscapes where environmental heterogeneity is relatively greater in space than in time (aseasonal landscapes, SH/TH > 0), species pools will be dominated by specialists (Figures 3 and 4). Consequently, environmental selection should be the primary mechanism driving community assembly in these landscapes. In contrast, generalists will dominate species pools in landscapes where environmental conditions change relatively more in time than in space (seasonal landscapes, SH/TH < 0). Consequently, it is reasonable to infer that mechanisms beyond environmental selection alone likely play a significant role in driving community assembly.

As typically observed in tropical mountains (Figure 5 panel A), climate (PC1 scores) varied more across elevations than over time in tropical Mount Cameroon (SH/TH = 2.12). In contrast, PC1 scores varied more in time than across elevations in the temperate Andrews Experimental Forest (SH/TH= −2.56).

**Figure 5:**
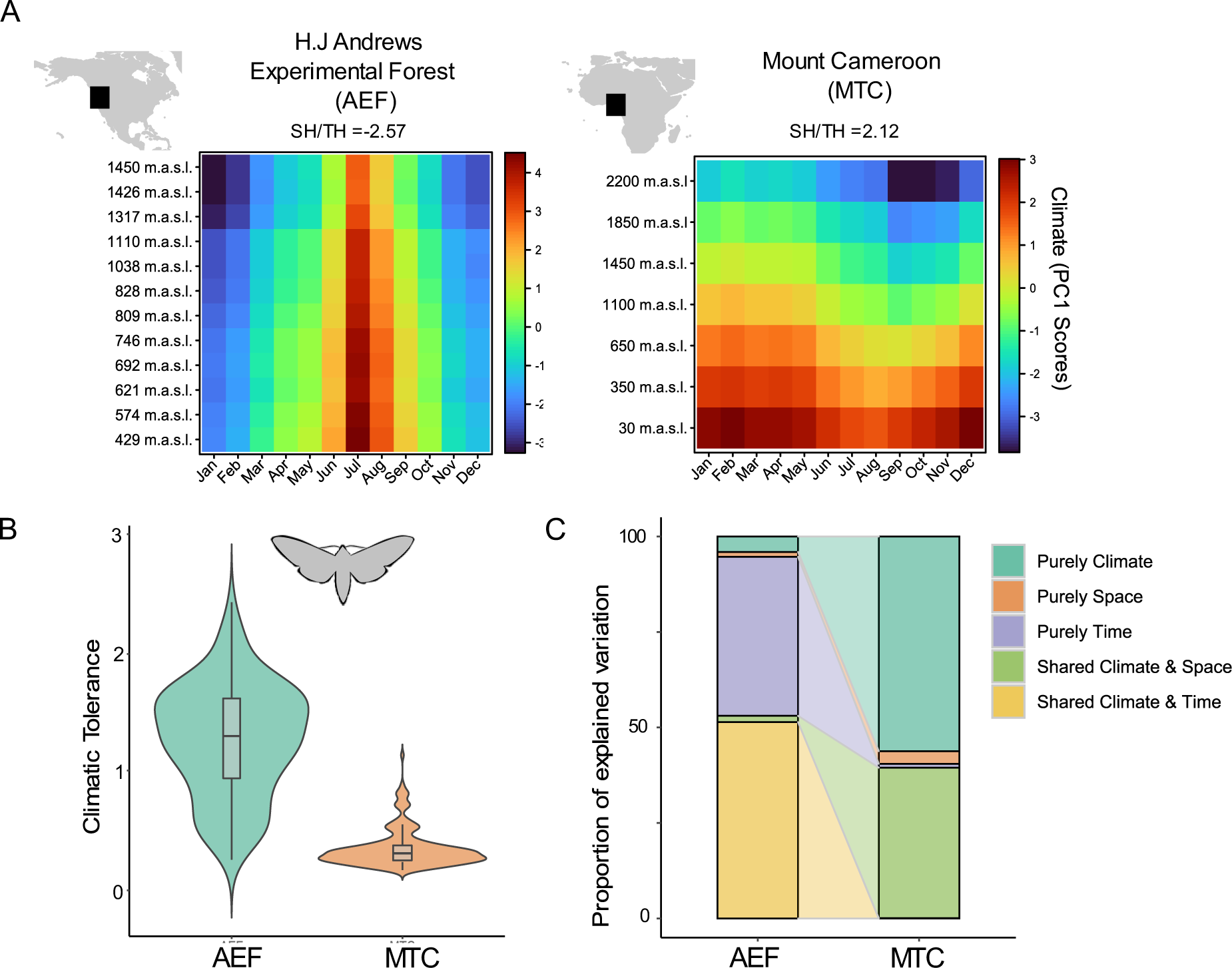
We analyzed the assembly of moth metacommunities in two different mountainous landscapes: The tropical and relatively aseasonal Mount Cameroon (MTC, SH/TH >0) and the temperate and relatively seasonal H.J Andrews experimental forest (AEF, SH/TH <0) (panel A). In the MTC, the regional pool is dominated by climate specialists, while climate generalists dominate the regional pool in the AEF (B). As such, deterministic species-environment sorting is the primary driver of community assembly in the MTC, whereas temporal autocorrelation on population dynamics caused by neutral dynamics and the temporal structure of climate are the main drivers of variation in community composition in the AEF (C). Shared contributions of climate, space, time, and time and space were extremely small in both metacommunities (< 0.1 %) and, therefore, were omitted in the plot.

In Figure 5 (panel B), we contrasted the degree of climatic tolerance of the dominant species in the regional pool of both landscapes. Here we report the results when the 33% most abundant species of each pool were considered to estimate the average niche breadth of each metacommunity (AEJ = 121 species; MTC = 185 species). The results were qualitatively the same when only the 10% or 25% of the most regionally abundant species were considered (not shown). As predicted, the aseasonal MTC bear a species pool dominated by climate specialists (i.e., species with a relatively narrow niche breadth). Conversely, the seasonal AEF favors the dominance of climate generalists in the pool (i.e., species with a relatively broad niche breadth).

As predicted (Figure 5, panel C), community composition in the aseasonal MTC (where specialists dominate the species pool) was mainly explained by climate variation alone. This pattern suggests a strong influence of environmental selection in community assembly in aseasonal landscapes. In contrast, beta diversity in the highly seasonal AEF (where generalists dominate the species pool) was mainly explained by temporal variation and their correlation with climate. This pattern suggests that temporal autocorrelation on population dynamics caused by neutral dynamics and climate’s temporal structure are the main drivers of variation in species composition among local communities (Legendre and Gauthier 2014).

## Conclusions, assumptions, and future directions

Our goal was to propose a conceptual framework for metacommunity assembly that recognizes the dependency of species pool attributes on landscapes characteristics, and how these two components together determine the relative important of assembly mechanisms. By doing so, we derived testable predictions underlying geographical patterns of metacommunity assembly when inferred from empirical data. Moreover, our study demonstrates that process-based simulations designed to study the drivers of metacommunity structure at the landscape scale can also recreate the patterns underlying relevant hypotheses in biogeography and macroecology.

However, we recognize that our conceptual framework and theoretical model did not consider other aspects of landscapes that are known to influence the coexistence of specialists and generalists, even though these could be incorporated in future model versions. For instance, recent empirical studies have shown that the spatial frequency of climate conditions at large scales is a relevant factor determining the degree of ecological specialization of species pools (Fournier et al. 2020).

Lastly, our framework for the geography of metacommunities represents a “bottom-up” perspective in which species pool dynamics are driven by mechanisms operating only at the landscape scale while deliberately discounting the effects of evolutionary and historical mechanisms operating at biogeographic scales. That said, our proposed framework is useful in further syntheses investigating the considerable variation in the relative importance of mechanisms observed in empirical metacommunities at different parts of large-scale ecological gradients. Future studies should explore how evolutionary processes mediate the relationships between dominant life-history strategies, landscape attributes, and assembly mechanisms at the metacommunity level (e.g., Mittelbach and Schemske 2015).

## Supporting information

Code and Data to run simulations and generate figures

## Acknowledgements

We thank Paul Savary and Pedro Henrique Pereira Braga for their insightful comments on the initial drafts of this manuscript. The AEF dataset was obtained upon work supported by the H.J. Andrews Experimental Forest and Long Term Ecological Research (LTER) program under the NSF grant LTER8 DEB-2025755. GK is supported by the Canada Research Chair in Spatial Ecology and Biogeography held by PPN.

## Notes

### Competing Interest Statement

The authors have declared no competing interest.

## References

Alzate, A., and R. E. Onstein. 2022. Understanding the relationship between dispersal and range size. Ecology Letters 2303–2323.

Beverton, R. J. H., and S. J. Holt. 1957. On the Dynamics of Exploited Fish Populations (Fish and F.). Springer Science and Business Media, London.

Blanchet, F. G., P. Legendre, and D. Borcard. 2008. Modelling directional spatial processes in ecological data. Ecological Modelling 215:325–336.

Borcard, D., P. Legendre, and P. Drapeau. 1992. Partialling out the Spatial Component of Ecological Variation. Ecology 73:1045–1055.

Büchi, L., P.-A. Christin, and A. H. Hirzel. 2009. The influence of environmental spatial structure on the life-history traits and diversity of species in a metacommunity. Ecological Modelling 220:2857–2864.

Büchi, L., and S. Vuilleumier. 2014. Coexistence of specialist and generalist species is shaped by dispersal and environmental factors. American Naturalist 183:612–624.

Buoro, M., and S. M. Carlson. 2014. Life-history syndromes: Integrating dispersal through space and time. Ecology Letters 17:756–767.

Chesson, P. 2000. Mechanisms of maintenance os species diversity. Annual Review of Ecology and Systematics 31:343–366.

Cottenie, K. 2005. Integrating environmental and spatial processes in ecological community dynamics. Ecology Letters 8:1175–1182.

Dray, S., D. Bauman, F. G. Blanchet, D. Borcard, S. Clappe, G. Guenard, T. Jombart, et al. 2022. “Adespatial: Multivariate Multiscale Spatial Analysis (version 0.3-7).”

Fournier, B., N. Mouquet, M. A. Leibold, and D. Gravel. 2017. An integrative framework of coexistence mechanisms in competitive metacommunities. Ecography 40:630–641.

Fournier, B., H. Vázquez-Rivera, S. Clappe, L. Donelle, P. H. P. Braga, and P. R. Peres-Neto. 2020. The spatial frequency of climatic conditions affects niche composition and functional diversity of species assemblages: the case of Angiosperms. (C. Violle, ed.) Ecology Letters 23:254–264.

Fukami, T. 2015. Historical Contingency in Community Assembly: Integrating Niches, Species Pools, and Priority Effects. Annual Review of Ecology, Evolution, and Systematics 46:1–23.

Gálvez, Á., P. R. Peres-Neto, A. Castillo-Escrivà, F. Bonilla, A. Camacho, E. M. García-Roger, S. Iepure, et al. 2022. Inconsistent response of taxonomic groups to space and environment in mediterranean and tropical pond metacommunities. Ecology 104:1–16.

Gaston, K. J., S. L. Chown, and K. L. Evans. 2008. Ecogeographical rules: elements of a synthesis 483–500.

Ghalambor, C. K. 2006. Are mountain passes higher in the tropics? janzen’s hypothesis revisited. Integrative and Comparative Biology 46:5–17.

Gilbert, B., and J. R. Bennett. 2010. Partitioning variation in ecological communities: do the numbers add up? Journal of Applied Ecology 47:1071–1082.

Grainger, T. N., J. M. Levine, and B. Gilbert. 2019. The Invasion Criterion: A Common Currency for Ecological Research. Trends in Ecology and Evolution 34:925–935.

Guzman, L. M., P. L. Thompson, D. S. Viana, B. Vanschoenwinkel, Z. Horváth, R. Ptacnik, A. Jeliazkov, et al. 2022. Accounting for temporal change in multiple biodiversity patterns improves the inference of metacommunity processes. Ecology 1–16.

Henriques-Silva, R., B. Frédéric, V. Calcagno, M. C. M. C. M. M. C. M. Urban, P. R. P. Peres-neto, F. F. Boivin, V. Calcagno, et al. 2015. On the evolution of dispersal via heterogeneity in spatial connectivity. Proceedings of the Royal Society B: Biological Sciences 282:21–23.

Highland, S. A., J. C. Miller, and J. A. Jones. 2013. Determinants of moth diversity and community in a temperate mountain landscape: Vegetation, topography, and seasonality. Ecosphere 4.

Holyoak, M., T. Caspi, and L. W. Redosh. 2020. Integrating Disturbance, Seasonality, Multi-Year Temporal Dynamics, and Dormancy Into the Dynamics and Conservation of Metacommunities. Frontiers in Ecology and Evolution 8:1–17.

Janzen, D. H. 1967. Why Mountain Passes are Higher in the Tropics. The American Naturalist 101:233–249.

Jocque, M., R. Field, L. Brendonck, and L. De Meester. 2010. Climatic control of dispersal-ecological specialization trade-offs: A metacommunity process at the heart of the latitudinal diversity gradient? Global Ecology and Biogeography 19:244–252.

Karger, D. N., O. Conrad, J. Böhner, T. Kawohl, H. Kreft, R. W. Soria-Auza, N. E. Zimmermann, et al. 2017. Climatologies at high resolution for the earth’s land surface areas. Scientific Data 4:170122.

Khattar, G., M. Macedo, R. Monteiro, and P. Peres-Neto. 2021. Determinism and stochasticity in the spatial–temporal continuum of ecological communities: the case of tropical mountains. Ecography 44:1391–1402.

Koffel, T., K. Umemura, E. Litchman, and C. A. Klausmeier. 2022. A general framework for species-abundance distributions: Linking traits and dispersal to explain commonness and rarity. Ecology Letters.

Kubisch, A., R. D. Holt, H.-J. Poethke, and E. A. Fronhofer. 2014. Where am I and why? Synthesizing range biology and the eco-evolutionary dynamics of dispersal. Oikos 123:5–22.

Lees, D. C., and A. Zilli. 2019. Moths: A complete Guide to Biology and Behavior. Natural History Museum: London, UK.

Lefcheck, J. S. 2020. PiecewiseSEM (R package version 2.1.2).

Legendre, P., and O. Gauthier. 2014. Statistical methods for temporal and space–time analysis of community composition data. Proceedings of the Royal Society B: Biological Sciences 281:20132728.

Lester, S. E., B. I. Ruttenberg, S. D. Gaines, and B. P. Kinlan. 2007. The relationship between dispersal ability and geographic range size. Ecology Letters 10:745–758.

Maicher, V., S. Sáfián, M. Murkwe, S. Delabye, Ł. Przybyłowicz, and P. Potocký. 2019. Seasonal shifts of biodiversity patterns and species’ elevation ranges of butterflies and moths along a complete rainforest elevational gradient on Mount Cameroon. Dryad.

Manion, G., M. Lisk, S. Ferrier, D. Nieto-Lugilde, K. Mokany, and M. C. Fitzpatrick. 2018. gdm: Generalized Dissimilarity Modeling. R package ver. 1.3.11.

Miller, J., and J. A. Jones. 2005. Spatial and temporal distribution and abundance of moths in the Andrews Experimental Forest, 1994 to 2008. H. J. Andrews Experimental Forest. Forest Science Data Bank, Corvallis. http://andlter.forestry.oregonstate.edu/data/abstract.aspx?dbcode=SA015.

Mittelbach, G. G., and D. W. Schemske. 2015. Ecological and evolutionary perspectives on community assembly. Trends in Ecology & Evolution 30:241–247.

Moritz, C., C. N. Meynard, V. Devictor, K. Guizien, C. Labrune, J. Guarini, and N. Mouquet. 2013. Disentangling the role of connectivity, environmental filtering, and spatial structure on metacommunity dynamics. Oikos 1401–1410.

Mouquet, N., and M. Loreau. 2003. Community Patterns in Source-Sink Metacommunities. The American naturalist.

Myers, J. A., J. M. Chase, I. Jiménez, P. M. Jørgensen, A. Araujo-Murakami, N. Paniagua-Zambrana, and R. Seidel. 2013. Beta-diversity in temperate and tropical forests reflects dissimilar mechanisms of community assembly. Ecology Letters 16:151–157.

Nishizawa, K., N. Shinohara, M. W. Cadotte, and A. S. Mori. 2022. The latitudinal gradient in plant community assembly processes: A meta-analysis. (J. Chase, ed.) Ecology Letters 25:1711–1724.

Oksanen, J., F. G. Blanchet, M. Friendly, R. Kindt, P. Legendre, D. McGlinn, P. R. Minchin, et al. 2020. vegan: Community Ecology Package.

Ovaskainen, O., J. Rybicki, and N. Abrego. 2019. What can observational data reveal about metacommunity processes? Ecography 42:1877–1886.

Peres-Neto, P. R., and P. Legendre. 2010. Estimating and controlling for spatial structure in the study of ecological communities. Global Ecology and Biogeography 19:174–184.

Peres-Neto, P. R., P. Legendre, S. Dray, and D. Borcard. 2006. Variation partitioning of species data matrices: estimation and comparison of fractions. Ecology 87:2614–25.

Peres-Neto, P. R., M. A. Leibold, and S. Dray. 2012. Assessing the effects of spatial contingency and environmental filtering on metacommunity phylogenetics. Ecology 93:14–30.

Qian, H., and R. E. Ricklefs. 2012. Disentangling the effects of geographic distance and environmental dissimilarity on global patterns of species turnover. Global Ecology and Biogeography 21:341–351.

R Core Team. 2023.R: A Language and Environment for Statistical Computing. Vienna, Austria.

Sheard, C., M. H. C. Neate-Clegg, N. Alioravainen, S. E. I. Jones, C. Vincent, H. E. A. MacGregor, T. P. Bregman, et al. 2020. Ecological drivers of global gradients in avian dispersal inferred from wing morphology. Nature Communications 11.

Sheldon, K. S., R. B. Huey, M. Kaspari, and N. J. Sanders. 2018. Fifty Years of Mountain Passes: A Perspective on Dan Janzen’s Classic Article. The American Naturalist 191:553–565.

Sheldon, K. S., and J. J. Tewksbury. 2014. The impact of seasonality in temperature on thermal tolerance and elevational range size. Ecology 95:2134–2143.

Shoemaker, L. G., and B. A. Melbourne. 2016. Linking metacommunity paradigms to spatial coexistence Mechanisms. Ecology 97:2436–2446.

Soininen, J. 2014. A quantitative analysis of species sorting across organisms and ecosystems. Ecology 95:3284–3292.

Thioulouse, J., S. Dray, A.--B. Dufour, A. Siberchicot, T. Jombart, and S. Pavoine. 2018. Multivariate Analysis of Ecological Data with {ade4}. Springer.

Thompson, P. L., L. M. Guzman, L. De Meester, Z. Horváth, R. Ptacnik, B. Vanschoenwinkel, D. S. Viana, et al. 2020. A process-based metacommunity framework linking local and regional scale community ecology. Ecology Letters 23:1314–1329.

Vellend, M., D. S. Srivastava, K. M. Anderson, C. D. Brown, J. E. Jankowski, E. J. Kleynhans, N. J. B. Kraft, et al. 2014. Assessing the relative importance of neutral stochasticity in ecological communities. Oikos 123:1420–1430.

Viana, D. S., and J. M. Chase. 2019. Spatial scale modulates the inference of metacommunity assembly processes. Ecology 100:1–9.

White, J. W., A. Rassweiler, J. F. Samhouri, A. C. Stier, and C. White. 2014. Ecologists should not use statistical significance tests to interpret simulation model results. Oikos 123:385–388.

Wisnoski, N. I., M. A. Leibold, and J. T. Lennon. 2019. Dormancy in metacommunities. American Naturalist 194:135–151.

Wisnoski, N. I., and L. G. Shoemaker. 2022. Seed banks alter metacommunity diversity: The interactive effects of competition, dispersal and dormancy. Ecology Letters 25:740–753.

Zuloaga, J., and J. T. Kerr. 2017. Over the top: do thermal barriers along elevation gradients limit biotic similarity? Ecography 40:478–486.

