## Supplemental Information for "The geography of metacommunities: landscape characteristics drive geographic variation in the assembly process through selecting species pool attributes"

**Running title:** The geography of metacommunities

**Authors:** Gabriel Khattar & Pedro Peres-Neto

**Affiliations:** Laboratory of Community and Quantitative Ecology, Department of Biology, Concordia University, Montreal, QC, Canada

**Supplementary Material I**

**Extended Model description**

**Simulated landscapes**

We started by randomly distributing 60 patches in a geographic space defined by x and y coordinates ranging from 0 to 60. The degree into which any given two patches are physically connected decays exponentially with Euclidean distance (**Δ**S) according to the following kernel function:

${Connectivity}_{i,j}=exp(-c*$ **Δ**S_ij_) (eq. 1)

where the term *c* is the rate at which connectivity decays with spatial distance. *Connectivity* values below a threshold of 10^-4^ were truncated to 0 so that individuals could not move between the focal pair of patches, thus creating patches that are truly disconnected (as in Fournier et al. 2017). By varying *c* (here from 0.1 to 0.9) but keeping the threshold constant, we could generate landscapes with contrasting degrees of average connectivity among patches. The degree of connectivity between any given pair of patches (eq.1) defines the weighted probabilities of spatial dispersal between these patches (see “Species pools and metacommunity dynamics” below).

The environmental conditions in each landscape were set to range in the interval [0,5] to scale with species environmental optima (see below) and varied in space across three different spatial types: random, autocorrelated, and linear gradient. In random landscapes, the initial environmental value of each patch was randomly drawn from a continuous uniform distribution *U(0,5).* In autocorrelated landscapes, we modeled environmental conditions using a multivariate normal distribution (*mu*=0) with a covariance matrix defined as the exponential decay of environmental similarity between pairs of patches with distance **Δ**S and the constant phi (here, set at 0.15). In gradient landscapes, we modeled the initial environment of each site as a linear function of their x and y coordinates.

To simulate seasonal environmental variation, we set local environmental conditions to follow a sinusoid function with 100 periods, each composed of 12-time steps (e.g., 100 years) plus a random error $N\left( 0,.1 \right)$. The amplitude of the sinusoidal variation in the environment over time was modulated by a multiplicative factor *s* (constant across all patches). As such, the higher the value assigned to *s*, the higher the seasonal variation in environmental conditions. The final 60 (patches) x 1200 (times) matrix containing the environmental values was used to calculate an index of spatiotemporal environmental heterogeneity, hereafter SH/TH. SH/TH was calculated as the log of the ratio between the average variance of the environment in space (i.e., SH - average variance across columns) and the average variance of the environment through time (i.e., TH - average variance across rows). SH/TH values higher than 0 are observed in landscapes environmental heterogeneity is stronger in space than in time (i.e., spatially heterogenous but aseasonal landscapes); values close to 0 indicate that the level of environmental heterogeneity is similar in space and time; values lower than 0 indicate that environmental heterogeneity is stronger in time than in space (spatially homogenous but highly seasonal landscapes).

***Species pools and metacommunity dynamics***

Our operational definition of species pool is the same as the one used in most empirical studies in metacommunity ecology: the set of all species that can be observed in the landscape where the metacommunity is located (*sensu* Fukami 2015) . At the beginning of each simulation round, we generated species pools of 100 species each (Sp). Species environmental optima (μ), environmental tolerance (σ), and dispersal ability (i.e., here defined as the species` emigration propensity, η) were randomly drawn from independent continuous uniform distributions with ranges of [0, 5], [0.1, 2], and [0.01, 0.5], respectively. By doing so, we ensured that: (1) all simulation rounds were seeded with species pools with the same initial distribution of trait values: (2) different combinations of σ and η (i.e., different life-history strategies) are equally likely to be observed across all landscapes (e.g., specialists and poor dispersers, specialists and strong dispersers, generalists and poor dispersers, and generalists and strong dispersers).

Considering that *N_i,j,t_* is the abundance of species *i* in site *j* in time *t*, population dynamics is governed by:

$N_{i,j,t}=Poisson\left( N_{i,j,t-1}*P_{i,j,t} \right)-{(E}_{i,j,t total})+(I_{i,j,t total})$ (eq. 2)

The first term of eq. 2 is a modified version of the commonly used Beverton-Holt model that equates discrete population growth as a function of within-patch selection and ecological drift (i.e., demographic stochasticity). $P_{i,j,t}$ is the local performance (i.e., growth rate) of species *i* when conditioned to competition and habitat selection in site *j* and time *t*, and is modeled as:

$P_{i,j,t}=R_{i,j,t}*\frac{1}{(1+\alpha_{intra}N_{i,j,t}+\alpha_{inter}\sum_{k\neq i}^{Sp} N_{k,j,t})}$ (eq. 3)

where *R_i,j,t_* is the influence of local environmental conditions on species performance given by a Gaussian response:

$R_{i,j,t}=u* \frac{1}{\sigma_{i}\sqrt{2\pi}}*\exp\left( \frac{-{({Env}_{j,t}-\mu_{i})}^{2}}{2{\sigma_{i}}^{2}} \right)$(eq. 4)

$where {Env}_{jt}$ represents local abiotic conditions. The term $1/(\sigma_{i} \surd2\pi)$ scales species responses to the environmental gradient, ensuring that, when competition is negligible (see below), all species that share the same environmental optima have identical cumulative growth rates along the environmental gradient regardless of their niche breadth (i.e., same areas below the performance-environment curves, as in Büchi and Vuilleumier 2014). $u$ (set at 10 after pre trials that showed it allows the persistence of a larger number of species over time) is a scaling factor that ensures that all species are able to reach positive growth (i.e., $P_{i,j,t}$>1) when local abiotic and biotic conditions are suitable.
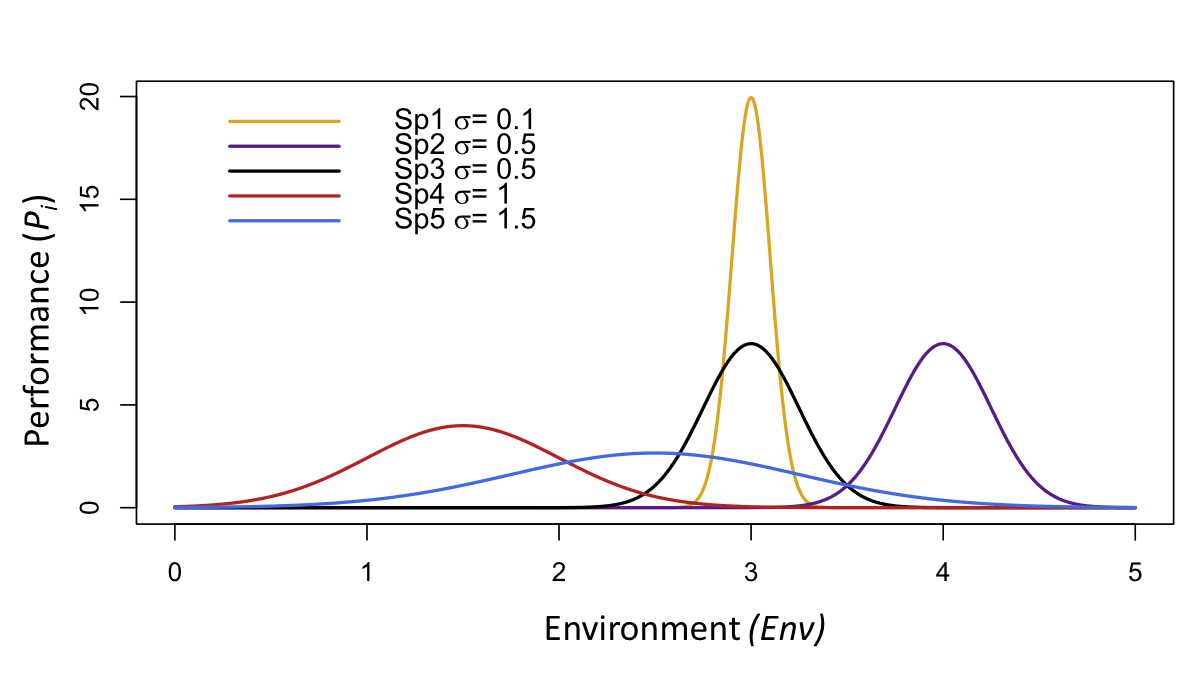

Figure SI: Performances of five different species (Sp1 – Sp5) at different environmental values ($Env$) when competition is negligible (i.e., at low population sizes). Species performances peak at their environmental optima, but the height and shape of performance curves depend on their niche breadth (σ). Sp1 has the highest level of specialization (σ=0.1) while Sp5 is the most generalist (σ=1.5). Species have the same cumulative growth rate along the environmental gradient (i.e., the same areas under the performance curves) regardless of their species-specific σ. Adapted from Buchi & Vuilleumier, 2014.

The term on the right of eq. 3 models the effects of density-dependent competition on population size at the intraspecific and interspecific levels. $\alpha_{intra}$ represents the per capita effects of species *i* on itself whereas $\alpha_{inter}$ is the per capita effect of all other species on the local performance of *i.* Here we assumed stabilizing competition in which $\alpha_{intra}$ > $\alpha_{inter}$. This assumption is relevant because stabilizing competition favors coexistence by increasing the chances of locally rare species to keep positive population growth when locally dominant species have reached equilibrium at high abundances (i.e., the so-called “invasibility criterium” for coexistence Chesson 2000, Grainger et al. 2019). By assuming stabilizing competition, we increased the chances of species with different life-history strategies to coexist in suitable habitats and, consequently, persist in the metacommunity (Thompson et al. 2020). We acknowledge that competition types other than stabilizing (e.g., equalizing: $\alpha_{intra}$ = $\alpha_{inter}$, destabilizing: $\alpha_{intra}$ < $\alpha_{inter}$) may be important to metacommunity dynamics, but evaluating their influence on the way landscapes and species pools are related is beyond the scope of the present study (but see Thompson et al. 2020, Wisnoski and Shoemaker 2022). Nevertheless, our simulations can be easily modified to different levels of intraspecific and interspecific competition as well as to incorporate distinct types of competition. Across all simulations $\alpha_{intra}$ and $\alpha_{inter}$ were set to 1/400 and 1/800 (minimum values that allowed for species regional persistence at high abundance on pre trials), respectively.

We incorporated the influence of ecological drift on local birth and survival by drawing the final local abundance of species *i* from a Poisson distribution (eq. 2) whose mean is given by the deterministic influence of abiotic density-dependent and biotic density-independent selection on population dynamics (following Shoemaker et al. 2020).

Individuals able to persist in the local community after within-patch selection and drift at time *t* could then disperse. To align our framework with recent developments in metacommunity ecology (e.g., Wisnoski et al. 2019, Wisnoski and Shoemaker 2022), we modeled two types of dispersal: spatial and temporal (Buoro and Carlson 2014). Here we define temporal dispersal as any type of physiological (e.g., diapause, dormancy) and/or behavioral strategies (e.g., hiding in refugia) that buffer local extinctions by allowing individuals to escape from temporally unsuitable biotic and abiotic local conditions at the costs of neither reproducing nor consuming available resources. This was operationalized in our simulations by temporally removing individuals from local communities and allowing them to return to the same patch at different moments in the future (see below). Temporal dispersal is relevant because, akin to spatial dispersal, it promotes local and regional coexistence when local abiotic and biotic conditions favor competing species in different periods (i.e., via temporal storage effects, Chesson 2000, Wisnoski and Shoemaker 2022). Therefore, dispersal in space and time can be understood as alternative risk-spreading strategies that can maximize species persistence in metacommunities under varying levels of spatial and temporal environmental heterogeneity (Buoro and Carlson 2014; Holyoak et al. 2020).

The total number of emigrants of species *i* leaving site *j* in time *t* ($E_{i,j,t,total}$) is given by binomial trials whose size is equal to the outcomes of within-patch dynamics (first term of eq. 2) and whose probability of success is given by the species-specific dispersal ability (*η*). Species with higher *η* are more likely to emigrate (i.e., produce more propagules) than individuals of species with lower *η*. To further investigate the effects of spatial and temporal dispersal on the model outcomes, we modeled different scenarios wherein species would be more or less likely to undergo either type of dispersal. This was done by manipulating the values of the parameter *DS* (Dispersal Strategy). *DS* is the probability of success in binomial trials that determine the number of emigrants in $E_{i,j,t,total}$ that would undergo temporal dispersal $(E_{i,j,t,time}$). It follows that the number of spatial emigrants $(E_{i,j,t,space}$) is then given by $E_{ijt, total}$- $E_{ijt,time}$. We modeled three different scenarios. In the “Equal Scenario” species had the same probability of emigrating through either spatial or temporal dispersal (*DS*=*0.5*), whereas in the “Mainly temporal dispersal” and ”Mainly spatial dispersal” scenarios, *DS* was set as very high (0.99) and very low (0.01), respectively, to all species.

The total number of immigrants of species *i* arriving at site *j* in time *t* $(I_{i,j,t,total}$) was given by the sum of spatial ${(I}_{i,j,t,space}$) and temporal $(I_{i,j,t,time}$) immigrants. $I_{i,j,t,space}$ was determined by randomly sampling spatial emigrants ${(E}_{i,j,t,space})$ with a probability equal to the degree of *Connectivity* (given by eq.1) between the focal and neighboring sites. It means that patch *j* is more likely to receive spatial immigrants from closer patches than distant ones. $I_{i,j,t,time}$ is given by the number of individuals engaged in temporal dispersal in site *j* in previous moments in time (*t-x*). Contrary to previous studies that assumed a constant rate of recovery from “dormancy” over time (e.g., Wisnoski et al. 2019), here we considered a more realistic temporal decay in the viability of dormant propagules. To do so, we modeled an exponential decrease in the number of individuals recovering from dormancy with time using an exponential kernel given by *exp(-dt*****Δ****t),* where ***Δ****t* is the difference in time between *t and t-x* (***Δ****t*, min = 1, max =11), and *dt* is the rate of decay. After pre-trials where we tested different values for *dt*, we fixed it at 0.5 because it was the lowest value that allowed species persistence in highly seasonal and disconnected landscapes. Consequently, individuals that, for instance, entered dormancy in time *t* are more likely to recover from dormancy in a nearby future (e.g., *t+1*) than in distant moments in time (e.g., *t+12*).

**Results across different dispersal scenarios**

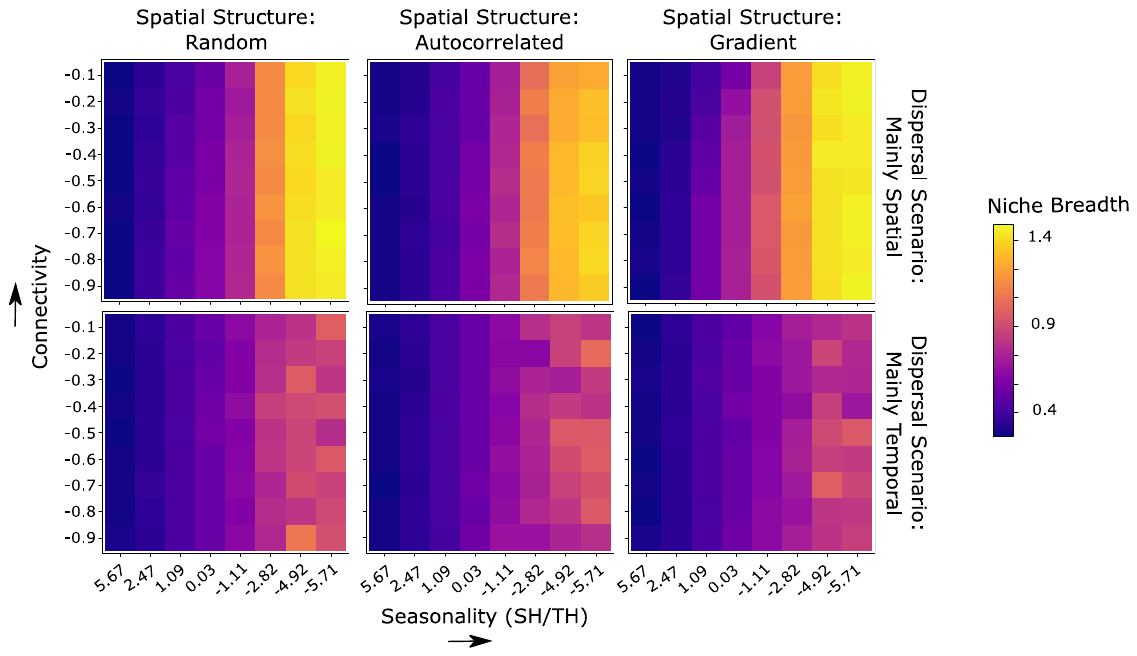

Figure SII: Landscape attributes determine the dominant niche breadth in species pools. Aseasonal (SH/TH > 0) landscapes select for environmental specialists (i.e., narrow niche breadth. Seasonal (SH/TH < 0) landscapes favor the dominance of environmental generalists. Interestingly, when considering metacommunities mainly structured by temporal dispersal, we observe an increase in the persistence of species with a relatively narrow niche breadth in seasonal landscapes. Numerical relationships are depicted in Supp. Inf. tables SIII and SVI

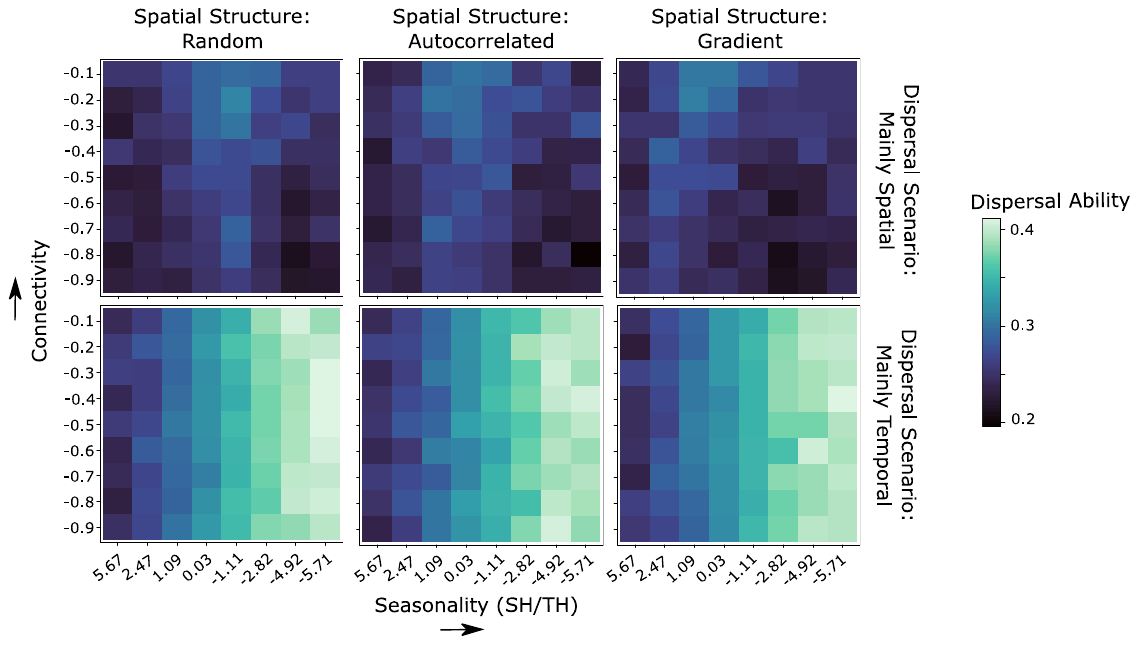

Figure SIII: Landscape attributes determine the dominant dispersal ability in species pools. When spatial dispersal is more frequent than temporal dispersal, dispersal ability was maximized in highly connected landscapes where environmental heterogeneity in space and time were similar (i.e., SH/TH ≈ 0). In contrast, when species were constrained to disperse mainly in time (the “Mostly Temporal” scenario), dispersal ability was maximized at high levels of seasonality (i.e., SH/TH < 0). Numerical relationships are depicted in Supp. Inf. tables III and IV

Table SI Results of Path Analyses fitted considering data on all dispersal scenarios pooled together. “NA” indicates non-applicable parameter estimation. “R.I. Joint All” is the amount of variation in the species data attributable to Environment ∩ Space (MEMs) ∩ Time (AEMs). In bold are the two most relevant pathways (larger standardized estimates) per moderators (Niche breadth and Dispersal ability) and endogenous variables (components of the variation partitioning approach)

| Pathway | Std. Estimate | Std. Error | R-squared |
| --- | --- | --- | --- |
| Niche Breadth <-- SH/TH | **-0.8876** | 0.0182 | 0.79 |
| Niche Breadth <-- Connectivity | -0.0309 | 0.0182 |  |
| Niche Breadth <-- Spatial Structure | **-0.0311** | 0.0182 |  |
| Dispersal Ability <-- SH/TH | **-0.5003** | 0.0338 | 0.28 |
| Dispersal Ability <-- Connectivity | **0.1219** | 0.0338 |  |
| Dispersal Ability <-- Spatial Structure | -0.0228 | 0.0338 |  |
| R.I. Environment <-- SH/TH | 0.0908 | 0.0652 | 0.83 |
| R.I. Environment <-- Connectivity | 0.0839 | 0.0163 |  |
| R.I. Environment <-- Spatial Structure | -0.3020 | 0.0163 |  |
| R.I. Environment <-- Niche Breadth | **-0.665** | 0.0568 |  |
| R.I. Environment<-- Dispersal Ability | **-0.3094** | 0.0305 |  |
| R.I. Space (MEMs) <-- SH/TH | -0.1251 | 0.1154 | 0.48 |
| R.I. Space (MEMs) <-- Connectivity | **-0.3444** | 0.0288 |  |
| R.I. Space (MEMs) <-- Spatial Structure | 0.1318 | 0.0288 |  |
| R.I. Space (MEMs) <-- Niche Breadth | **0.414** | 0.1005 |  |
| R.I. Space (MEMs) <-- Dispersal Ability | 0.1689 | 0.0541 |  |
| R.I. Time (AEMs)<-- SH/TH | **-0.586** | 0.054 | 0.89 |
| R.I. Time (AEMs)<-- Connectivity | 0.1125 | 0.0135 |  |
| R.I. Time (AEMs) <-- Spatial Structure | -0.1109 | 0.0135 |  |
| R.I. Time (AEMs) <--Niche Breadth | **0.2906** | 0.047 |  |
| R.I. Time (AEMs) <-- Dispersal Ability | 0.1644 | 0.0253 |  |
| R.I. Environment ∩ Space (MEMs) <-- SH/TH | -0.2405 | 0.0712 | 0.8 |
| R.I. Environment ∩ Space (MEMs) <-- Connectivity | -0.0124 | 0.0178 |  |
| R.I. Environment ∩ Space (MEMs) <-- Spatial Structure | 0.2371 | 0.0178 |  |
| R.I. Environment ∩ Space (MEMs) <-- Niche Breadth | **-0.9829** | 0.0621 |  |
| R.I. Environment ∩ Space (MEMs) <-- Dispersal Ability | **-0.3681** | 0.0334 |  |
| R.I. Environment ∩ Time (AEMs) <-- SH/TH | 0.2847 | 0.0684 | 0.82 |
| R.I. Environment ∩ Time (AEMs) <-- Connectivity | 0.0326 | 0.0171 |  |
| R.I. Environment ∩ Time (AEMs) <-- Spatial Structure | 0.0876 | 0.0171 |  |
| R.I. Environment ∩ Time (AEMs) <--Niche Breadth | **1.0117** | 0.0596 |  |
| R.I. Environment ∩ Time (AEMs) <-- Dispersal Ability | **0.4752** | 0.032 |  |
| R.I. Space (MEMs) ∩ Time (AEMs) <-- SH/TH | **1.4666** | 0.1217 | 0.42 |
| R.I. Space (MEMs) ∩ Time (AEMs) <-- Connectivity | -0.0564 | 0.0304 |  |
| R.I. Space (MEMs) ∩ Time (AEMs) <-- Spatial Structure | 0.2004 | 0.0304 |  |
| R.I. Space (MEMs) ∩ Time (AEMs) <-- Niche Breadth | **0.8471** | 0.106 |  |
| R.I. Space (MEMs) ∩ Time (AEMs) <-- Dispersal Ability | 0.2706 | 0.057 |  |
| R.I. Joint All <-- SH/TH | **-0.3035** | 0.1521 | 0.1 |
| R.I. Joint All <-- Connectivity | 0.0169 | 0.038 |  |
| R.I. Joint All <-- Spatial Structure | -0.2171 | 0.038 |  |
| R.I. Joint All <-- Niche Breadth | -0.2497 | 0.1325 |  |
| R.I. Joint All <-- Dispersal Ability | **-0.3409** | 0.0713 |  |
| Correlated Errors |  |  |  |
| Niche Breadth<--> Dispersal Ability | -0.7892 | NA | NA |
| R.I. Space (MEMs) <--> R.I. Environment | -0.1255 | NA | NA |
| R.I. Time (AEMs) <--> R.I. Environment | -0.4176 | NA | NA |
| R.I. Environment ∩ Space (MEMs) <--> R.I. Environment | 0.058 | NA | NA |
| R.I. Environment ∩ Time (AEMs) <--> R.I. Environment | -0.3087 | NA | NA |
| R.I. Space (MEMs) ∩ Time (AEMs) <--> R.I. Environment | -0.3943 | NA | NA |
| R.I. Joint All <--> R.I. Environment | -0.2855 | NA | NA |
| R.I. Time (AEMs) <--> R.I. Space (MEMs) | 0.2067 | NA | NA |
| R.I. Environment ∩ Space (MEMs) <--> R.I. Space | -0.387 | NA | NA |
| R.I. Environment ∩ Time (AEMs) <--> R.I. Space | -0.4693 | NA | NA |
| R.I. Space (MEMs) ∩ Time (AEMs) <-->RI.Space | -0.1753 | NA | NA |
| R.I. Joint All <--> R.I. Space | -0.2244 | NA | NA |
| R.I. Environment ∩ Time (AEMs) <--> R.I. Time (AEMs) | 0.0451 | NA | NA |
| R.I. Space (MEMs) ∩ Time (AEMs) <--> R.I. Time | -0.2935 | NA | NA |
| R.I. Joint All <--> R.I. Time (AEMs) | 0.5094 | NA | NA |
| R.I. Environment ∩ Time (AEMs) <--> R.I. Environment ∩ Space (MEMs) | -0.6875 | NA | NA |
| R.I. Space (MEMs) ∩ Time (AEMs) <--> R.I. Environment ∩ Space (MEMs) | -0.3975 | NA | NA |
| R.I. Joint All <--> R.I. Environment ∩ Space (MEMs) | -0.3803 | NA | NA |
| R.I. Space (MEMs) ∩ Time (AEMs) <--> R.I. Environment ∩ Time (AEMs) | 0.5846 | NA | NA |
| R.I. Joint All <--> R.I. Space (MEMs) ∩ Time (AEMs) | 0.0545 | NA | NA |

Table SII Results of Path Analyses fitted considering the Equal scenario. “NA” indicates non-applicable parameter estimation. “R.I. Joint All” is the amount of variation in the species data attributable to. Environment ∩ Space (MEMs) ∩ Time (AEMs). In bold are the two most relevant pathways (larger standardized estimates) per moderators (Niche breadth and Dispersal ability) and endogenous variables (components of the variation partitioning approach)

| Pathway | Std. Estimate | Std. Error | R-squared |
| --- | --- | --- | --- |
| Niche Breadth <-- SH/TH | **-0.9551** | 0.0213 | 0.91 |
| Niche Breadth <-- Connectivity | -0.044 | 0.0213 |  |
| Niche Breadth <-- Spatial Strucure | **-0.0357** | 0.0213 |  |
| Dispersal Ability <-- SH/TH | **-0.8014** | 0.0225 | 0.69 |
| Dispersal Ability <-- Connectivity | **0.227** | 0.0225 |  |
| Dispersal Ability <-- Spatial Strucure | -0.0114 | 0.0225 |  |
| R.I. Environment <-- SH/TH | -0.1066 | 0.0906 | 0.91 |
| R.I. Environment <-- Connectivity | 0.0078 | 0.0209 |  |
| R.I. Environment <-- Spatial Strucure | **-0.3412** | 0.0196 |  |
| R.I. Environment <-- Niche Breadth | **-1.0356** | 0.0699 |  |
| R.I. Environment<-- Dispersal Ability | 0.0664 | 0.0662 |  |
| R.I. Space (MEMs) <-- SH/TH | 0.1354 | 0.2248 | 0.53 |
| R.I. Space (MEMs) <-- Connectivity | **-0.4432** | 0.052 |  |
| R.I. Space (MEMs) <-- Spatial Strucure | 0.174 | 0.0486 |  |
| R.I. Space (MEMs) <-- Niche Breadth | **0.5053** | 0.1736 |  |
| R.I. Space (MEMs) <-- Dispersal Ability | 0.2624 | 0.1643 |  |
| R.I. Time (AEMs)<-- SH/TH | **-0.3312** | 0.0885 | 0.92 |
| R.I. Time (AEMs)<-- Connectivity | 0.1897 | 0.0205 |  |
| R.I. Time (AEMs) <-- Spatial Strucure | -0.1307 | 0.0191 |  |
| R.I. Time (AEMs) <--Niche Breadth | **0.628** | 0.0684 |  |
| R.I. Time (AEMs) <-- Dispersal Ability | -0.013 | 0.0647 |  |
| R.I. Environment ∩ Space (MEMs) <-- SH/TH | **-0.4309** | 0.124 | 0.86 |
| R.I. Environment ∩ Space (MEMs) <-- Connectivity | -0.0784 | 0.0287 |  |
| R.I. Environment ∩ Space (MEMs) <-- Spatial Strucure | 0.2196 | 0.0268 |  |
| R.I. Environment ∩ Space (MEMs) <-- Niche Breadth | **-1.293** | 0.0957 |  |
| R.I. Environment ∩ Space (MEMs) <-- Dispersal Ability | -0.0124 | 0.0906 |  |
| R.I. Environment ∩ Time (AEMs) <-- SH/TH | **0.4961** | 0.1002 | 0.9 |
| R.I. Environment ∩ Time (AEMs) <-- Connectivity | 0.1064 | 0.0232 |  |
| R.I. Environment ∩ Time (AEMs) <-- Spatial Strucure | 0.0957 | 0.0217 |  |
| R.I. Environment ∩ Time (AEMs) <--Niche Breadth | **1.2949** | 0.0774 |  |
| R.I. Environment ∩ Time (AEMs) <-- Dispersal Ability | 0.166 | 0.0732 |  |
| R.I. Space (MEMs) ∩ Time (AEMs) <-- SH/TH | **1.4316** | 0.2024 | 0.5 |
| R.I. Space (MEMs) ∩ Time (AEMs) <-- Connectivity | 0.0082 | 0.0468 |  |
| R.I. Space (MEMs) ∩ Time (AEMs) <-- Spatial Strucure | 0.2915 | 0.0438 |  |
| R.I. Space (MEMs) ∩ Time (AEMs) <-- Niche Breadth | **1.0485** | 0.1562 |  |
| R.I. Space (MEMs) ∩ Time (AEMs) <-- Dispersal Ability | -0.1613 | 0.1479 |  |
| R.I. Joint All <-- SH/TH | **-0.6459** | 0.2549 | 0.35 |
| R.I. Joint All <-- Connectivity | 0.1774 | 0.0589 |  |
| R.I. Joint All <-- Spatial Strucure | -0.2539 | 0.0551 |  |
| R.I. Joint All <-- Niche Breadth | -0.0782 | 0.1968 |  |
| R.I. Joint All <-- Dispersal Ability | **-0.9339** | 0.1862 |  |
| Correlated Errors |  |  |  |
| Niche Breadth<--> Dispersal Ability | -0.4485 | NA | NA |
| R.I. Space (MEMs) <--> R.I. Environment | -0.2327 | NA | NA |
| R.I. Time (AEMs) <--> R.I. Environment | -0.2005 | NA | NA |
| R.I. Environment ∩ Space (MEMs) <--> R.I. Environment | -0.5315 | NA | NA |
| R.I. Environment ∩ Time (AEMs) <--> R.I. Environment | 0.2409 | NA | NA |
| R.I. Space (MEMs) ∩ Time (AEMs) <--> R.I. Environment | -0.3842 | NA | NA |
| R.I. Joint All <--> R.I. Environment | -0.4287 | NA | NA |
| R.I. Time (AEMs) <--> R.I. Space (MEMs) | 0.2485 | NA | NA |
| R.I. Environment ∩ Space (MEMs) <--> R.I. Space | -0.559 | NA | NA |
| R.I. Environment ∩ Time (AEMs) <--> R.I. Space | -0.4387 | NA | NA |
| R.I. Space (MEMs) ∩ Time (AEMs) <-->RI.Space | -0.1381 | NA | NA |
| R.I. Joint All <--> R.I. Space | 0.0712 | NA | NA |
| R.I. Environment ∩ Time (AEMs) <--> R.I. Time (AEMs) | -0.1717 | NA | NA |
| R.I. Space (MEMs) ∩ Time (AEMs) <--> R.I. Time | -0.3601 | NA | NA |
| R.I. Joint All <--> R.I. Time (AEMs) | 0.3121 | NA | NA |
| R.I. Environment ∩ Time (AEMs) <--> R.I. Environment ∩ Space (MEMs) | -0.62 | NA | NA |
| R.I. Space (MEMs) ∩ Time (AEMs) <--> R.I. Environment ∩ Space (MEMs) | -0.5673 | NA | NA |
| R.I. Joint All <--> R.I. Environment ∩ Space (MEMs) | -0.1733 | NA | NA |
| R.I. Space (MEMs) ∩ Time (AEMs) <--> R.I. Environment ∩ Time (AEMs) | 0.6593 | NA | NA |
| R.I. Joint All <--> R.I. Space (MEMs) ∩ Time (AEMs) | -0.0548 | NA | NA |

Table SIII Results of Path Analyses fitted considering the “Mostly spatial” scenario. “NA” indicates non-applicable parameter estimation. “R.I. Joint All” is the amount of variation in the species data attributable to Environment ∩ Space (MEMs) ∩ Time (AEMs). In bold are the two most relevant pathways (larger standardized estimates) per moderators (Niche breadth and Dispersal ability) and endogenous variables (components of the variation partitioning approach)

| Pathway | Std. Estimate | Std. Error | R-squared |
| --- | --- | --- | --- |
| Niche Breadth <-- SH/TH | **-0.9594** | 0.0226 | 0.92 |
| Niche Breadth <-- Connectivity | **-0.0298** | 0.0225 |  |
| Niche Breadth <-- Spatial Structure | -0.0023 | 0.0226 |  |
| Dispersal Ability <-- SH/TH | **0.071** | 0.0258 | 0.46 |
| Dispersal Ability <-- Connectivity | **0.5063** | 0.0148 |  |
| Dispersal Ability <-- Spatial Structure | -0.0197 | 0.018 |  |
| R.I. Environment <-- SH/TH | 0.1061 | 0.0898 | 0.91 |
| R.I. Environment <-- Connectivity | -0.0051 | 0.024 |  |
| R.I. Environment <-- Spatial Structure | **-0.2663** | 0.0198 |  |
| R.I. Environment <-- Niche Breadth | **-0.7439** | 0.0794 |  |
| R.I. Environment<-- Dispersal Ability | 0.1896 | 0.0695 |  |
| R.I. Space (MEMs) <-- SH/TH | -0.1579 | 0.2158 | 0.51 |
| R.I. Space (MEMs) <-- Connectivity | **-0.4226** | 0.0577 |  |
| R.I. Space (MEMs) <-- Spatial Structure | 0.0204 | 0.0475 |  |
| R.I. Space (MEMs) <-- Niche Breadth | **0.4189** | 0.1908 |  |
| R.I. Space (MEMs) <-- Dispersal Ability | 0.0139 | 0.167 |  |
| R.I. Time (AEMs)<-- SH/TH | **-0.2635** | 0.0916 | 0.89 |
| R.I. Time (AEMs)<-- Connectivity | 0.2482 | 0.0245 |  |
| R.I. Time (AEMs) <-- Spatial Structure | -0.1388 | 0.0202 |  |
| R.I. Time (AEMs) <--Niche Breadth | **0.6416** | 0.081 |  |
| R.I. Time (AEMs) <-- Dispersal Ability | -0.1196 | 0.0709 |  |
| R.I. Environment ∩ Space (MEMs) <-- SH/TH | **-0.5518** | 0.1288 | 0.83 |
| R.I. Environment ∩ Space (MEMs) <-- Connectivity | -0.0466 | 0.0345 |  |
| R.I. Environment ∩ Space (MEMs) <-- Spatial Strucure | 0.3076 | 0.0283 |  |
| R.I. Environment ∩ Space (MEMs) <-- Niche Breadth | **-1.4026** | 0.1139 |  |
| R.I. Environment ∩ Space (MEMs) <-- Dispersal Ability | -0.0703 | 0.0997 |  |
| R.I. Environment ∩ Time (AEMs) <-- SH/TH | **0.4617** | 0.1201 | 0.85 |
| R.I. Environment ∩ Time (AEMs) <-- Connectivity | 0.0096 | 0.0321 |  |
| R.I. Environment ∩ Time (AEMs) <-- Spatial Structure | 0.024 | 0.0264 |  |
| R.I. Environment ∩ Time (AEMs) <--Niche Breadth | **1.3963** | 0.1062 |  |
| R.I. Environment ∩ Time (AEMs) <-- Dispersal Ability | 0.1912 | 0.0929 |  |
| R.I. Space (MEMs) ∩ Time (AEMs) <-- SH/TH | **1.4685** | 0.1641 | 0.31 |
| R.I. Space (MEMs) ∩ Time (AEMs) <-- Connectivity | -0.1843 | 0.0439 |  |
| R.I. Space (MEMs) ∩ Time (AEMs) <-- Spatial Strucure | 0.3442 | 0.0361 |  |
| R.I. Space (MEMs) ∩ Time (AEMs) <-- Niche Breadth | **1.1932** | 0.1451 |  |
| R.I. Space (MEMs) ∩ Time (AEMs) <-- Dispersal Ability | 0.1504 | 0.127 |  |
| R.I. Joint All <-- SH/TH | -0.0807 | 0.3 | 0.34 |
| R.I. Joint All <-- Connectivity | 0.3189 | 0.0803 |  |
| R.I. Joint All <-- Spatial Strucure | -0.1707 | 0.066 |  |
| R.I. Joint All <-- Niche Breadth | **-0.4103** | 0.2653 |  |
| R.I. Joint All <-- Dispersal Ability | **-0.6547** | 0.2322 |  |
| Correlated Errors |  |  |  |
| Niche Breadth<--> Dispersal Ability | -0.6538 | NA | NA |
| R.I. Space (MEMs) <--> R.I. Environment | -0.0362 | NA | NA |
| R.I. Time (AEMs) <--> R.I. Environment | -0.2486 | NA | NA |
| R.I. Environment ∩ Space (MEMs) <--> R.I. Environment | -0.2255 | NA | NA |
| R.I. Environment ∩ Time (AEMs) <--> R.I. Environment | 0.0639 | NA | NA |
| R.I. Space (MEMs) ∩ Time (AEMs) <--> R.I. Environment | -0.3977 | NA | NA |
| R.I. Joint All <--> R.I. Environment | -0.5913 | NA | NA |
| R.I. Time (AEMs) <--> R.I. Space (MEMs) | 0.3263 | NA | NA |
| R.I. Environment ∩ Space (MEMs) <--> R.I. Space | -0.5183 | NA | NA |
| R.I. Environment ∩ Time (AEMs) <--> R.I. Space | -0.3609 | NA | NA |
| R.I. Space (MEMs) ∩ Time (AEMs) <-->RI.Space | -0.3008 | NA | NA |
| R.I. Joint All <--> R.I. Space | -0.1541 | NA | NA |
| R.I. Environment ∩ Time (AEMs) <--> R.I. Time (AEMs) | -0.0218 | NA | NA |
| R.I. Space (MEMs) ∩ Time (AEMs) <--> R.I. Time | -0.2049 | NA | NA |
| R.I. Joint All <--> R.I. Time (AEMs) | 0.474 | NA | NA |
| R.I. Environment ∩ Time (AEMs) <--> R.I. Environment ∩ Space (MEMs) | -0.6659 | NA | NA |
| R.I. Space (MEMs) ∩ Time (AEMs) <--> R.I. Environment ∩ Space (MEMs) | -0.5368 | NA | NA |
| R.I. Joint All <--> R.I. Environment ∩ Space (MEMs) | -0.3417 | NA | NA |
| R.I. Space (MEMs) ∩ Time (AEMs) <--> R.I. Environment ∩ Time (AEMs) | 0.6927 | NA | NA |
| R.I. Joint All <--> R.I. Space (MEMs) ∩ Time (AEMs) | -0.0912 | NA | NA |

Table S-IV Results of Path Analyses fitted considering the “Mostly temporal” scenario. “NA” indicates non-applicable parameter estimation. “R.I. Joint All” is the amount of variation in the species data attributable to Environment ∩ Space (MEMs) ∩ Time (AEMs). In bold are the two most relevant pathways (larger standardized estimates) per moderators (Niche breadth and Dispersal ability) and endogenous variables (components of the variation partitioning approach)

| Pathway | Std. Estimate | Std. Error | R-squared |
| --- | --- | --- | --- |
| Niche Breadth <-- SH/TH | **-0.9597** | 0.0226 | 0.92 |
| Niche Breadth <-- Connectivity | -0.0204 | 0.0225 |  |
| Niche Breadth <-- Spatial Structure | **-0.0944** | 0.0226 |  |
| Dispersal Ability <-- SH/TH | **-0.9739** | 0.0258 | 0.95 |
| Dispersal Ability <-- Connectivity | 0.0092 | 0.0258 |  |
| Dispersal Ability <-- Spatial Structure | **-0.0489** | 0.0258 |  |
| R.I. Environment <-- SH/TH | -0.2863 | 0.0898 | 0.87 |
| R.I. Environment <-- Connectivity | 0.019 | 0.024 |  |
| R.I. Environment <-- Spatial Structure | -0.3604 | 0.0198 |  |
| R.I. Environment <-- Niche Breadth | **-0.5859** | 0.0794 |  |
| R.I. Environment<-- Dispersal Ability | **-0.5833** | 0.0695 |  |
| R.I. Space (MEMs) <-- SH/TH | **1.1738** | 0.2158 | 0.61 |
| R.I. Space (MEMs) <-- Connectivity | -0.186 | 0.0577 |  |
| R.I. Space (MEMs) <-- Spatial Structure | 0.2789 | 0.0475 |  |
| R.I. Space (MEMs) <-- Niche Breadth | 0.3517 | 0.1908 |  |
| R.I. Space (MEMs) <-- Dispersal Ability | **1.4976** | 0.167 |  |
| R.I. Time (AEMs)<-- SH/TH | **-0.9475** | 0.0916 | 0.96 |
| R.I. Time (AEMs)<-- Connectivity | 0.0566 | 0.0245 |  |
| R.I. Time (AEMs) <-- Spatial Structure | -0.0933 | 0.0202 |  |
| R.I. Time (AEMs) <--Niche Breadth | -0.1084 | 0.081 |  |
| R.I. Time (AEMs) <-- Dispersal Ability | **0.1361** | 0.0709 |  |
| R.I. Environment ∩ Space (MEMs) <-- SH/TH | -0.3803 | 0.1288 | 0.84 |
| R.I. Environment ∩ Space (MEMs) <-- Connectivity | -0.0267 | 0.0345 |  |
| R.I. Environment ∩ Space (MEMs) <-- Spatial Structure | 0.1555 | 0.0283 |  |
| R.I. Environment ∩ Space (MEMs) <-- Niche Breadth | **-0.7419** | 0.1139 |  |
| R.I. Environment ∩ Space (MEMs) <-- Dispersal Ability | **-0.5294** | 0.0997 |  |
| R.I. Environment ∩ Time (AEMs) <-- SH/TH | **0.7365** | 0.1201 | 0.81 |
| R.I. Environment ∩ Time (AEMs) <-- Connectivity | 0.0409 | 0.0321 |  |
| R.I. Environment ∩ Time (AEMs) <-- Spatial Structure | 0.1929 | 0.0264 |  |
| R.I. Environment ∩ Time (AEMs) <--Niche Breadth | **0.9544** | 0.1062 |  |
| R.I. Environment ∩ Time (AEMs) <-- Dispersal Ability | 0.66 | 0.0929 |  |
| R.I. Space (MEMs) ∩ Time (AEMs) <-- SH/TH | **1.078** | 0.1641 | 0.68 |
| R.I. Space (MEMs) ∩ Time (AEMs) <-- Connectivity | 0.029 | 0.0439 |  |
| R.I. Space (MEMs) ∩ Time (AEMs) <-- Spatial Structure | 0.1037 | 0.0361 |  |
| R.I. Space (MEMs) ∩ Time (AEMs) <-- Niche Breadth | **0.8709** | 0.1451 |  |
| R.I. Space (MEMs) ∩ Time (AEMs) <-- Dispersal Ability | -0.5634 | 0.127 |  |
| R.I. Joint All <-- SH/TH | **-0.8534** | 0.3 | 0.17 |
| R.I. Joint All <-- Connectivity | -0.0279 | 0.0803 |  |
| R.I. Joint All <-- Spatial Structure | -0.2935 | 0.066 |  |
| R.I. Joint All <-- Niche Breadth | -0.1682 | 0.2653 |  |
| R.I. Joint All <-- Dispersal Ability | **-0.3901** | 0.2322 |  |
| Correlated Errors |  |  |  |
| Niche Breadth<--> Dispersal Ability | 0.3462 | NA | NA |
| R.I. Space (MEMs) <--> R.I. Environment | 0.1498 | NA | NA |
| R.I. Time (AEMs) <--> R.I. Environment | -0.5352 | NA | NA |
| R.I. Environment ∩ Space (MEMs) <--> R.I. Environment | -0.1065 | NA | NA |
| R.I. Environment ∩ Time (AEMs) <--> R.I. Environment | -0.4334 | NA | NA |
| R.I. Space (MEMs) ∩ Time (AEMs) <--> R.I. Environment | -0.1236 | NA | NA |
| R.I. Joint All <--> R.I. Environment | -0.1193 | NA | NA |
| R.I. Time (AEMs) <--> R.I. Space (MEMs) | 0.4642 | NA | NA |
| R.I. Environment ∩ Space (MEMs) <--> R.I. Space | -0.6688 | NA | NA |
| R.I. Environment ∩ Time (AEMs) <--> R.I. Space | -0.7772 | NA | NA |
| R.I. Space (MEMs) ∩ Time (AEMs) <-->RI.Space | -0.2832 | NA | NA |
| R.I. Joint All <--> R.I. Space | -0.0047 | NA | NA |
| R.I. Environment ∩ Time (AEMs) <--> R.I. Time (AEMs) | 0.0276 | NA | NA |
| R.I. Space (MEMs) ∩ Time (AEMs) <--> R.I. Time | -0.0298 | NA | NA |
| R.I. Joint All <--> R.I. Time (AEMs) | 0.1037 | NA | NA |
| R.I. Environment ∩ Time (AEMs) <--> R.I. Environment ∩ Space (MEMs) | -0.6743 | NA | NA |
| R.I. Space (MEMs) ∩ Time (AEMs) <--> R.I. Environment ∩ Space (MEMs) | -0.6418 | NA | NA |
| R.I. Joint All <--> R.I. Environment ∩ Space (MEMs) | -0.336 | NA | NA |
| R.I. Space (MEMs) ∩ Time (AEMs) <--> R.I. Environment ∩ Time (AEMs) | 0.7693 | NA | NA |
| R.I. Joint All <--> R.I. Space (MEMs) ∩ Time (AEMs) | 0.1545 | NA | NA |

**Empirical Support: Data information**

**Moth metacommunity in the MTC (data from Maicher et al. 2019)**

Data publicly available taken from <https://doi.org/10.5061/dryad.mgqnk98vr>

Data cleaning:

Our analyses considered only moths that were collected using light traps (i.e., we filtered the data to remove butterflies and fruit-feeding moths). As such, out of the 1,099 species of Lepidoptera collected, our final analyses only considered 561 species (see MTC_Metacommunity.txt file). Information about the sampling design can be found in Maicher et al. (2019).

Species were sampled across 7 elevations (30, 300, 650, 1100, 1450, 1850, 2200m.a.s.l) in three different moments of their growing season between the years of 2014-2017. According to Table 1 in Maicher et al. (2019), the “Wet to Dry” season comprises the months of October to December; the Dry season goes from January to February; the Dry to Wet season goes from March to May. We used this information to extract the climate variables from (Karger et al. 2017) for each sample site between 2014-2017. The climate data used in further analysis (Climate_moth_MTC.txt) represent the monthly values of mean, maximum, minimum temperature, and precipitation per elevation averaged across 2014-2017 .

**Moth metacommunity in the AEF (**Miller and Jones 2005)

Data publicly available taken from <https://doi.org/10.6073/pasta/0cebe58bcc514e2bbf890ee7b2ea21c1>

Data cleaning:

Moths were sampled in the AEF from 1994 to 2004. However, only in 2004, the year we used in our analyses, were moths systematically sampled in the totality of the growing season (from May to October). Moths were sampled in 12 plots (each with one or two sub-plots) along an elevational gradient ranging from 400 to 1400m.a.s.l. We summed all moths collected across sub-plots in a given moment in time to define species abundance in a community. Monthly climate data was extracted from the average latitude and longitude of subplots in a plot (Climate_moth_AEF.txt). No moths were collected in the plots in high elevations in May, so they were removed. In total, 367 species were considered in the final analyses (AEJ_Metacommunity.txt)

**References**

Büchi, L., and S. Vuilleumier. 2014. Coexistence of specialist and generalist species is shaped by dispersal and environmental factors. American Naturalist 183:612–624.

Buoro, M., and S. M. Carlson. 2014. Life-history syndromes: Integrating dispersal through space and time. Ecology Letters 17:756–767.

Chesson, P. 2000. Mechanisms of maintenance os species diversity. Annual Review of Ecology and Systematics 31:343–366.

Fournier, B., N. Mouquet, M. A. Leibold, and D. Gravel. 2017. An integrative framework of coexistence mechanisms in competitive metacommunities. Ecography 40:630–641.

Grainger, T. N., J. M. Levine, and B. Gilbert. 2019. The Invasion Criterion: A Common Currency for Ecological Research. Trends in Ecology and Evolution 34:925–935.

Holyoak, M., T. Caspi, and L. W. Redosh. 2020. Integrating Disturbance, Seasonality, Multi-Year Temporal Dynamics, and Dormancy Into the Dynamics and Conservation of Metacommunities. Frontiers in Ecology and Evolution 8:1–17.

Karger, D. N., O. Conrad, J. Böhner, T. Kawohl, H. Kreft, R. W. Soria-Auza, N. E. Zimmermann, et al. 2017. Climatologies at high resolution for the earth’s land surface areas. Scientific Data 4:170122.

Maicher, V., S. Sáfián, M. Murkwe, S. Delabye, Ł. Przybyłowicz, P. Potocký, I. N. Kobe, et al. 2019. Seasonal shifts of biodiversity patterns and species’ elevation ranges of butterflies and moths along a complete rainforest elevational gradient on Mount Cameroon. Journal of Biogeography 47:342–354.

Miller, J., and J. A. Jones. 2005. Spatial and temporal distribution and abundance of moths in the Andrews Experimental Forest, 1994 to 2008. H. J. Andrews Experimental Forest. Forest Science Data Bank, Corvallis. http://andlter.forestry.oregonstate.edu/data/abstract.aspx?dbcode=SA015.

Shoemaker, L. G., L. L. Sullivan, I. Donohue, J. S. Cabral, R. J. Williams, M. M. Mayfield, J. M. Chase, et al. 2020. Integrating the underlying structure of stochasticity into community ecology. Ecology 101:1–17.

Thompson, P. L., L. M. Guzman, L. De Meester, Z. Horváth, R. Ptacnik, B. Vanschoenwinkel, D. S. Viana, et al. 2020. A process-based metacommunity framework linking local and regional scale community ecology. Ecology Letters 23:1314–1329.

Wisnoski, N. I., M. A. Leibold, and J. T. Lennon. 2019. Dormancy in metacommunities. American Naturalist 194:135–151.

Wisnoski, N. I., and L. G. Shoemaker. 2022. Seed banks alter metacommunity diversity: The interactive effects of competition, dispersal and dormancy. Ecology Letters 25:740–753.
