## Supplementary material for "The geography of metacommunities: landscape characteristics drive geographic variation in the assembly process through selecting species pool attributes": Code and Data to run simulations and generate figures: Analyzing simulated data-Heat Maps and Path models.R

```
library(gplots)
library(lattice)
library(viridisLite)
library(viridis)
library(RColorBrewer)
library(latticeExtra)
library(ggplot2)
library(piecewiseSEM)

Data_path_model_total<-read.table("Metacommunity_Data_Simulations.txt",header=T,sep="\t", dec=".")
dim(Data_path_model_total)

### Heat Maps: Figures 2, SII, SIII#####

#Sub-setting data based on the dispersal scenario and Landscape structure

#Spatial_structure<- 1 =Random; 2= Autocorrelated;3 = Linear gradient

Traits_Equal_Random<-subset(Data_path_model_total,Spat.Struc == '1' & Dispersal_Mode=="Equal")[,c(3,4,5,6,8)]
Traits_Equal_Random_summ<-aggregate(Traits_Equal_Random[,c(2:4)],by = list(Landscape=Traits_Equal_Random$Landscape_factor,Connectivity=as.factor(Traits_Equal_Random$Connectivity)),mean)

Traits_Equal_Auto<-subset(Data_path_model_total,Spat.Struc == '2' & Dispersal_Mode=="Equal")[,c(3,4,5,6,8)]
Traits_Equal_Auto_summ<-aggregate(Traits_Equal_Auto[,c(2:4)],by = list(Landscape=Traits_Equal_Auto$Landscape_factor,Connectivity=as.factor(Traits_Equal_Auto$Connectivity)),mean)

Traits_Equal_Gradient<-subset(Data_path_model_total,Spat.Struc == '3' & Dispersal_Mode=="Equal")[,c(3,4,5,6,8)]
Traits_Equal_Gradient_summ<-aggregate(Traits_Equal_Gradient[,c(2:4)],by = list(Landscape=Traits_Equal_Gradient$Landscape_factor,Connectivity=as.factor(Traits_Equal_Gradient$Connectivity)),mean)


Traits_Spatial_Random<-subset(Data_path_model_total,Spat.Struc == '1' & Dispersal_Mode=="Spatial")[,c(3,4,5,6,8)]
Traits_Spatial_Random_summ<-aggregate(Traits_Spatial_Random[,c(2:4)],by = list(Landscape=Traits_Spatial_Random$Landscape_factor,Connectivity=as.factor(Traits_Spatial_Random$Connectivity)),mean)

Traits_Spatial_Auto<-subset(Data_path_model_total,Spat.Struc == '2' & Dispersal_Mode=="Spatial")[,c(3,4,5,6,8)]
Traits_Spatial_Auto_summ<-aggregate(Traits_Spatial_Auto[,c(2:4)],by = list(Landscape=Traits_Spatial_Auto$Landscape_factor,Connectivity=as.factor(Traits_Spatial_Auto$Connectivity)),mean)


Traits_Spatial_Gradient<-subset(Data_path_model_total,Spat.Struc == '3' & Dispersal_Mode=="Spatial")[,c(3,4,5,6,8)]
Traits_Spatial_Gradient_summ<-aggregate(Traits_Spatial_Gradient[,c(2:4)],by = list(Landscape=Traits_Spatial_Gradient$Landscape_factor,Connectivity=as.factor(Traits_Spatial_Gradient$Connectivity)),mean)


Traits_Temporal_Random<-subset(Data_path_model_total,Spat.Struc == '1' & Dispersal_Mode=="Temporal")[,c(3,4,5,6,8)]
Traits_Temporal_Random_summ<-aggregate(Traits_Temporal_Random[,c(2:4)],by = list(Landscape=Traits_Temporal_Random$Landscape_factor,Connectivity=as.factor(Traits_Temporal_Random$Connectivity)),mean)

Traits_Temporal_Auto<-subset(Data_path_model_total,Spat.Struc == '2' & Dispersal_Mode=="Temporal")[,c(3,4,5,6,8)]
Traits_Temporal_Auto_summ<-aggregate(Traits_Temporal_Auto[,c(2:4)],by = list(Landscape=Traits_Temporal_Auto$Landscape_factor,Connectivity=as.factor(Traits_Temporal_Auto$Connectivity)),mean)


Traits_Temporal_Gradient<-subset(Data_path_model_total,Spat.Struc == '3' & Dispersal_Mode=="Temporal")[,c(3,4,5,6,8)]
Traits_Temporal_Gradient_summ<-aggregate(Traits_Temporal_Gradient[,c(2:4)],by = list(Landscape=Traits_Temporal_Gradient$Landscape_factor,Connectivity=as.factor(Traits_Temporal_Gradient$Connectivity)),mean)

#Average values of SH/TH across all simulations. This will be in the x axis of the heatmaps
Landscape_levels<-aggregate(Data_path_model_total$SH_TH,by = list(Landscape=Data_path_model_total$Landscape_factor),mean)


Trait_Across_Simulations_list<-list(Traits_Equal_Random_summ,
                                    Traits_Equal_Auto_summ,
                                    Traits_Equal_Gradient_summ,
                                    Traits_Spatial_Random_summ,
                                    Traits_Spatial_Auto_summ,
                                    Traits_Spatial_Gradient_summ,
                                    Traits_Temporal_Random_summ,
                                    Traits_Temporal_Auto_summ,
                                    Traits_Temporal_Gradient_summ)

# Matrices in this list have mean NB values across levels of connectivity (collumns) and SH/TH (rows)

Heat_map_list_NB_Global_list<-lapply(Trait_Across_Simulations_list,function(x)as.matrix(reshape(x[,c("Landscape","Connectivity","NB")], 
                  idvar = "Landscape", timevar = "Connectivity", direction = "wide")[,-1]))

Heat_map_list_NB_Global_list<-lapply(Heat_map_list_NB_Global_list,function(x){dimnames(x)<-list(c(as.character(round(Landscape_levels[,2],2))),c(seq(-0.9,-0.1,length.out=9)));x})

Heat_map_list_DA_Global_list<-lapply(Trait_Across_Simulations_list,function(x)as.matrix(reshape(x[,c("Landscape","Connectivity","DA")], 
                                                                                                idvar = "Landscape", timevar = "Connectivity", direction = "wide")[,-1]))
# Matrices in this list have mean DA values across levels of connectivity (collumns) and SH/TH (rows)
Heat_map_list_DA_Global_list<-lapply(Heat_map_list_DA_Global_list,function(x){dimnames(x)<-list(c(as.character(round(Landscape_levels[,2],2))),c(seq(-0.9,-0.1,length.out=9)));x})


coul=plasma(100)
my.min <- min(unlist(Heat_map_list_NB_Global_list))

my.max <- max(unlist(Heat_map_list_NB_Global_list))

my.at <- seq(my.min, my.max, length.out=length(coul))
my.ckey <- list(at=my.at, col=coul)

###Niche breadth Equal Dispersal scenario (Figure 3 Main Manuscript )
NB_Equal_Random_Heat<-levelplot(Heat_map_list_NB_Global_list[[1]],col.regions=coul,at=my.at, colorkey=my.ckey,xlab="", ylab="",scales = list(tck = c(1,0),cex=1.5));NB_Equal_Random_Heat
NB_Equal_Auto_Heat<-levelplot(Heat_map_list_NB_Global_list[[2]],col.regions=coul,at=my.at, colorkey=my.ckey,xlab="", ylab="",scales = list(tck = c(1,0),cex=1.5));NB_Equal_Auto_Heat
NB_Equal_Grad_Heat<-levelplot(Heat_map_list_NB_Global_list[[3]],col.regions=coul,at=my.at, colorkey=my.ckey,xlab="", ylab="",scales = list(tck = c(1,0),cex=1.5));NB_Equal_Grad_Heat

### Niche breadth Mainly Spatial Dispersal scenario (Figure SII Supp.Material )

NB_Spatial_Random_Heat<-levelplot(Heat_map_list_NB_Global_list[[4]],col.regions=coul,at=my.at, colorkey=my.ckey,xlab="", ylab="",scales = list(tck = c(1,0),cex=1.5));NB_Spatial_Random_Heat
NB_Spatial_Auto_Heat<-levelplot(Heat_map_list_NB_Global_list[[5]],col.regions=coul,at=my.at, colorkey=my.ckey,xlab="", ylab="",scales = list(tck = c(1,0),cex=1.5));NB_Spatial_Auto_Heat
NB_Spatial_Grad_Heat<-levelplot(Heat_map_list_NB_Global_list[[6]],col.regions=coul,at=my.at, colorkey=my.ckey,xlab="", ylab="",scales = list(tck = c(1,0),cex=1.5));NB_Spatial_Grad_Heat

### Niche breadth Mainly Temporal Dispersal scenario (Figure SII Supp.Material )
NB_Temporal_Random_Heat<-levelplot(Heat_map_list_NB_Global_list[[7]],col.regions=coul,at=my.at, colorkey=my.ckey,xlab="", ylab="",scales = list(tck = c(1,0),cex=1.5));NB_Temporal_Random_Heat
NB_Temporal_Auto_Heat<-levelplot(Heat_map_list_NB_Global_list[[8]],col.regions=coul,at=my.at, colorkey=my.ckey,xlab="", ylab="",scales = list(tck = c(1,0),cex=1.5));NB_Temporal_Auto_Heat
NB_Temporal_Grad_Heat<-levelplot(Heat_map_list_NB_Global_list[[9]],col.regions=coul,at=my.at, colorkey=my.ckey,xlab="", ylab="",scales = list(tck = c(1,0),cex=1.5));NB_Temporal_Grad_Heat


coul=mako(100)
my.min <- min(unlist(Heat_map_list_DA_Global_list))

my.max <- max(unlist(Heat_map_list_DA_Global_list))

my.at <- seq(my.min, my.max, length.out=length(coul))
my.ckey <- list(at=my.at, col=coul)

###Dispersal ability Equal Dispersal scenario (Figure 3 Main Manuscript )
DA_Equal_Random_Heat<-levelplot(Heat_map_list_DA_Global_list[[1]],col.regions=coul,at=my.at, colorkey=my.ckey,xlab="", ylab="",scales = list(tck = c(1,0),cex=1.5));DA_Equal_Random_Heat
DA_Equal_Auto_Heat<-levelplot(Heat_map_list_DA_Global_list[[2]],col.regions=coul,at=my.at, colorkey=my.ckey,xlab="", ylab="",scales = list(tck = c(1,0),cex=1.5));DA_Equal_Auto_Heat
DA_Equal_Grad_Heat<-levelplot(Heat_map_list_DA_Global_list[[3]],col.regions=coul,at=my.at, colorkey=my.ckey,xlab="", ylab="",scales = list(tck = c(1,0),cex=1.5));DA_Equal_Grad_Heat

###Dispersal ability Spatial Dispersal scenario (Figure SII Supp.Material )

DA_Spatial_Random_Heat<-levelplot(Heat_map_list_DA_Global_list[[4]],col.regions=coul,at=my.at, colorkey=my.ckey,xlab="", ylab="",scales = list(tck = c(1,0),cex=1.5));DA_Spatial_Random_Heat
DA_Spatial_Auto_Heat<-levelplot(Heat_map_list_DA_Global_list[[5]],col.regions=coul,at=my.at, colorkey=my.ckey,xlab="", ylab="",scales = list(tck = c(1,0),cex=1.5));DA_Spatial_Auto_Heat
DA_Spatial_Grad_Heat<-levelplot(Heat_map_list_DA_Global_list[[6]],col.regions=coul,at=my.at, colorkey=my.ckey,xlab="", ylab="",scales = list(tck = c(1,0),cex=1.5));DA_Spatial_Grad_Heat

###Dispersal ability Temporal Dispersal scenario (Figure SII Supp.Material )

DA_Temporal_Random_Heat<-levelplot(Heat_map_list_DA_Global_list[[7]],col.regions=coul,at=my.at, colorkey=my.ckey,xlab="", ylab="",scales = list(tck = c(1,0),cex=1.5));DA_Temporal_Random_Heat
DA_Temporal_Auto_Heat<-levelplot(Heat_map_list_DA_Global_list[[8]],col.regions=coul,at=my.at, colorkey=my.ckey,xlab="", ylab="",scales = list(tck = c(1,0),cex=1.5));DA_Temporal_Auto_Heat
DA_Temporal_Grad_Heat<-levelplot(Heat_map_list_DA_Global_list[[9]],col.regions=coul,at=my.at, colorkey=my.ckey,xlab="", ylab="",scales = list(tck = c(1,0),cex=1.5));DA_Temporal_Grad_Heat


####### Path Analyses (Figure 4  and tables SI-SIV) #####


###  Running the pathmodels based on the whole dataset leads to overfitting. We will run on averages instead 
Data_path_model_summed<-aggregate(Data_path_model_total[,c("SH_TH","NB","DA","Spat.Struc",
                                                           "RI.Env","RI.Space","RI.Time",
                                                           "RI.Env_Space","RI.Env_Time",
                                                           "RI.Space_Time","RI.Joint.All") ],by = list(Landscape=Data_path_model_total$Landscape_factor,Connectivity=Data_path_model_total$Connectivity,Spatial_Structure=Data_path_model_total$Spat.Struc,Dispersal_mode=Data_path_model_total$Dispersal_Mode),mean)

Data_path_model_summed<-Data_path_model_summed[,-1]# Remove Landscape factor variable


Data_path_model_summed_scaled<-data.frame(scale(Data_path_model_summed[,-3]))# Scaling
Data_path_model_summed_scaled$Dispersal_Mode<-Data_path_model_summed$Dispersal_Mode

### Fitting Path models 

#Global Model 

Model_psem_RI<-psem(
  lm(NB~SH_TH+Connectivity+Spat.Struc,Data_path_model_summed_scaled),
  lm(DA~SH_TH+Connectivity+Spat.Struc,Data_path_model_summed_scaled),
  
  lm(RI.Env~SH_TH+Connectivity+Spat.Struc+NB+DA,Data_path_model_summed_scaled),
  lm(RI.Space~SH_TH+Connectivity+Spat.Struc+NB+DA,Data_path_model_summed_scaled),
  lm(RI.Time~SH_TH+Connectivity+Spat.Struc+NB+DA,Data_path_model_summed_scaled),
  lm(RI.Env_Space~SH_TH+Connectivity+Spat.Struc+NB+DA,Data_path_model_summed_scaled),
  lm(RI.Env_Time~SH_TH+Connectivity+Spat.Struc+NB+DA,Data_path_model_summed_scaled),
  lm(RI.Space_Time~SH_TH+Connectivity+Spat.Struc+NB+DA,Data_path_model_summed_scaled),
  lm(RI.Joint.All~SH_TH+Connectivity+Spat.Struc+NB+DA,Data_path_model_summed_scaled),
  
  ##Correlated Errors

  DA%~~%NB,
  RI.Space%~~%RI.Env,
  RI.Time%~~% RI.Env ,
  RI.Env_Space %~~%RI.Env,
  RI.Env_Time%~~% RI.Env,
  RI.Joint.All %~~%RI.Env,
  RI.Time %~~% RI.Space ,
  RI.Env_Space %~~% RI.Space,
  RI.Env_Time %~~% RI.Space,
  RI.Space_Time %~~%RI.Space,
  RI.Joint.All %~~% RI.Space ,
  RI.Env_Space %~~% RI.Time,
  RI.Env_Time %~~% RI.Time,
  RI.Space_Time %~~% RI.Time,
  RI.Joint.All %~~%RI.Time,
  RI.Env_Time %~~% RI.Env_Space,
  RI.Space_Time %~~% RI.Env_Space,
  RI.Joint.All %~~% RI.Env_Space,
  RI.Space_Time %~~% RI.Env_Time,
  RI.Joint.All %~~% RI.Env_Time,
  RI.Joint.All %~~% RI.Space_Time,
  RI.Space_Time %~~%RI.Env
  
)

#Table SI
Global_model<-summary(Model_psem_RI)

#Arrow Width in  figure 4
Size_arrows_fig_4<-data.frame(Global_model$coefficients[,c("Response","Predictor","Std.Error","Std.Estimate")],Size_arrow=Global_model$coefficients[,c("Std.Estimate")]*20)


####Equal 

Equal_Only<-subset(Data_path_model_summed_scaled,Data_path_model_summed_scaled$Dispersal_Mode=="Equal")

Model_psem_RI_Equal_Only<-psem(
  lm(NB~SH_TH+Connectivity+Spat.Struc,Equal_Only),
  lm(DA~SH_TH+Connectivity+Spat.Struc,Equal_Only),
  
  lm(RI.Env~SH_TH+Connectivity+Spat.Struc+NB+DA,Equal_Only),
  lm(RI.Space~SH_TH+Connectivity+Spat.Struc+NB+DA,Equal_Only),
  lm(RI.Time~SH_TH+Connectivity+Spat.Struc+NB+DA,Equal_Only),
  lm(RI.Env_Space~SH_TH+Connectivity+Spat.Struc+NB+DA,Equal_Only),
  lm(RI.Env_Time~SH_TH+Connectivity+Spat.Struc+NB+DA,Equal_Only),
  lm(RI.Space_Time~SH_TH+Connectivity+Spat.Struc+NB+DA,Equal_Only),
  lm(RI.Joint.All~SH_TH+Connectivity+Spat.Struc+NB+DA,Equal_Only),
  
  ##Correlated Errors
  
  DA%~~%NB,
  RI.Space%~~%RI.Env,
  RI.Time%~~% RI.Env ,
  RI.Env_Space %~~%RI.Env,
  RI.Env_Time%~~% RI.Env,
  RI.Joint.All %~~%RI.Env,
  RI.Time %~~% RI.Space ,
  RI.Env_Space %~~% RI.Space,
  RI.Env_Time %~~% RI.Space,
  RI.Space_Time %~~%RI.Space,
  RI.Joint.All %~~% RI.Space ,
  RI.Env_Space %~~% RI.Time,
  RI.Env_Time %~~% RI.Time,
  RI.Space_Time %~~% RI.Time,
  RI.Joint.All %~~%RI.Time,
  RI.Env_Time %~~% RI.Env_Space,
  RI.Space_Time %~~% RI.Env_Space,
  RI.Joint.All %~~% RI.Env_Space,
  RI.Space_Time %~~% RI.Env_Time,
  RI.Joint.All %~~% RI.Env_Time,
  RI.Joint.All %~~% RI.Space_Time,
  RI.Space_Time %~~%RI.Env
)


#Table SII
Equal_coeffs<-summary(Model_psem_RI_Equal_Only)


#Spatial Only
Spatial_Only<-subset(Data_path_model_summed_scaled,Data_path_model_summed_scaled$Dispersal_Mode=="Spatial")


Model_psem_RI_Spatial_Only<-psem(
  lm(NB~SH_TH+Connectivity+Spat.Struc,Spatial_Only),
  lm(DA~SH_TH+Connectivity+Spat.Struc,Spatial_Only),
  
  lm(RI.Env~SH_TH+Connectivity+Spat.Struc+NB+DA,Spatial_Only),
  lm(RI.Space~SH_TH+Connectivity+Spat.Struc+NB+DA,Spatial_Only),
  lm(RI.Time~SH_TH+Connectivity+Spat.Struc+NB+DA,Spatial_Only),
  lm(RI.Env_Space~SH_TH+Connectivity+Spat.Struc+NB+DA,Spatial_Only),
  lm(RI.Env_Time~SH_TH+Connectivity+Spat.Struc+NB+DA,Spatial_Only),
  lm(RI.Space_Time~SH_TH+Connectivity+Spat.Struc+NB+DA,Spatial_Only),
  lm(RI.Joint.All~SH_TH+Connectivity+Spat.Struc+NB+DA,Spatial_Only),
  
  ##Correlated Errors
  
  DA%~~%NB,
  RI.Space%~~%RI.Env,
  RI.Time%~~% RI.Env ,
  RI.Env_Space %~~%RI.Env,
  RI.Env_Time%~~% RI.Env,
  RI.Joint.All %~~%RI.Env,
  RI.Time %~~% RI.Space ,
  RI.Env_Space %~~% RI.Space,
  RI.Env_Time %~~% RI.Space,
  RI.Space_Time %~~%RI.Space,
  RI.Joint.All %~~% RI.Space ,
  RI.Env_Space %~~% RI.Time,
  RI.Env_Time %~~% RI.Time,
  RI.Space_Time %~~% RI.Time,
  RI.Joint.All %~~%RI.Time,
  RI.Env_Time %~~% RI.Env_Space,
  RI.Space_Time %~~% RI.Env_Space,
  RI.Joint.All %~~% RI.Env_Space,
  RI.Space_Time %~~% RI.Env_Time,
  RI.Joint.All %~~% RI.Env_Time,
  RI.Joint.All %~~% RI.Space_Time,
  RI.Space_Time %~~%RI.Env
  
)


#Table SIII
Spatial_coeffs<-summary(Model_psem_RI_Spatial_Only)


#Temporal only
Temporal_Only<-subset(Data_path_model_summed_scaled,Data_path_model_summed_scaled$Dispersal_Mode=="Temporal")

Model_psem_RI_Temporal_Only<-psem(
  lm(NB~SH_TH+Connectivity+Spat.Struc,Temporal_Only),
  lm(DA~SH_TH+Connectivity+Spat.Struc,Temporal_Only),
  
  lm(RI.Env~SH_TH+Connectivity+Spat.Struc+NB+DA,Temporal_Only),
  lm(RI.Space~SH_TH+Connectivity+Spat.Struc+NB+DA,Temporal_Only),
  lm(RI.Time~SH_TH+Connectivity+Spat.Struc+NB+DA,Temporal_Only),
  lm(RI.Env_Space~SH_TH+Connectivity+Spat.Struc+NB+DA,Temporal_Only),
  lm(RI.Env_Time~SH_TH+Connectivity+Spat.Struc+NB+DA,Temporal_Only),
  lm(RI.Space_Time~SH_TH+Connectivity+Spat.Struc+NB+DA,Temporal_Only),
  lm(RI.Joint.All~SH_TH+Connectivity+Spat.Struc+NB+DA,Temporal_Only),
  
  ##Correlated Errors
  
  DA%~~%NB,
  RI.Space%~~%RI.Env,
  RI.Time%~~% RI.Env ,
  RI.Env_Space %~~%RI.Env,
  RI.Env_Time%~~% RI.Env,
  RI.Joint.All %~~%RI.Env,
  RI.Time %~~% RI.Space ,
  RI.Env_Space %~~% RI.Space,
  RI.Env_Time %~~% RI.Space,
  RI.Space_Time %~~%RI.Space,
  RI.Joint.All %~~% RI.Space ,
  RI.Env_Space %~~% RI.Time,
  RI.Env_Time %~~% RI.Time,
  RI.Space_Time %~~% RI.Time,
  RI.Joint.All %~~%RI.Time,
  RI.Env_Time %~~% RI.Env_Space,
  RI.Space_Time %~~% RI.Env_Space,
  RI.Joint.All %~~% RI.Env_Space,
  RI.Space_Time %~~% RI.Env_Time,
  RI.Joint.All %~~% RI.Env_Time,
  RI.Joint.All %~~% RI.Space_Time,
  RI.Space_Time %~~%RI.Env
  
)

#Table SIV
Temporal_coeffs<-summary(Model_psem_RI_Temporal_Only)
```
