## Supplementary material for "The geography of metacommunities: landscape characteristics drive geographic variation in the assembly process through selecting species pool attributes": Code and Data to run simulations and generate figures: Empirical data-Analyses -Figure 5.R

```
#Empirical support : The assembly of moth metacommunities in a tropical and temperate mountain. 

####### Panel A: Landscape spatiotemporal heterogeneity#####
library(reshape2)
library(gplots)
library(viridis)
library(lattice)
library(plyr)


#Andrews Experimental Forest -USA (AEF)
#Climate data AEF. Names of rows are given by month_plotID number. Variables are given by the average across all plot sub-units.
#See more information in https://andlter.forestry.oregonstate.edu/data/abstract.aspx?dbcode=SA015

Data_Climate_AEJ<-read.table("Climate_moth_AEF.txt",sep="\t", dec=".",header = T)

Data_Climate_AEJ$Time<- dplyr::recode_factor(Data_Climate_AEJ$Time,"T1"="Jan","T2"="Feb","T3"="Mar","T4"="April","T5"="May","T6"="Jun",
                                             "T7"="Jul","T8"="Ago","T9"="Sep","T10"="Oct","T11"="Nov","T12"="Dec")

rownames(Data_Climate_AEJ)<-paste0(Data_Climate_AEJ$Time,"_",Data_Climate_AEJ$Sites)


Data_Climate_AEJ$Precipitation<-log(Data_Climate_AEJ$Precipitation)#Make it normal

PCA_env<-princomp(scale(Data_Climate_AEJ[,3:6]),cor=T)
summary(PCA_env)
Data_Climate_AEJ$PC1<-PCA_env$scores[,1]

Matrix_heat_AEJ<-dcast(Data_Climate_AEJ[,c(1,2,10)],Sites~Time,value.var = "PC1")
rownames(Matrix_heat_AEJ)<-Matrix_heat_AEJ$Sites
Matrix_heat_AEJ<-Matrix_heat_AEJ[as.character(Data_Climate_AEJ$Sites[1:12]),-1]
row.names(Matrix_heat_AEJ)<-paste(Data_Climate_AEJ$Elev[1:12],"m.a.s.l.")

Time_var<-mean(apply(Matrix_heat_AEJ,1,var))
Space_var<-mean(apply(Matrix_heat_AEJ,2,var))
Spatiotemporal_Heterogeneity_AEJ<-round(log(Space_var/Time_var),2)#SH/TH

coul=turbo(100)
my.min <- min(Matrix_heat_AEJ)

my.max <- max(Matrix_heat_AEJ)

my.at <- seq(my.min, my.max, length.out=length(coul))
my.ckey <- list(at=my.at, col=coul)

levelplot(as.matrix(t(Matrix_heat_AEJ)),aspect=1,col.regions=coul,at=my.at, colorkey=my.ckey,xlab="", ylab="",main=paste("SH/TH =",as.character(Spatiotemporal_Heterogeneity_AEJ)),scales = list(x=list(rot=45),tck = c(1,0),cex=2))

# Mount Cameroon 
#Average climatic variables per month between 2014 - 2017. (See details in Maicher et al. 2019)

Data_Climate_MTC<-read.table("Climate_moth_MTC.txt",header=T,sep="\t", dec=".") 

Data_Climate_MTC$Precipitation<-log(Data_Climate_MTC$Precipitation)#Make it Normal

PCA_env<-princomp(scale(Data_Climate_MTC[,3:6]),cor=T)
summary(PCA_env)
Data_Climate_MTC$PC1<-PCA_env$scores[,1]

Matrix_heat<-dcast(Data_Climate_MTC[,c(1,2,9)],Sites~Time,value.var = "PC1")
rownames(Matrix_heat)<-Matrix_heat$Sites
Matrix_heat<-Matrix_heat[c("S1","S2","S3","S4","S5","S6","S7"),
                         c("T1","T2","T3","T4","T5","T6","T7","T8","T9","T10","T11","T12")]

colnames(Matrix_heat)<-c("Jan","Feb","Mar","Apr","May","Jun","Jul","Aug","Sep","Oct","Nov","Dec")
rownames(Matrix_heat)<-c("30 m.a.s.l.","350 m.a.s.l.","650 m.a.s.l.","1100 m.a.s.l.","1450 m.a.s.l.","1850 m.a.s.l","2200 m.a.s.l")


Time_var<-mean(apply(Matrix_heat,1,var))
Space_var<-mean(apply(Matrix_heat,2,var))
Spatiotemporal_Heterogeneity_MTC=round(log(Space_var/Time_var),2)

coul=turbo(100)
my.min <- min(Matrix_heat)

my.max <- max(Matrix_heat)

my.at <- seq(my.min, my.max, length.out=length(coul))
my.ckey <- list(at=my.at, col=coul)

levelplot(as.matrix(t(Matrix_heat)),aspect=1,col.regions=coul,at=my.at, colorkey=my.ckey,xlab="", ylab="",main=paste("SH/TH =",as.character(Spatiotemporal_Heterogeneity_MTC)),scales = list(x=list(rot=45),tck = c(1,0),cex=2))

####Panel B: Niche Breadth#####


library(dplyr)
library(reshape2)
library(plyr)
library(ade4)
library(vegan)
library(ggplot2)
library(adespatial)
#Here we pooled the metacommunities and climates so niche breadths are calculate in the same parameter space and, consequently, are comparable. 

#Species x sites distribution of both mountains pooled together. From lines 1-70 is the metacommunity in AEJ. 
#The rest is the metacommunity in MTC


#Metacommunity in the AEF
AEJ_MC<-read.table("AEJ_Metacommunity.txt",header=T,sep="\t", dec=".",row.names = "Sites")
names(AEJ_MC) <- gsub("\\.", " ", names(AEJ_MC))#Fixing names
Species_pool_AEJ<-colnames(AEJ_MC)

#Subseting the climate data to contain only the months and sites where the metacommunity was sampled
Env_AEJ<-Data_Climate_AEJ[rownames(AEJ_MC),3:8]


#Metacommunity in the MTC
MTC_MC<-read.table("MTC_Metacommunity.txt",header=T,sep="\t", dec=".",row.names = "Sites")
names(MTC_MC) <- gsub("\\.", " ", names(MTC_MC))#Fixing names
Species_pool_MTC<-colnames(MTC_MC)


#Calculating average climate variables per-season, following information of when data was collected  (See details in Maicher et al. 2019, table 1)
#"WTD"= Wet to dry season
#"Dry" = "Dry season
# "DTW" = Dry to wet season
Wet_to_Dry<-subset(Data_Climate_MTC,Data_Climate_MTC$Time%in%c("T10","T11","T12"))
Wet_to_Dry<-aggregate(. ~ Sites, data=Wet_to_Dry[,-2], FUN=mean)
Wet_to_Dry$Time<-rep("WTD",nrow(Wet_to_Dry))

Dry<-subset(Data_Climate_MTC,Data_Climate_MTC$Time%in%c("T1","T2","T3"))
Dry<-aggregate(. ~ Sites, data=Dry[,-2], FUN=mean)
Dry$Time<-rep("DRY",nrow(Dry))

Dry_to_Wet<-subset(Data_Climate_MTC,Data_Climate_MTC$Time%in%c("T4","T5","T6"))
Dry_to_Wet<-aggregate(. ~ Sites, data=Dry_to_Wet[,-2], FUN=mean)
Dry_to_Wet$Time<-rep("DTW",nrow(Dry_to_Wet))

Env_MTC<-rbind(Wet_to_Dry,Dry,Dry_to_Wet)
rownames(Env_MTC)<-paste0(Env_MTC$Time,"_",Env_MTC$Sites)
Env_MTC<-Env_MTC[,c(-1,-8)]
Env_MTC<-Env_MTC[,c(1,3,2,4,5,6)]

sites_ID<-c(rownames(AEJ_MC),rownames(MTC_MC))

#Below we pooled the metacommunities and climates so niche breadths are calculate in the same multivariate space and, consequently, are comparable. 

#Species x sites distribution of both mountains pooled together. From lines 1-70 is the metacommunity in AEJ. 
#The rest is the metacommunity in MTC


MC_AEJ_MTC_list<-list(AEJ_MC,MTC_MC)


allNms <- unique(unlist(lapply(MC_AEJ_MTC_list, function(x)names(x))))
Moths_data_pooled<-do.call(rbind,
                           lapply(MC_AEJ_MTC_list,
                                  function(x) data.frame(c(x, sapply(setdiff(allNms, names(x)),
                                                                     function(y) NA)))))

Moths_data_pooled[is.na(Moths_data_pooled)==T]<-0
rownames(Moths_data_pooled)<-sites_ID


names(Moths_data_pooled) <- gsub("\\.", " ", names(Moths_data_pooled))#Fixing names

names(Env_MTC)<-names(Env_AEJ)
Moths_Env_data_pooled<-rbind(Env_AEJ,Env_MTC)
Moths_Env_data_pooled<-Moths_Env_data_pooled[rownames(Moths_data_pooled),]


#Calculating Niche breadth based on Doledec (2000). This index estimates species-specific climatic tolerance 
#(i.e., niche breadth) based on the dispersal of samples that contain the target species in the multivariate climatic space.
dudi.pca.env <- dudi.pca(Moths_Env_data_pooled[,1:4], scale = TRUE, scan = FALSE, nf=4)
niche.species <- niche(dudi.pca.env, Y = Moths_data_pooled, scann = FALSE)
d.niche <- data.frame(
  niche.pos=niche.species$li,
  as.data.frame(niche.param(niche.species))
)


d.niche$Location<-c(rep("AEF",length(Species_pool_AEJ)),rep("MTC",length(Species_pool_MTC)))

Tol_subset_Moth_AEJ<-subset(d.niche,d.niche$Location%in%c("AEF"))
rownames(Tol_subset_Moth_AEJ) <- gsub("\\.", " ", rownames(Tol_subset_Moth_AEJ))


#Selecting the 33% dominant in each meta
Dominant_species_AEJ<-sort(colSums(AEJ_MC),decreasing = T)[1:round(length(Species_pool_AEJ)/3)]

Tol_subset_Moth_Top_30_AEJ<-Tol_subset_Moth_AEJ[names(Dominant_species_AEJ),]

Tol_subset_Moth_MTC<-subset(d.niche,d.niche$Location%in%c("MTC"))
rownames(Tol_subset_Moth_MTC) <- gsub("\\.", " ", rownames(Tol_subset_Moth_MTC))


Dominant_species_MTC<-sort(colSums(MTC_MC),decreasing = T)[1:round(length(Species_pool_MTC)/3)]

Tol_subset_Moth_Top_30_MTC<-Tol_subset_Moth_MTC[names(Dominant_species_MTC),]


Tol_top_30_only_moths<-rbind(Tol_subset_Moth_Top_30_AEJ,Tol_subset_Moth_Top_30_MTC)


#### Figure 5 Panel B
p <- ggplot(Tol_top_30_only_moths, aes(x=Location, y= Tol,fill=Location)) +
  geom_violin(scale = "width",trim = FALSE,adjust = 1,alpha=0.5)+
  geom_boxplot(outlier.shape = NA,width=0.1,fill="gray",alpha=0.5)+
  ylab("Climate Tolerance")+
  xlab("")+
  scale_fill_brewer(palette="Dark2") + 
  theme_classic()+
  theme(axis.title.y = element_text(size = 14),
        axis.text.x = element_text(size = 14),
        legend.position = "none");p


#### Panel C: Variation and Beeta-diversity Partitioning####
library(stringr)
library(vegan)
library(reshape2)
library(ggplot2)
library(gplots)
library(gdm)
library(adespatial)


#Functions

# #Variation Partitioning Using GDMs. Community= Species by site by time matrix.
#Coords = Spatial Coodinates
#SpatialMEM= Positive Spatial MEMs
#TemporalAEM= Temporal AEM
Var_part_GDMS4<-function(Community,Env_obs,Coords,SpatialMEM,TemporalAEM){
  
  X2<-Community
  
  Env_obs<-Env_obs[rownames(X2),]
  
  Env_X<-as.matrix(Env_obs)
  
  Coord_X<-Coords[rownames(X2),]
  names<-row.names(X2)
  Spatial_MEM_GDM<-SpatialMEM[rownames(X2),]
  Temporal_AEM_GDM<-TemporalAEM[rownames(X2),]
  
  
  data_gdm_total<-data.frame(names,Lat=Coord_X$Lat,Long=Coord_X$Long,Env_X,Spatial_MEM_GDM,Temporal_AEM_GDM)# Formating Env data to  GDMs
  data_gdm_Spaitaltemporal_Data<-data.frame(names,Lat=Coord_X$Lat,Long=Coord_X$Long,Env_X=rep(1,nrow(Env_X)),Spatial_MEM_GDM,Temporal_AEM_GDM)# Formating Env data to  GDMs
  data_gdm_Only_env<-data.frame(names,Lat=Coord_X$Lat,Long=Coord_X$Long,Env_X)# Formating Env data to  GDMs
  data_gdm_Only_Spatial<-data.frame(names,Lat=Coord_X$Lat,Long=Coord_X$Long,Env_X=rep(1,nrow(Env_X)),Spatial_MEM_GDM)# Formating Env data to  GDMs
  data_gdm_Only_Temporal<-data.frame(names,Lat=Coord_X$Lat,Long=Coord_X$Long,Env_X=rep(1,nrow(Env_X)),Temporal_AEM_GDM)# Formating Env data to  GDMs
  data_gdm_Env_Temporal<-data.frame(names,Lat=Coord_X$Lat,Long=Coord_X$Long,Env_X,Temporal_AEM_GDM)# Formating Env data to  GDMs
  data_gdm_Env_Spatial<-data.frame(names,Lat=Coord_X$Lat,Long=Coord_X$Long,Env_X,Spatial_MEM_GDM)# Formating Env data to  GDMs
  
  
  
  # Dummy env variables nescessary to variaiton partitioning approach
  
  #Dependent variable in the formart nescessary for the GDMs
  X_default<-cbind(Coord_X,X2,names)
  X_matrix<-melt(X_default,id.vars=c("Lat","Long","names"),variable.name = "sppCol")
  
  
  X_gdm2_Full_model= formatsitepair(X_matrix,bioFormat=2,XColumn="Lat",YColumn = "Long",predData =data_gdm_total,sppColumn= "sppCol",siteColumn="names",abundColumn ="value",abundance=T)
  Total_deviation<-gdm(X_gdm2_Full_model,geo = F)
  if(is.null(Total_deviation)==T){
    Explained_Total_deviation<-0
  }else{
    Explained_Total_deviation<-Total_deviation$explained
  }
  
  
  X_gdm2_Only_Env= formatsitepair(X_matrix,bioFormat=2,XColumn="Lat",YColumn = "Long",predData =data_gdm_Only_env,sppColumn= "sppCol",siteColumn="names",abundColumn ="value",abundance=T)
  Only_Env_Deviation<-gdm(X_gdm2_Only_Env,geo = F)
  if(is.null(Only_Env_Deviation)==T){
    Explained_Only_Env_Deviation<-0
  }else{
    Explained_Only_Env_Deviation<-Only_Env_Deviation$explained
  }
  
  
  X_gdm2_Only_Spatial= formatsitepair(X_matrix,bioFormat=2,XColumn="Lat",YColumn = "Long",predData =data_gdm_Only_Spatial,sppColumn= "sppCol",siteColumn="names",abundColumn ="value",abundance=T)
  Only_Spatial_Deviation<-gdm(X_gdm2_Only_Spatial,geo = F)
  if(is.null(Only_Spatial_Deviation)==T){
    Explained_Only_Spatial_Deviation<-0
  }else{
    Explained_Only_Spatial_Deviation<-Only_Spatial_Deviation$explained
  }
  
  
  X_gdm2_Only_Temporal= formatsitepair(X_matrix,bioFormat=2,XColumn="Lat",YColumn = "Long",predData =data_gdm_Only_Temporal,sppColumn= "sppCol",siteColumn="names",abundColumn ="value",abundance=T)
  Only_Temporal_Deviation<-gdm(X_gdm2_Only_Temporal,geo = F)
  if(is.null(Only_Temporal_Deviation)==T){
    Explained_Only_Temporal_Deviation<-0
  }else{
    Explained_Only_Temporal_Deviation<-Only_Temporal_Deviation$explained
  }
  
  X_gdm2_Env_Temporal= formatsitepair(X_matrix,bioFormat=2,XColumn="Lat",YColumn = "Long",predData =data_gdm_Env_Temporal,sppColumn= "sppCol",siteColumn="names",abundColumn ="value",abundance=T)
  Env_Temporal_Deviation<-gdm(X_gdm2_Env_Temporal,geo = F)
  if(is.null(Env_Temporal_Deviation)==T){
    Explained_Env_Temporal_Deviation<-0
  }else{
    Explained_Env_Temporal_Deviation<-Env_Temporal_Deviation$explained
  }
  
  X_gdm2_Env_Spatial= formatsitepair(X_matrix,bioFormat=2,XColumn="Lat",YColumn = "Long",predData =data_gdm_Env_Spatial,sppColumn= "sppCol",siteColumn="names",abundColumn ="value",abundance=T)
  Env_Spatial_Deviation<-gdm(X_gdm2_Env_Spatial,geo = F)
  if(is.null(Env_Spatial_Deviation)==T){
    Explained_Env_Spatial_Deviation<-0
  }else{
    Explained_Env_Spatial_Deviation<-Env_Spatial_Deviation$explained
  }
  
  
  X_gdm2_Spatial_Temporal= formatsitepair(X_matrix,bioFormat=2,XColumn="Lat",YColumn = "Long",predData =data_gdm_Spaitaltemporal_Data,sppColumn= "sppCol",siteColumn="names",abundColumn ="value",abundance=T)
  Spatial_Temporal_Deviation<-gdm(X_gdm2_Spatial_Temporal,geo = F)
  if(is.null(Spatial_Temporal_Deviation)==T){
    Explained_Spatial_Temporal_Deviation<-0
  }else{
    Explained_Spatial_Temporal_Deviation<-Spatial_Temporal_Deviation$explained
  }
  
  
  Env_alone<-Explained_Total_deviation-Explained_Spatial_Temporal_Deviation#a
  Space_alone<-Explained_Total_deviation-Explained_Env_Temporal_Deviation#b
  Time_alone<-Explained_Total_deviation-Explained_Env_Spatial_Deviation#c
  
  a_d<-Explained_Env_Temporal_Deviation-Explained_Only_Temporal_Deviation
  b_e<-Explained_Env_Spatial_Deviation-Explained_Only_Env_Deviation
  c_f<-Explained_Spatial_Temporal_Deviation-Explained_Only_Spatial_Deviation
  
  
  Env_Space_joint<-a_d-Env_alone
  Space_time_joint<-b_e-Space_alone
  Env_time_joint<-c_f-Time_alone
  
  Joint_all<-Explained_Total_deviation-a_d-b_e-c_f
  Results<-c(Env_alone,Space_alone,Time_alone,Env_Space_joint,Env_time_joint,Space_time_joint,Joint_all)/Explained_Total_deviation
  names(Results)<-c("Env Alone","Space Alone","Time Alone","Env_Space_Joint","Env_Time_Joint","Space_Time_Joint","Joint_all")
  return(Results)
  
}


#AEJ

Spatial_MEM_AEJ<-dbmem(Env_AEJ[,c("Lat","Long")],MEM.autocor = c("positive"))
Temporal_AEM_AEJ<-aem.time(6)$aem[rep(1:6, each=12)[-10:-12],]#Six months pooled together are considered in this data + removing the three empty sites in may (May_39,May_3,May_13)

dimnames(Temporal_AEM_AEJ)<-list(rownames(Spatial_MEM_AEJ),paste0("AEM",1:ncol(Temporal_AEM_AEJ)))

Var_part_AEJ<-Var_part_GDMS4(Community = AEJ_MC,Env_obs = Env_AEJ[,1:4],Coords = Env_AEJ[,5:6],SpatialMEM =Spatial_MEM_AEJ,TemporalAEM =Temporal_AEM_AEJ )

#MTC


Spatial_MEM_MTC<-dbmem(Env_MTC[,c("Lat","Long")],MEM.autocor = c("positive"))
Temporal_AEM_MTC<-aem.time(3)$aem[rep(1:3, each=7),]#Three seasons were considered in this data
dimnames(Temporal_AEM_MTC)<-list(rownames(Spatial_MEM_MTC),paste0("AEM",1:ncol(Temporal_AEM_MTC)))

Var_part_MTC<-Var_part_GDMS4(Community = MTC_MC,Env_obs = Env_MTC[,1:4],Coords = Env_MTC[,5:6],SpatialMEM =Spatial_MEM_MTC,TemporalAEM =Temporal_AEM_MTC )


### Figure 5 Panel C
Data_bar_plot<-data.frame(RI=c(Var_part_AEJ,Var_part_MTC),
                          Mechanism=rep(names(Var_part_MTC),2),
                          Region=rep(c("AEF","MTC"),each=7))


Data_bar_plot[Data_bar_plot<0]<-0
Data_bar_plot$Mechanism2<-rep(c("Purely Climate","Purely Space","Purely Time","Shared Climate & Space",
                                "Shared Climate & Time","Shared Space & Time","Shared Climate & Space & Time"),2)

Data_bar_plot2<-subset(Data_bar_plot,Data_bar_plot$Mechanism2%in%c("Purely Climate","Purely Space","Purely Time","Shared Climate & Space",
                                                                   "Shared Climate & Time","Shared Climate & Space & Time"))

#Plot panel C
ggplot(Data_bar_plot2, aes(Region, RI, fill = Mechanism2, group = Mechanism2)) +
  geom_col(position = "fill", width = 0.5, colour = "black",alpha=0.5) +
  scale_fill_manual(breaks=c("Purely Climate","Purely Space","Purely Time","Shared Climate & Space",
                             "Shared Climate & Time","Shared Climate & Space & Time"),values=c("#1B9E77","#D95F02","#7570B3","#66A61E","#E6AB02","#A6761D"))+
  ylab("Proportion of total explained variation")+
  xlab("")+
  geom_area(aes(x = c("AEF" = 1.25, "MTC" = 1.75)[Region]), 
            position = "fill", colour = "grey", alpha = 0.3,
            outline.type = "both")+
  theme_classic()+
  theme(
    axis.title.y = element_text(size = 14),
    axis.text.x = element_text(size = 14),
    legend.title = element_blank(),
  )
```
