## Supplementary material for "The geography of metacommunities: landscape characteristics drive geographic variation in the assembly process through selecting species pool attributes": Code and Data to run simulations and generate figures: README.txt

Title: The geography of metacommunities: landscape characteristics drive geographic variation in the assembly process 
through selecting species pool attributes 

Brief Description 
In this study, we used simulation models to  demonstrate  that a conceptual framework centered on 
the dependency of species pool characteristics on landscape attributes is valuable for understanding 
the geography of metacommunity assembly, i.e., understanding variation in the relative importance of mechanisms 
that assemble different metacommunities distributed along broad-scale ecological gradients or across biogeographic regions. 
We used empirical data on moth metacommunities in a tropical and temperate mountain to provide empirical validation of some of 
the predictions derived from our simulation models.

In this folder one will find:
1- Simulations Notated and Analyses.R- Code to run the simulation models, assess the niche breadth and dispersal ability of dominant species at the end of a simulation round (1200 time-steps), and run variation partitioning in the simulated data. 
2- Metacommunity_Data_Simulations.txt-  data file with the results of simulations performed using code in provided in NAME1 across 20 replicates X  8 spatiotemporal heterogeneity levels (values described in line 593 of Simulations Notated and Analyses.R) x 9 connectivity levels (values described in line 592 of Simulations Notated and Analyses.R) x 3 Spatial levels of spatial structure of env. conditions x 3 dispersal scenarios ("Equal", "Mainly Spatial", "Mainly Temporal"). 
3- Analyzing simulated data-Heat Maps and Path models.R - Code used to analyse data provided in Metacommunity_Data_Simulations.txt and to generate figures 3, figure S2, figure S3, and information needed to create figure 4 and tables SI-SIV
4- MTC_Metacommunity.txt- Data file containing the spatial-temporal distribution of moths in Mount Cameroon (MTC). See  more information in (Maicher et al. 2019)
5- AEJ_Metacommunity.txt - Data file containing the spatial-temporal distribution of moths in Andrew`s Experimental Forest (AEF). See more information in (Miller and Jones 2005).
6- Climate_moth_MTC.txt -Climate data for MTC. Extracted from (Karger et al. 2017)
7- Climate_moth_AEF.txt – Climate data for AEF(Karger et al. 2017)
8- Empirical data-Analyses -Figure 5. R-  Code to clean data provided in items 4-7 and generate figure 5 in main manuscript

Data files description:
Metacommunity_Data_Simulations.txt

Data file resulting from the simulations performed using code in provided in Simulations Notated and Analyses.R
12960 rows x  15 columns

Each row is the result of a simulation iteration. 20 replicates X  8 spatiotemporal heterogeneity levels (values described in line 593 of NAME1) 
x 9 connectivity levels (values described in line 592 Name) x 3 Spatial levels of spatial structure of env. conditions x 
3 dispersal scenarios ("Equal", "Mainly Spatial", "Mainly Temporal")

Column IDs and description:

Spat.Struct = Spatial structure of environmental conditions. Ordinal. 1 =Random; 2= Autocorrelated;3 = Linear gradient.
Dispersal Mode = Dispersal Scenarios. Categorical. “Equal”, “Spatial”, “Temporal”
Connectivity = Landscape Connectivity [-0.9; -0.1]
SH_TH = SH/TH index
NB = Niche breadth (Weighted  mean at the metacommunity scale)
DA = Dispersal ability (Weighted  mean at the metacommunity scale)
DS = Parameter defening species dispersal strategy (0.5 = Equal, 0.99 = Preference to disperse in time, 0.01= Preference to disperse in space) 
Landscape_Factor = Treating SH/TH as ordinal to facilitate further summary analyses
RI. Env = Importance (ranked)  of environment alone in the variation partitioning
RI. Space = Importance (ranked)  of spatial MEMs alone in the variation partitioning
RI. Time = Importance (ranked)    of temporal  AEMs alone in the variation partitioning
RI. Env_Space = Importance (ranked) of covariation between Env. and MEMs in the variation partitioning
RI.Env_Time = Importance (ranked) of covariation between Env. and AEMs in the variation partitioning
RI.Space_Time = Importance (ranked) of covariation between MEMs. and AEMs in the variation partitioning
RI.Joint.All = Joint contribution (ranked) of all variables.

MTC_Metacommunity.txt
Data file containing the spatial-temporal distribution of moths in Mount Cameroon (MTC). 
See  more information in (Maicher et al. 2019). Each row contains the moths sampled in a give elevation different moments of their growing season (Wet to Dry, Dry, Dry to Wet)
21 elev-month x 561 species

AEJ_Metacommunity.txt
Data file containing the spatial-temporal distribution of moths in Andrew`s Experimental Forest (AEF). 
See more information in (Miller and Jones 2005). Each row contains the moths sampled in a give plot (numbers) in a given month of their growing season.
69 elev-month x 367 species

Climate_moth_MTC.txt
Climate data for MTC. Extracted from Karger et al. 2017.
Climate in the sampled sites of the MTC (average between the years 2015-2017)

Column ID

Sites = Sites name
Time = Moment of the year
Mean_Temp = Mean air temperature (Celsius)
Min_Temp = Minimum air temperature (Celsius)
Max_Temp = Maximum air temperature (Celsius)
Precipitation = Precipitation (kg m-2 month1 /100)
Lat = Latitude
Long = Longitude
Elev= Average elevation 

Climate_moth_AEF.txt
Climate data for AEF extracted from Karger et al. 2017.
Climate in the sampled sites of the AEF in the year of 2004

Column ID
Sites = Sites name
Time = Moment of the year
Mean_Temp = Mean air temperature (Celsius)
Min_Temp = Minimum air temperature (Celsius)
Max_Temp = Maximum air temperature (Celsius)
Precipitation = Precipitation (kg m-2 month1 /100)
Lat = Latitude
Long = Longitude
Elev= Average elevation 

 


To validate and illustrate the utility of our conceptual framework, we built a process-based metacommunity model wherein species pools with the same initial distribution of continuous traits (here, ecological specialization and dispersal ability) were allowed to colonize and reach coexistence in landscapes varying in their levels of seasonality, physical connectivity, and spatial structure (autocorrelation) of environmental (habitat) conditions. By assessing the degree of ecological specialization and dispersal ability of the species that could persist and dominate the metacommunity in each type of landscape, we were able to understand how different landscape characteristics select for different dominant life-history strategies observed in species pools.
Following we used variation partitioning to analyse the simulated data. This allowed us to contrast the effects of environmental factors (typically assumed as the result of niche-based processes), spatial (typically assumed as the result of dispersal limitation), and temporal variation (typically assumed as the result of temporal autocorrelation in population dynamics generated by neutral mechanisms) in explaining variation in community composition. . By establishing the links between landscape attributes, species pool characteristics, and associated inferences about community assembly derived from statistical models, we were able to foster theoretical predictions about the variation in the relative importance of mechanisms observed in empirical metacommunities located at different parts of large-scale ecological gradients or in different biogeographic regions
To provide empirical support for some of the theoretical predictions derived from our conceptual framework, we analyzed data on moth metacommunities in a tropical and a temperate mountainous landscape that show distinct patterns of spatial and temporal environmental heterogeneity. As such, we were able to contrast the predictions generated by our conceptual model against the outcomes observed in metacommunities that exhibit significant variation in both species pool characteristics and landscape attributes
