## Supplementary material for "The geography of metacommunities: landscape characteristics drive geographic variation in the assembly process through selecting species pool attributes": Code and Data to run simulations and generate figures: Simulations Notated and Analyses.R

```
# Title: Title: The geography of metacommunities: landscape characteristics drive geographic variation in the assembly 
#process through selecting species pool attributes 

#Running title: The geography of metacommunities

#Metacommunity simulation models

# Simulations followed a modified Beverton-Holt model that determine local population dynamics as 
#a function of dispersal, habitat selection, competition (intraspecific and interspecific) and demographic stochasticity. 
#Species pools with similar initial characteristics were set to colonize 
#and coexist in landscapes with varying levels of physical connectivity, seasonality, and spatial distribution of environmental conditions.
#Dispersal in space is spatially explicit. Dispersal in time (via dormancy) is also considered. Customized code presented here was 
#inspired in Thompson at al 2020, Shoemaker et al 2019, Wisnoski et al 2019, Buchi et al 2014

#Simulations can be summarized into 3 steps (See Supp. Information for more details): 

#Step 1: Within patch dynamics: Species local Performance (local population growth)  is given by functions:
#"habitat_suitability" and "competition". 
#Note that species-specific performances were re-scaled according to their environmental tolerance so that 
#neither generalists or specialists are favored across simulations. (see Supp.Material 1 for details)

#Step 2: Emmigration: Individuals will leave the patch either through spatial or temporal (engaging in dormancy) dispersals. The total number of individuals per species in the dispersal pool 
#is given by species dispersal ability (the success probability in a binomial distribution)

#Step 3: Immigration: Spatial Emmigrants in time "t" will colonize neighboring communities  according to their distance to the original patch (spatially explicit dispersal)
# Individuals that engaged in dormancy in time time "t-1" will recover from dormancy in "t" given a negative exponential kernel 

#Demographic stochasticity (modelled in the form of random draws of known probablity distributions) was considered in all steps described above. 

##################### Essential functions and packages ####################
library(reshape2)
library(gdm)
library(vegan)
library(ggplot2)
library(viridis)
library(WaveletComp)

library(adiv)
library(igraph)
library(adespatial)
library(entropart)
library(ggpubr)
library(lattice)


#Variation Partitioning Using GDMs. Community= Species by site by time matrix.
#Coords = Spatial Coodinates
#SpatialMEM= Positive Spatial MEMs
#TemporalAEM= Temporal AEM
Var_part_GDMS4<-function(Community,Env_obs,Coords,SpatialMEM,TemporalAEM){
  
  X2<-Community
  
  Env_obs<-Env_obs[rownames(X2),]
  
  Env_X<-as.matrix(Env_obs)
  
  Coord_X<-Coords[rownames(X2),]
  names<-row.names(X2)
  Spatial_MEM_GDM<-SpatialMEM[rownames(X2),]
  Temporal_AEM_GDM<-TemporalAEM[rownames(X2),]
  
  
  data_gdm_total<-data.frame(names,Lat=Coord_X$Lat,Long=Coord_X$Long,Env_X,Spatial_MEM_GDM,Temporal_AEM_GDM)# Formating Env data to  GDMs
  data_gdm_Spaitaltemporal_Data<-data.frame(names,Lat=Coord_X$Lat,Long=Coord_X$Long,Env_X=rep(1,nrow(Env_X)),Spatial_MEM_GDM,Temporal_AEM_GDM)# Formating Env data to  GDMs
  data_gdm_Only_env<-data.frame(names,Lat=Coord_X$Lat,Long=Coord_X$Long,Env_X)# Formating Env data to  GDMs
  data_gdm_Only_Spatial<-data.frame(names,Lat=Coord_X$Lat,Long=Coord_X$Long,Env_X=rep(1,nrow(Env_X)),Spatial_MEM_GDM)# Formating Env data to  GDMs
  data_gdm_Only_Temporal<-data.frame(names,Lat=Coord_X$Lat,Long=Coord_X$Long,Env_X=rep(1,nrow(Env_X)),Temporal_AEM_GDM)# Formating Env data to  GDMs
  data_gdm_Env_Temporal<-data.frame(names,Lat=Coord_X$Lat,Long=Coord_X$Long,Env_X,Temporal_AEM_GDM)# Formating Env data to  GDMs
  data_gdm_Env_Spatial<-data.frame(names,Lat=Coord_X$Lat,Long=Coord_X$Long,Env_X,Spatial_MEM_GDM)# Formating Env data to  GDMs
  
  
  
  # Dummy env variables nescessary to variaiton partitioning approach
  
  #Dependent variable in the formart nescessary for the GDMs
  X_default<-cbind(Coord_X,X2,names)
  X_matrix<-melt(X_default,id.vars=c("Lat","Long","names"),variable.name = "sppCol")
  
  
  X_gdm2_Full_model= formatsitepair(X_matrix,bioFormat=2,XColumn="Lat",YColumn = "Long",predData =data_gdm_total,sppColumn= "sppCol",siteColumn="names",abundColumn ="value",abundance=T)
  Total_deviation<-gdm(X_gdm2_Full_model,geo = F)
  if(is.null(Total_deviation)==T){
    Explained_Total_deviation<-0
  }else{
    Explained_Total_deviation<-Total_deviation$explained
  }
  
  
  X_gdm2_Only_Env= formatsitepair(X_matrix,bioFormat=2,XColumn="Lat",YColumn = "Long",predData =data_gdm_Only_env,sppColumn= "sppCol",siteColumn="names",abundColumn ="value",abundance=T)
  Only_Env_Deviation<-gdm(X_gdm2_Only_Env,geo = F)
  if(is.null(Only_Env_Deviation)==T){
    Explained_Only_Env_Deviation<-0
  }else{
    Explained_Only_Env_Deviation<-Only_Env_Deviation$explained
  }
  
  
  X_gdm2_Only_Spatial= formatsitepair(X_matrix,bioFormat=2,XColumn="Lat",YColumn = "Long",predData =data_gdm_Only_Spatial,sppColumn= "sppCol",siteColumn="names",abundColumn ="value",abundance=T)
  Only_Spatial_Deviation<-gdm(X_gdm2_Only_Spatial,geo = F)
  if(is.null(Only_Spatial_Deviation)==T){
    Explained_Only_Spatial_Deviation<-0
  }else{
    Explained_Only_Spatial_Deviation<-Only_Spatial_Deviation$explained
  }
  
  
  X_gdm2_Only_Temporal= formatsitepair(X_matrix,bioFormat=2,XColumn="Lat",YColumn = "Long",predData =data_gdm_Only_Temporal,sppColumn= "sppCol",siteColumn="names",abundColumn ="value",abundance=T)
  Only_Temporal_Deviation<-gdm(X_gdm2_Only_Temporal,geo = F)
  if(is.null(Only_Temporal_Deviation)==T){
    Explained_Only_Temporal_Deviation<-0
  }else{
    Explained_Only_Temporal_Deviation<-Only_Temporal_Deviation$explained
  }
  
  X_gdm2_Env_Temporal= formatsitepair(X_matrix,bioFormat=2,XColumn="Lat",YColumn = "Long",predData =data_gdm_Env_Temporal,sppColumn= "sppCol",siteColumn="names",abundColumn ="value",abundance=T)
  Env_Temporal_Deviation<-gdm(X_gdm2_Env_Temporal,geo = F)
  if(is.null(Env_Temporal_Deviation)==T){
    Explained_Env_Temporal_Deviation<-0
  }else{
    Explained_Env_Temporal_Deviation<-Env_Temporal_Deviation$explained
  }
  
  X_gdm2_Env_Spatial= formatsitepair(X_matrix,bioFormat=2,XColumn="Lat",YColumn = "Long",predData =data_gdm_Env_Spatial,sppColumn= "sppCol",siteColumn="names",abundColumn ="value",abundance=T)
  Env_Spatial_Deviation<-gdm(X_gdm2_Env_Spatial,geo = F)
  if(is.null(Env_Spatial_Deviation)==T){
    Explained_Env_Spatial_Deviation<-0
  }else{
    Explained_Env_Spatial_Deviation<-Env_Spatial_Deviation$explained
  }
  
  
  X_gdm2_Spatial_Temporal= formatsitepair(X_matrix,bioFormat=2,XColumn="Lat",YColumn = "Long",predData =data_gdm_Spaitaltemporal_Data,sppColumn= "sppCol",siteColumn="names",abundColumn ="value",abundance=T)
  Spatial_Temporal_Deviation<-gdm(X_gdm2_Spatial_Temporal,geo = F)
  if(is.null(Spatial_Temporal_Deviation)==T){
    Explained_Spatial_Temporal_Deviation<-0
  }else{
    Explained_Spatial_Temporal_Deviation<-Spatial_Temporal_Deviation$explained
  }
  
  
  Env_alone<-Explained_Total_deviation-Explained_Spatial_Temporal_Deviation#a
  Space_alone<-Explained_Total_deviation-Explained_Env_Temporal_Deviation#b
  Time_alone<-Explained_Total_deviation-Explained_Env_Spatial_Deviation#c
  
  a_d<-Explained_Env_Temporal_Deviation-Explained_Only_Temporal_Deviation
  b_e<-Explained_Env_Spatial_Deviation-Explained_Only_Env_Deviation
  c_f<-Explained_Spatial_Temporal_Deviation-Explained_Only_Spatial_Deviation
  
  
  Env_Space_joint<-a_d-Env_alone
  Space_time_joint<-b_e-Space_alone
  Env_time_joint<-c_f-Time_alone
  
  Joint_all<-Explained_Total_deviation-a_d-b_e-c_f
  Results<-c(Env_alone,Space_alone,Time_alone,Env_Space_joint,Env_time_joint,Space_time_joint,Joint_all)/Explained_Total_deviation
  names(Results)<-c("Env Alone","Space Alone","Time Alone","Env_Space_Joint","Env_Time_Joint","Space_Time_Joint","Joint_all")
  return(Results)
  
}


#Habitat suitaiblity
habitat_suitability<-function(Env_cond,Sp_opt_conditions,nbreadth){
  exp(-((Env_cond - Sp_opt_conditions)^2/(2*nbreadth^2)))
}

#Competition  intra and interspecific interactions (alpha_intra vs. alpha_inter). Returns a matrix giving the degree of intraspecific competition
#per species per site
competition<-function(N,alpha_intra,alpha_inter){
  competition_matrix<-matrix(0,nrow(N),ncol(N))
  for (c in 1:ncol(N)){
    for(d in 1:nrow(N)){
      Intra<-N[d,c]
      Inter<-abs(Intra-sum(N[d,]))
      competition_matrix[d,c]<-  1 / ( 1 + alpha_intra*Intra+alpha_inter*Inter)
    }
  }
  return(competition_matrix)
}


#Function created to estimate spatiotemporal environmental heterogeneity index of simulated landscapes (SH/TH).
#It returns the ratio between the average variance of spatial heterogeneity (i.e., mean variance of columns) 
#and the average variance of temporal heterogeneity (i.e., mean variance of rows). 
Ratio_temporal_spatial_variance<-function(x){
  Temporal_var<-mean(apply(x,1,var))
  Spatial_var<-mean(apply(x,2,var))
  Ratio<-Spatial_var/Temporal_var
  return(Ratio)
}

#Multivaiate normal distribution to simmulate autocorrelated landscapes (See function Landscape_function4)
rmvn <- function(n, mu = 0, V = matrix(1)) {
  p <- length(mu)
  if (any(is.na(match(dim(V), p)))) 
    stop("Dimension problem!")
  D <- chol(V)
  t(matrix(rnorm(n * p), ncol = p) %*% D + rep(mu, rep(n, p)))
}

#Landscape Function

#Function to simulate landscapes with varying levels of SH/TH, Connectivity, and Spatial Autocrrelation.
#N_sites= Number of sites, Seasonality= amplitude of the sinusoidal variation in environment over time. The higher the value assigned to Seasonality, the higher is the seasonal variation in environmental conditions. 
#Time= Total time steps considered
#Spatial_Env_Grad: Type of landscape
#Connectivity: Value that determines the rate at which connectivity (probability of dispersal in space) decays with spatial distance. 

#Returns a list:
#1:A matrix with local environmental conditions (rows) over time (columns).
#2: Site Coordinates
#3: Matrix showing the probability of dispersal among pacthes weighted by their mutual distances (Connectivity)
#4:The value of SH/TH 

#If plot_landscapes = T, two figures are produced: 1 Shows Landscape in first seasonal cycle; 2 Shows landscape in time 1
#Note that only connections above the threshold (1e-4) are shown in the plot. 
Landscape_function4<-function(N_sites,Seasonality,time,Spatial_Env_Grad=c("Gradient","Random","Auto"),Connectivity,plot_landscape,phi){
  #Ensuring that all patches are reachable (i.e.no isolated patches )
  for(ii in 1:10000){
    plot_coord=data.frame(Lat=runif(N_sites,0,60),Long=runif(N_sites,0,60))#Set coordinates
    rownames(plot_coord)<-sites_names
    Prob_dispersal_among_plots<-as.matrix(exp(-Connectivity*dist(plot_coord)))
    
    Prob_dispersal_among_plots[Prob_dispersal_among_plots<1e-4]<-0 #Connectivity threshold.
    if(is_connected(graph_from_adjacency_matrix(Prob_dispersal_among_plots, weighted=TRUE, mode="undirected",diag = F))==T){
      break
    }
    Attempt<-ii
    print(paste("Attempt #",Attempt, ":Generated Landscape is not fully connected. Trying again."))
  } 
  
  #In landscapes of type == "Gradient" environment changes in space as a linear function 
  #of sites coordinates + error. Values are then rescaled to range between 0 and 5
  #In landscapes of type == "Random"  equally spaced environmental values along the gradient [0;5] are randomly assigned them to each patch
  #In landscapes of type "Auto" environmental conditions are modelled using a multivariate normal distribution (mu=0) 
  #whose covariance matrix incorporated the exponential 
  #decay of environmental similarity between pairs of patches as a function of their distances and the constant phi (set at 0.15).
  
  if(Spatial_Env_Grad=="Gradient"){
    Elevation=plot_coord[,1]+plot_coord[,2]+rnorm(N_sites,0,0.1) 
    Elevation_grad=(5-0)/(max(Elevation)-min(Elevation))*(Elevation-max(Elevation))+5
  }else{
    if(Spatial_Env_Grad=="Random"){
      Elevation=runif(N_sites,0,5)
      Elevation_grad<-(5-0)/(max(Elevation)-min(Elevation))*(Elevation-max(Elevation))+5#Rescaling values to 5
    }else{
      
      Elevation<-rmvn(1, rep(0,N_sites), exp(-phi * as.matrix(dist(plot_coord))))
      Elevation_grad<-(5-0)/(max(Elevation)-min(Elevation))*(Elevation-max(Elevation))+5#Rescaling values to 5
      
    }
  }
  
  
  Landscape<-matrix(0,nrow=N_sites,ncol=time)
  for(i in 1:length(Elevation_grad)){
    Landscape[i,]=(Elevation_grad[i]+Seasonality*periodic.series(start.period = 12, length = time+3)+rnorm(time+3,0,.01))[-1:-3]
  }
  
  Landscape_std<-as.matrix(5/(max(Landscape)-min(Landscape))*(Landscape-max(Landscape))+5)
  row.names(Landscape_std)<-sites_names
  colnames(Landscape_std)<-paste0("T",seq (from= 1, to =time))
  
  
  #Generating a figure showing landscape connectivity and SH/TH
  if(plot_landscape==T){
    Plot_land<-Landscape_std[names(sort(rowMeans(Landscape_std))),]
    coul=turbo(100)
    my.at <- seq(min(Plot_land[,1:12]), max(Plot_land[,1:12]), length.out=length(coul))
    my.ckey <- list(at=my.at, col=coul)
    Spatiotemporal_plot<-levelplot(t(Plot_land[,1:12]),col.regions=coul,at=my.at, colorkey=my.ckey,xlab="Time", ylab="Sites",main="Landscape in the first seasonal cycle (T1-T12)",scales = list(tck = c(1,1),cex=1),aspect="fill")
    
    # Spatiotemporal_plot<-heatmap.2(Plot_land[,1:12], scale = "none", col = turbo(100), 
    #           trace = "none", density.info = "none",Colv=F,Rowv=F,dendrogram = 'none')
    # 
    Landscape_data<-data.frame(Lat=plot_coord$Lat,
                               Long=plot_coord$Long,
                               Env=Landscape_std[,1])
    
    Diss_modify_total <- data.frame(t(combn(as.character(rownames(Landscape_data)),2)), as.numeric(as.dist(Prob_dispersal_among_plots)))
    names(Diss_modify_total) <- c("Start", "End", "distance")
    
    
    Connectted_landscapes<-subset(Diss_modify_total,Diss_modify_total$distance>0)
    
    dat2 <- rbind(Landscape_data[c(Connectted_landscapes[,1]),],
                  Landscape_data[c(Connectted_landscapes[,2]),],
                  Landscape_data[c(Connectted_landscapes[,3])])
    dat2$gr<-rep(1:nrow(Connectted_landscapes),times=2)
    
    mid=mean(Landscape_std)
    Plot_Connect_step1<-ggplot(Landscape_data, aes(x = Lat, y = Long,fill=Env)) + 
      geom_line(aes(group = gr),linetype="dashed", data = dat2)+
      ggtitle('Landscape in T1')+
      ylab("Y")+
      xlab("X")+
      #scale_size_continuous(range = c(1, 10))+
      geom_point(shape = 21, size = 5)+
      scale_fill_gradientn(colours = turbo(100),
                           limits=c(0,5))+
      #scale_size_continuous(range = c(0, 1))+
      coord_equal()+
      theme_bw()
    
    #print(Plot_Connect_step1)
    Plots<-ggarrange(Spatiotemporal_plot,Plot_Connect_step1,ncol = 2,nrow = 1)
    print(Plots)
    
  }
  Prob_dispersal_among_plots_final<-as.matrix(Prob_dispersal_among_plots)
  dimnames(Prob_dispersal_among_plots_final)<-list(c(rownames(plot_coord)),
                                                   c(rownames(plot_coord)))
  
  
  Ratio_SH_TH<-log(Ratio_temporal_spatial_variance(Landscape_std))
  
  
  Results<-list(Landscape_std,plot_coord,Prob_dispersal_among_plots_final,Ratio_SH_TH)
  names(Results)<-c("Environment","Coord","Probability of Dispersal Among Plots","log SH/TH")
  return(Results)
  
}

##################

###### Metacommunity Simulation  Model####


#The Metacom_Function returns a list containing:
#1- 1200 site-by-species matrix (one per time step).
#2- A dataframe with the traits of the randomly generated species pool that seeded the landscape.

#It has two arguments: "Landscape_object" which is the landscape created using the Landscape_function4. "
#Dispersal type" is the type of dispersal strategy considered (See main text for details).


Metacom_Function<-function(Landscape_object,Dispersal_type=c("Equal","Temporal_Only","Spatial_Only")){
  
  Env_cond<-Landscape_object[[1]]#Environment Matrix
  Spatiotemporal_env_heterogeneity<-Landscape_object[[4]]#SH/TH
  
  plot_coord<-Landscape_object[[2]]#Plot Coordinates
  Prob_dispersal_among_plots<-Landscape_object[[3]]#Probability of exchanging individuals among plots based on Connectivity and distance
  
  #Modelling the decay of viability of dormant individuals with time. It also 
  # decays following a kernel. It is the same across all species. After pre-trials where we tested different values for dt,
  #we fixed it at 0.5 because it was the lowest value that allowed species persistence in highly seasonal and disconnected landscapes.
  #See more in  Supp. Material I
  Prob_recover_dormancy_over_time<-as.matrix(exp(-0.5*dist(1:12)))[-1,1]# Maximum temporal distance was set to 12 time steps
  
  
  #Simulating Species traits
  Sp_opt_conditions<-runif(N_sp,0,5) # Species enviomnetal optima μ
  
  nbreadth<-runif(N_sp,0.1,2)# Species niche breath
  names(nbreadth)<-sp_names
  
  
  Dispersal_ability<-runif(N_sp,0.01,0.5)# Species dispersal ability η
  names(Dispersal_ability)<-sp_names
  

  # We also wanted to investigate the effects of different dispersal modes in community assembly.
  # The “Equal” scenario assumed that species do not have
  #a preferred way to disperse, and, consequently, are equally capable of engaging in temporal and 
  #spatial dispersal. The “Only Spatial Dispersal�� scenario assumed that the only mode of dispersal is spatial (i.e., temporal dispersal probability was very low). 
  #Finally, the “Only Temporal Dispersal” assumed that the only mode of dispersal is temporal (i.e., spatial dispersal probability was very low). 
  
  #These 3 scenarios were simulated by manipulating the values of the "Dispersal_strategy", vector. 
  #Dispersal_strategy (DS) is the probability of success in a binomial distribution that would determine the number of emigrants of each species 
  #that would engage in temporal dispersal.  
  #In the “Equal” scenario all species had the same probability of emigrating through spatial and temporal  dispersal (DS=0.5), 
  #whereas in Only Temporal and Only spatial and dispersal cases, DT was set to be very high (0.9) and very low (0.01) respectively to all species, respectively. 
  #
  
  
  if(Dispersal_type=="Equal"){
    Dispersal_strategy<-rep(0.5,N_sp) 
  }
  
  if(Dispersal_type=="Temporal_Only"){
    Dispersal_strategy<-rep(0.9,N_sp) 
  }
  
  if(Dispersal_type=="Spatial_Only"){
    Dispersal_strategy<-rep(0.1,N_sp) 
  }
  
  #Creating a data frame to see species traits
  Species_traits<-data.frame(Sp_opt_conditions,nbreadth,Dispersal_ability,Dispersal_strategy)
  #Species_traits$Dispersal_strategy<-sample(rep(Dispersal_mode,each=N_sp/2),replace=F)
  rownames(Species_traits)<-sp_names
  
  
  #Beginning simulations
  #Community matrices (Each will store the metacommunity in different momemnts in time)
  Communities<-lapply(1:time,function(x) matrix(0,N_sites,N_sp,
                                                dimnames = list(c(sites_names),c(sp_names))))
  
  
  
  #Storing the dormant individuals.
  Dormancy_pool<-lapply(1:(time+120),function(x) matrix(0,N_sites,N_sp,
                                                        dimnames = list(c(sites_names),c(sp_names))))## Abundance of innactive species across sites in a specific moment in time
  
  #Species that disperse in space in a specific moment in time
  Emmigration_pool<-lapply(1:time,function(x) matrix(0,N_sites,N_sp,
                                                     dimnames = list(c(sites_names),c(sp_names))))# Species that disperse in space in a specific moment in time
  
  
  
  
  habitat_suitability_space_time<-list()
  
  
  #### Simulation begins here. It may take some a while to finish
  t=1
  while(t <= 1200){
    print(paste0("time_",t))
    
    #habitat suitability for each species at each site in t
    habitat_suitability_each_spxsite<-matrix(NA,N_sites,N_sp,dimnames = list(c(sites_names),c(sp_names)))
    
    for(i in 1:N_sp){
      habitat_suitability_sps<-habitat_suitability(Env_cond[,t],Species_traits[i,"Sp_opt_conditions"],Species_traits[i,"nbreadth"])
      habitat_suitability_each_spxsite[,i]<-habitat_suitability_sps
      
    }
    
    
    ##### Simulate community composition
    
    #In the first 120 time steps we 
    #seeded each patch with species populations randomly drawn from a 
    #Poisson distribution (lambda= 0.5). This allowed all species to have a chance to establish in 
    #all habitat patches and increase in abundance provided that local abiotic and biotic conditions are suitable. 
    
    if(t==1){
      Community_step1<-apply(Communities[[t]],2,function(x)rpois(length(x),0.5))
      row.names(Community_step1)<-row.names(Communities[[t]])
      
    }else{
      if(t%in%2:120){
        
        Seed<-t(sapply(1:nrow(Communities[[t]]),function(x)rpois(N_sp,0.5)))
        Community_step1=Communities[[t-1]]+Seed
        row.names(Community_step1)<-row.names(Communities[[t]])
      }else{
        
        Community_step1=Communities[[t-1]]
        row.names(Community_step1)<-row.names(Communities[[t]])
      }
      
    }  
    
    ####Step 1
    comp<-competition(Community_step1,alpha_intra,alpha_inter)
    
    #Calculating local performances. Term after %*% scales species responses to the environmental gradient, 
    #ensuring that, in the absence of competition, all species have identical cumulative growth rates along the environmental gradient regardless 
    #of their niche breadth (i.e., same areas below the performance-environment curves, as in Büchi and Vuilleumier 2014).
    #10 is a scaling factor.
    
    Growth_rate<-(10*comp*habitat_suitability_each_spxsite) %*% diag((1/(Species_traits[,"nbreadth"]*sqrt(2*pi))))
    
    # Here Demographic stochasticity 
    Community_step2<-matrix(0,nrow(Community_step1),ncol(Community_step1))
    dimnames(Community_step2)<-dimnames(Community_step1)
    for(l in 1:nrow(Community_step2)){
      Community_step2[l,]<-rpois(ncol(Community_step2),round(Community_step1[l,]*Growth_rate[l,]))
    }
    
    
    ###Step2: Emigration in space and time. Here we demonstrate step by step emigration and immigration was done. 
    #It is not optimized intentionally to serve  as a easy-to-follow part of code.
    
    #Those that can potentially emigrate (Active individuals)
    Potential_Disperses<-which(Community_step2>0,arr.ind = T)
    Potential_Disperses<-cbind(Potential_Disperses,Species_traits[Potential_Disperses[,2],"Dispersal_ability"])
    colnames(Potential_Disperses)<-c("Site","Species","Dispersal ability")
    
    #Each species per site will send a number of disperses to the dispersal pool based on their dispersal ability
    success_dispersal<-cbind(Potential_Disperses,
                             rbinom(n=nrow(Potential_Disperses),size=Community_step2[Potential_Disperses[,1:2]],prob=Potential_Disperses[,3]),
                             Species_traits[Potential_Disperses[,2],"Dispersal_strategy"])
    colnames(success_dispersal)<-c("Site","Species","Dispersal ability","Number of Dispersers","Dispersal Strategy")
    
    
    #Among those that "decided to emigrate", let`s define how many will emigrate ("Dormancy column) in time based on Dispersal strategy (i.e., engage in dormancy)
    Dispersers_dormancy<-cbind(success_dispersal,rbinom(n=nrow(success_dispersal),size=success_dispersal[,"Number of Dispersers"],prob=success_dispersal[,"Dispersal Strategy"]))
    colnames(Dispersers_dormancy)<-c("Site","Species","Dispersal ability","Number of Dispersers","Dispersal Strategy","Dormancy")
    
    #Seeding the dormancy pool
    new_dormants<-matrix(0,N_sites,N_sp,
                         dimnames = list(c(sites_names),c(sp_names)))
    new_dormants[Dispersers_dormancy[,1:2]]<-Dispersers_dormancy[,"Dormancy"]
    dimnames(new_dormants)<-list(c(sites_names),c(sp_names))
    
    #Spatial Emigrants = Those that "decided"  to emigrate ("Number of Dispersers" Column) -  those that engaged in dormancy ("Dormancy" column)
    Emigration<-matrix(0,N_sites,N_sp,
                       dimnames = list(c(sites_names),c(sp_names)))
    
    Emigration[Dispersers_dormancy[,1:2]]<-Dispersers_dormancy[,"Number of Dispersers"]-Dispersers_dormancy[,"Dormancy"]
    dimnames(Emigration)<-list(c(sites_names),c(sp_names))
    
    
    Community_step3<-Community_step2-Emigration-new_dormants# 
    
    ###Step 3: Immigration 
    
    #Immigration in time
    # Individuals that engage in dormancy will recover from dormancy in the same site
    #but in a different moment (future) in time. We modeled an exponential decrease in the probability 
    #of recovering from dormancy. As such, individuals that, for instance, entered in dormancy in time t 
    #are more likely to recover from dormancy in a nearby future (e.g., t+1) 
    #than in distant moment in time (e.g., t+12). 
    
    #This part of the code can be optimized. Benchmark tests have shown that working with arrays considerably
    #speeds the process. I will fix that. The idea here is to populate the matrices in the "Dormancy_pool" list that represent
    #t> than current t.
    Dormancy_origns<-which(new_dormants>0,arr.ind = T)
    Dormant_ind<-cbind(Dormancy_origns,new_dormants[Dormancy_origns])
    colnames(Dormant_ind)<-c("Sites","Sp","Number of Dormants")
    
    if(sum(Dormant_ind)==0){
      Dormancy_pool<-Dormancy_pool
    }else{
      for(kk in 1:nrow(Dormant_ind)){
        Destination<-sample(t+1:11,Dormant_ind[kk,"Number of Dormants"],prob=Prob_recover_dormancy_over_time,replace=T)
        Abund<-table(Destination)
        for(s in 1:length(Abund)){
          Dormancy_pool[[as.numeric(names(Abund))[s]]][Dormant_ind[kk,"Sites"],Dormant_ind[kk,"Sp"]]<-Dormancy_pool[[as.numeric(names(Abund))[s]]][Dormant_ind[kk,"Sites"],Dormant_ind[kk,"Sp"]]+as.numeric(Abund[s])
          
        }
      }
      
    }
    
    #Immigration in space.
    #The sampling probability was given by the degree of connectivity between sites, 
    #meaning that patch j is more likely to receive spatial immigrants from closer 
    #patches than distant ones. 
    
    if(sum(Emigration)==0){
      imigration_final<-matrix(0,N_sites,N_sp,
                               dimnames = list(c(sites_names),c(sp_names)))    
    }else{
      #Non-Random colonization from the migration pool
      Origins<-which(Emigration>0,arr.ind =T)# Source site
      Imigrants<-cbind(Origins,Emigration[Origins])
      colnames(Imigrants)<-c("Sites","Sp","Number of Emigrants")
      imigration_final<-matrix(0,N_sites,N_sp,
                               dimnames = list(c(sites_names),c(sp_names)))  
      
      
      for(i in 1:nrow(Origins)){
        Imigrants_destination<-sample(sites_names,Imigrants[i,"Number of Emigrants"],Prob_dispersal_among_plots[,Imigrants[i,"Sites"]],replace = T)
        Abundance<-as.numeric(table(Imigrants_destination))
        imigration_final[names(table(Imigrants_destination)),Imigrants[i,"Sp"]]<-imigration_final[names(table(Imigrants_destination)),Imigrants[i,"Sp"]]+Abundance
      }
      
    }
    
    
    #Final communities
    Communities[[t]]<-Community_step3+imigration_final+Dormancy_pool[[t]]
    t=t+1
  }
  #### END OF SIMULATIONS#####  
  return(list(Communities,Species_traits))
}


#######Fixed parameters######
alpha_intra<-1/400# Intra-specific competition coeff.
alpha_inter<-1/800# Inter-specific competition coeff.
N_sp<-100#Initial species pool size
sp_names<-paste0("sp",seq(1,N_sp))
N_sites<-60# Number of patches considered
time=1200# Duration of simulation rouns
sites_names<-paste("Site",seq(1,N_sites),sep="_")


##### Simulations and Variation partitioning results#####

#Simulating Landscapes
#Generating a landscape that is asesaonal (Seasonlity_level=0.1) 
#and connectivity is low (Connectivity_level=0.9)
Connectivity_level=0.9# Values used in the manuscript were c(0.1,0.2,0.3,0.4,0.5,0.6,0.7,0.8,0.9)): The larger the value, the less physically connected is the landscape
Seasonlity_level=0.1# Values used in the simulations reported in the manuscript were c(0.1,0.5,1,1.7,3,7,20,30): The larger the value the more seasonal is the landscape in relation to spatial environmental heterogeneity

Land_tes<-Landscape_function4(N_sites = N_sites,Seasonality = Seasonlity_level,time = 1200,Spatial_Env_Grad = "Random",Connectivity =Connectivity_level ,plot_landscape =T ,phi=0.15)


# Simulating metacommunity dynamics on the landscape generated above
Metacomm_results<-Metacom_Function(Landscape_object=Land_tes,Dispersal_type="Equal")


#Only metacommunities in the last seasonal cycle were considered in further analyses.
Metacommunity_last_seasonal_cycle<-Metacomm_results[[1]][-1:-(time-12)]

#Assessing the species that remained in the metacommunity
Active_pool_composition<-as.data.frame(t(sapply(Metacommunity_last_seasonal_cycle,colSums)))
Active_pool_composition<-Active_pool_composition[which(rowSums(Active_pool_composition)>0),
                                                 which(colSums(Active_pool_composition)>0)]


#Seeing the traits of the remaining species
Species_traits_operational_pool<-Metacomm_results[[2]][colnames(Active_pool_composition),]

#Estimating relative abundance weighted  Niche breadth and dispersal ability
Weigthed_niche_breadth<-weighted.mean(Species_traits_operational_pool[,"nbreadth"],w=colSums(Active_pool_composition))
Weigthed_Dispersal_ability<-weighted.mean(Species_traits_operational_pool[,"Dispersal_ability"],w=colSums(Active_pool_composition))


#Using the simulated data to run  variation partitioning

#Creating the site-time-species matrix 
Big_community<-as.data.frame(do.call('rbind',Metacommunity_last_seasonal_cycle))
rownames(Big_community)<-paste0(rep(paste0("T",1:length(Metacommunity_last_seasonal_cycle)),each=N_sites),"_",rep(paste0("S",1:N_sites),length(Metacommunity_last_seasonal_cycle)))

#Removing empty rows and columns
Big_community<-Big_community[which(rowSums(Big_community)>0),which(colSums(Big_community)>0)]


# Extracting environmental variables
Env_pooled<-data.frame(Env=as.vector(Land_tes$Environment[,-1:-(time-12)]))
rownames(Env_pooled)<-paste0(rep(paste0("T",1:length(Metacommunity_last_seasonal_cycle)),each=N_sites),"_",rep(paste0("S",1:N_sites),length(Metacommunity_last_seasonal_cycle)))
Env_pooled<-subset(Env_pooled,rownames(Env_pooled)%in%rownames(Big_community))

# Extracting Spatial variables (Positive MEMs)
Spatial_MEM<-dbmem(data.frame(Land_tes$Coord),MEM.autocor = c("positive"))
Spatial_MEM_pooled<-do.call("rbind", rep(list(Spatial_MEM), length(Metacommunity_last_seasonal_cycle)))
rownames(Spatial_MEM_pooled)<-paste0(rep(paste0("T",1:length(Metacommunity_last_seasonal_cycle)),each=N_sites),"_",rep(paste0("S",1:N_sites),length(Metacommunity_last_seasonal_cycle)))
Spatial_MEM_pooled<-subset(Spatial_MEM_pooled,rownames(Spatial_MEM_pooled)%in%rownames(Big_community))

# Extracting coordinates variables (They are nescessary to run the GDMs)
Coords_pooled<-do.call("rbind", rep(list(Land_tes$Coord), length(Metacommunity_last_seasonal_cycle)))
rownames(Coords_pooled)<-paste0(rep(paste0("T",1:length(Metacommunity_last_seasonal_cycle)),each=N_sites),"_",rep(paste0("S",1:N_sites),length(Metacommunity_last_seasonal_cycle)))
Coords_pooled<-subset(Coords_pooled,rownames(Coords_pooled)%in%rownames(Big_community))

# Extracting Temporal variables (AEMs). All sites in a given time must have the same AEMs values
Temporal_AEM_pooled<-aem.time(length(Metacommunity_last_seasonal_cycle))$aem[rep(1:length(Metacommunity_last_seasonal_cycle), each=N_sites),]
dimnames(Temporal_AEM_pooled)<-list(paste0(rep(paste0("T",1:length(Metacommunity_last_seasonal_cycle)),each=N_sites),"_",rep(paste0("S",1:N_sites),length(Metacommunity_last_seasonal_cycle))),
                                    paste0("AEM",1:ncol(Temporal_AEM_pooled)))

Temporal_AEM_pooled<-subset(Temporal_AEM_pooled,rownames(Temporal_AEM_pooled)%in%rownames(Big_community))


#Estimating the components of the variation partitioning approach.They are relative importances,i.e.,
#portion of the total deviance explained that is attributable to each variable alone and their correlation (Joint)
New_Mean_RI_Spatial_beta_list_Per_landscape<-Var_part_GDMS4(Community = Big_community,Env_obs = Env_pooled,Coords = Coords_pooled,SpatialMEM=Spatial_MEM_pooled,TemporalAEM=Temporal_AEM_pooled)

#The components of variation partitioning are then ranked in ascending order of importance to facilitate the comparison across simulation iterations 
rank(New_Mean_RI_Spatial_beta_list_Per_landscape)

#In this study, the code above was used to run metacommunity dynamics across
# 20 replicates X  8 spatiotemporal heterogeneity levels (values described in line 593) x 9 connectivity levels (values described in line 592) 
#x 3 Spatial levels of spatial structure of env. conditions x 3 dispersal scenarios ("Equal", "Mainly Spatial", "Mainly Temporal")
```
